# Cryo-EM of ATP-driven dynamics and itraconazole binding of the fungal drug efflux ABC pump *Candida glabrata* Cdr1

**DOI:** 10.1101/2025.09.22.677697

**Authors:** Jorgaq Pata, Benjamin Wiseman, Eleftherios Zarkadas, Rim Baccouch, Nour Samrouth, Célia Desbois, Loïck Moissonier, Alexis Moreno, Atanu Banerjee, Stéphanie Aguerro, Emmanuel Bettler, Marie Dujardin, Sandrine Magnard, Raphaël Terreux, Guy Schoehn, Martin Högbom, Ahcène Boumendjel, Erwin Lamping, Richard D Cannon, Rajendra Prasad, Vincent Chaptal, Pierre Falson

## Abstract

Azole resistance in *Candida* species is often caused by the overexpression of Cdr1. Despite its clinical relevance, the structural basis for its ATP-driven efflux pump function remains elusive. We present four high-resolution cryo-EM structures for *Candida glabrata* Cdr1 under active turnover conditions in the absence and presence of ATP-Mg²⁺, itraconazole, and vanadate. Additional transient cryo-EM structures were unveiled by 3D variability analysis offering a detailed view of the step-by-step transitions triggered by ATP-hydrolysis. The motion cascade starts with a 4 Å piston-like retraction of the C-helix from the γ-phosphate/vanadate of the hydrolyzed ATP. This causes the nearby transmembrane helix-1 (TMH-1) to open the drug-binding site *via* lateral displacement and unwinding of the inner-leaflet region of TMH-2. A reverse ‘squeeze-and-push’ motion of TMH-2 possibly drives substrate extrusion. High resolution structures also reveal how itraconazole adapts its shape to fit into the drug-binding site. Our findings provide a dynamic structural framework for Cdr1-mediated azole resistance and the conserved chemo-mechanical cycle of ABC proteins, including non-membranous members.

## Introduction

Fungi infect more than 300 million people worldwide with an estimated 6.5 million people developing severe invasive infections, causing ∼2.5 million casualties each year (Denning, 2024). Fungi also infect plants and animals, causing the loss of one third of the global agricultural production (2017). Human infections are mainly caused by *Candida* species, opportunistic fungi that can invade tissues of immunocompromised hosts, such as those undergoing surgery or co-infected with other microorganisms (Hoenigl *et al*, 2022). Most of these infections are caused by the commensal yeast *Candida albicans*; however, other *Candida* species, such as *C. glabrata* (also known as *Nakaseomyces glabratus*) (Bolotin-Fukuhara & Fairhead, 2014) and *C. auris* (Du *et al*, 2020), are of an increasing concern due to their reduced or complete lack of sensitivity to antifungal therapies.

Fungicidal and fungistatic drugs have been developed to combat these infections (Drouhet & Dupont, 1987) by targeting yeast-specific biosynthetic pathways (Campoy & Adrio, 2017). Azoles are the most widely used antifungals, accounting for approximately 25% of the market (Jorgensen & Heick, 2021). These drugs bind to Cyp51 (Drouhet & Dupont, 1987; Monk *et al*, 2020), inhibiting the demethylation of ergosterol precursors—the lipid that constitutes 20% of yeast plasma membranes (van Meer *et al*, 2008).

As a first line of defense against azole antifungals, fungi overexpress plasma membrane efflux pumps that expel azole antifungals from the cytoplasm, preventing their action in the cytosol (Cannon *et al*, 1998; Prasad & Kapoor, 2005). In *Candida* species, the most prominent protein belongs to the ATP-binding cassette (ABC) superfamily (Decottignies & Goffeau, 1997; Gaur *et al*, 2005; Kumari *et al*, 2018), namely *Candida* drug resistance 1 (Cdr1). The protein shares 55% sequence identity with *Candida albicans* Cdr1 and 70% with the pleiotropic drug resistance 5 (Pdr5) transporter of *Saccharomyces cerevisiae* (Supplementary Figure 1A-C).

This resistance phenotype has typically been reported in patients for whom fluconazole-based therapy failed to eradicate *C. glabrata* infections (Ferrari *et al*, 2009). Figure 1A illustrates how the initially fluconazole-sensitive (EC_50_ ∼33 µM) clinical *C. glabrata* strain DSY562 responded to fluconazole exposure by overexpressing *Cg*Cdr1 through a mutation in the transcription factor Pdr1 (Ferrari *et al*., 2009), as depicted in Supplementary Figure 1D. The resulting strain DSY565 was insensitive to fluconazole, but remained sensitive to itraconazole, although at a 10-fold higher concentration than DSY562 (EC_50_ ∼223 nM *vs*. 27 nM). The upregulation of ABC efflux pumps also confers resistance to many other drugs due to their ability to bind and expel structurally diverse compounds, a phenotype identified as pleiotropic drug resistance (PDR) (Balzi & Goffeau, 1991).

**Figure 1.**
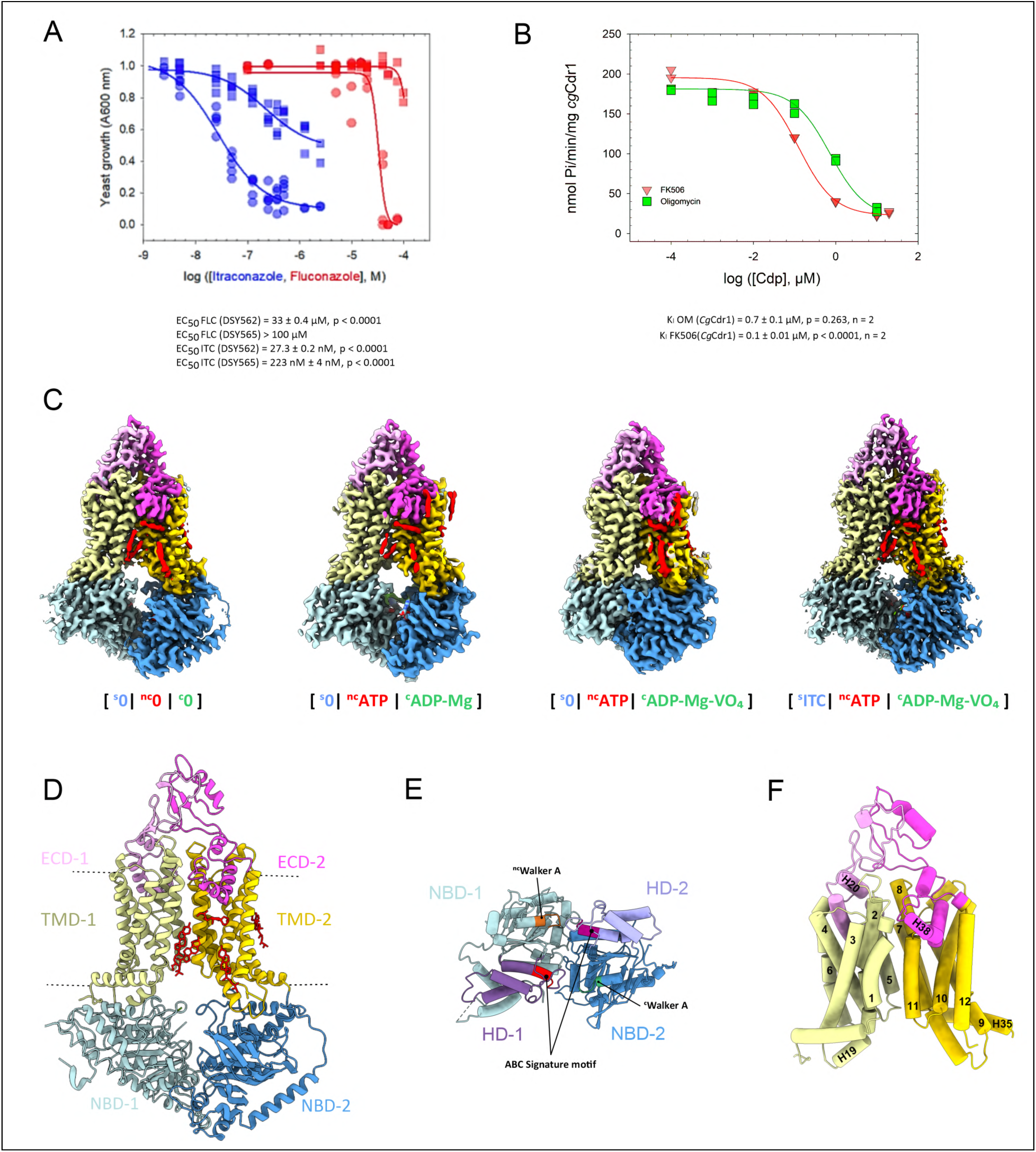
Phenotype and mechanism of azole antifungal resistance. **A.** Fluconazole and itraconazole resistance phenotypes of *C. glabrata* clinical isolates. Azole-sensitive (DSY562, circles) and azole-resistant (DSY565, overexpressing *Cg*Cdr1, squares) strains were exposed to increasing concentrations of fluconazole (red) or itraconazole (blue); *n* = 5. **B.** Inhibitor-sensitivity of the ATPase activity of purified *Cg*Cdr1. The methods and curve-fitting equations are described in Materials and Methods. **C.** Density maps of the four resolved states for *Cg*Cdr1, with the domains colored as indicated in panel D. **D.** *Cg*Cdr1 [^s^0|^nc^0|^c^0] cartoon model with the main domains distinguished by different colorings; ergosterol molecules are in red. **E.** Membrane-view of a cartoon model of the nucleotide-binding domains of *Cg*Cdr1 apo. The domains are colored as in Figure 1D. The helical domains HD-1 and HD-2 are depicted and colored in purple, lilac and the ABC signature motifs are colored in red and burgundy. **F.** Frontal cartoon view of the transmembrane domains, numbered and named according to the sequence presentation of *Cg*Cdr1 in Supplementary Figure 1C. The domains are colored as in panel D.

About one third of all yeast ABC transporters are PDR transporters (Decottignies & Goffeau, 1997; Gaur *et al*., 2005). PDR transporters are ∼1,500-residue long polypeptides organized as pseudo-dimers, with two transmembrane domains (TMDs), each composed of six transmembrane helices (TMHs), and two cytoplasmic nucleotide-binding domains (NBDs), whose head-to-tail dimerization forms two nucleotide-binding sites (NBS) (Supplementary Figure 1E). Efflux pump substrates bind to a drug-binding site (DBS) centrally located at the interface between the two TMDs and are then expelled into the extracellular space through a mechanism driven by ATP binding and hydrolysis at the NBSs. In PDR exporters, these domains form two NBD-TMD tandems with a characteristic type V subfamily topology (Thomas *et al*, 2020); for a review see (Banerjee *et al*, 2021; Banerjee *et al*, 2023; Moreno *et al*, 2019). The first NBS (NBS-1) consists of the Walker A and B motifs of NBD-1 and the ABC signature motif, also referred to as the C-loop, of NBD-2. NBS-1 binds but does not hydrolyze ATP due to substitutions of key catalytic residues. It is referred to hereafter as ^nc^NBS for “non-catalytic nucleotide-binding site”. The second NBS (NBS-2) consists reciprocally of the Walker A and B motifs of NBD-2 and the ABC signature motif of NBD-1. It binds and hydrolyzes ATP and is referred to hereafter as the catalytic NBS, ^c^NBS (Supplementary Figure 1E).

Structural information, both static and dynamic, is still lacking in explaining the evolutionary adaptation that rendered NBS-1 non-catalytic. However, our team and others have highlighted its critical importance both in *Sc*Pdr5 (Downes *et al*, 2013; Sauna *et al*, 2008) and in *Ca*Cdr1 (Banerjee *et al*, 2020; Banerjee *et al*, 2018). In each case, the drug-susceptibility phenotype of strains with mutations at the periphery of the outer leaflet region of the TMD was reported to be reverted by secondary site mutations located in the ^nc^NBS, far from the primary mutation. Notably, one of these secondary site mutations involved the glutamine residue of the ABC signature motif – Q1005 in *Ca*Cdr1 and Q1002 in *Cg*Cdr1 – suggesting a close functional relationship between these regions, the structural basis of which remains unknown; for a review see (Banerjee *et al*., 2021).

Recently, the cryo-EM structures of functional Pdr5 reconstituted in peptidiscs provided the first molecular-level insights into the organization of PDR exporters (Harris *et al*, 2021), followed by the structure of Cdr1 from *C. albicans* in complex with fluconazole and milbemycin oxime, but with questionably low ATPase activity (Peng *et al*, 2024). Indeed, obtaining structural information on *Candida* PDR transporters has been more challenging, primarily due to difficulties in preserving the functional integrity of these proteins once extracted from their native membrane environment.

Here, we describe several high-resolution cryo-EM structures of purified *C. glabrata Cg*Cdr1 with functionally assessed ATPase activity, in the apo form and in complex with itraconazole, nucleotides, and vanadate. The high quality of the datasets enabled us to explore their structural variability, providing an experimentally verified, detailed description of the conformational dynamics following ATP hydrolysis and the alternating access mechanism of the pump. These structures also reveal how the membrane domain of *Cg*Cdr1 accommodates itraconazole within its drug-binding site (DBS) and how the substrate adapts a n-shaped conformation to perfectly fit into the DBS.

### Functional detergent-purified *Cg*Cdr1

*Cg*Cdr1 was overexpressed in a *S. cerevisiae* membrane protein expression system, conferring azole resistance to the hyper-sensitive yeast host (Lamping *et al*, 2007) (Supplementary Figure 2A). Most of *Cg*Cdr1 was enriched in the 20,000x g membrane fraction (C20K). Fifty percent of the ATPase activity in this fraction corresponded to approximately 150–200 nmol Pi·min⁻¹·mg⁻¹. This ATPase activity was sensitive to nanomolar concentrations of oligomycin (Decottignies *et al*, 1994) or FK506 (Kralli & Yamamoto, 1996), two well-established PDR pump inhibitors (Supplementary Figure 2B). Their effects were not additive, as no further inhibition was observed when both compounds were added simultaneously (Supplementary Figure 2B), confirming that each molecule specifically targets *Cg*Cdr1. The inhibition constants (*K*i) of oligomycin and FK506 were estimated to be 110 and 20 nM, respectively (Supplementary Figure 2C). The oligomycin/FK506-sensitive ATPase activity of 150–200 nmol·min⁻¹·mg⁻¹ was therefore attributable to *Cg*Cdr1. Assuming that 4.4% of the CK20 membrane fraction was accounted for by *Cg*Cdr1 (Supplementary Figure 2A), the specific ATPase activity of *Cg*Cdr1 was estimated to be 3-5 µmol·min⁻¹·mg⁻¹ (⬄ 1 ATP hydrolyzed/*Cg*Cdr1/100 ms), consistent with previous reports for Pdr5 (2 µmol·min⁻¹·mg⁻¹) (Wagner *et al*, 2019). Measuring the ATPase activity in the presence of increasing concentrations of itraconazole resulted in a ∼30% reduction of the *Cg*Cdr1-specific ATPase activity, with half-maximal effect in the micromolar range (Supplementary Figure 2D). As observed for FK506, inhibition of the ATPase activity by itraconazole was also not additive in combination with 20 µM oligomycin (Supplementary Figure 2D).

*Cg*Cdr1 was subsequently extracted and purified using a mixture of trans-PCC-α-M (PCC) (Hovers *et al*, 2011) and dicarboxylate oside 9b (DCOD9b) (Comsa *et al*, 2018), as previously described (Pata *et al*, 2023) (Supplementary Figure 2E). The oligomycin-sensitive ATPase activity of *Cg*Cdr1 (200 nmol·min⁻¹·mg⁻¹) in this detergent mixture was comparable to the oligomycin-sensitive ATPase activity of the CK20 membrane fraction with a *K*_I_ of 0.7 and 0.1 µM for oligomycin and FK506, respectively (Figure 1B). All these biochemical characterizations confirmed that *Cg*Cdr1 remained functional in the PCC-DCOD9b environment, and that it was amenable to structural studies.

### High resolution cryo-EM structures of *Cg*Cdr1

*Cg*Cdr1 particles of the purified and ligand-free form, representing the apo state were imaged. The protein was also imaged under active turnover conditions by incubating the purified sample with 5 mM ATP-Mg²⁺ for 30 min at 4 °C, followed by the addition of 1 mM orthovanadate for a further 20 min. Vanadate, in its inorganic form Vi, replaces the inorganic phosphate (Pi) generated by ATP hydrolysis, forming a stable ADP-VO₄²⁻-Mg²⁺ complex that mimics the ADP-PO₄²⁻-Mg²⁺ complex (Harris *et al*., 2021; Loo & Clarke, 2002). In addition, *Cg*Cdr1 was incubated under the same conditions but including 30 µM itraconazole (Figure 1C, Supplementary Figures 3-6). The loading status of the different binding sites in *Cg*Cdr1 was denoted as [^s^X|^nc^X|^c^X], where *s* represents the substrate bound to the DBS, and *nc* and *c* indicate nucleotides bound to the ^nc^NBS and ^c^NBS, respectively.

The structure of *Cg*Cdr1 closely resembles that of *Sc*Pdr5 (Harris *et al*., 2021) and *Ca*Cdr1 (Peng *et al*., 2024) with a RMSD of 1.4 Å and 1.8 Å, respectively across all Cα, confirming a general fold for fungal PDR transporters. The protein displays a pseudo dimeric organization, with each half containing a NBD, a TMD, and one extracellular domain (ECD) (Figure 2C-F). In the nucleotide-bound structures, a linker domain is observed near the ^nc^NBS (green in Figure 2A). This domain comprises a short *N*-terminal segment (residues 124–151) and the stretch connecting TMH-6 to NBD-2 (residues 788–847). Initially identified in Pdr5, this structural feature is a hallmark of PDR transporters and mainly observed in closed-NBD forms. In the apo-state structure, neither of the linker domains was formed due to the absence of nucleotides which prevented domain dimerization (Figure 1E). The TMDs are organized into six-helix bundles that provide a pathway for substrates into the extracellular space (Figure 1F). TMH-5 and TMH-11 connect to two helical regions, bent at a 90° angle that form two short amphipathic helices (H-20 and H-38 in Supplementary Figure 1C) at the extracellular water-membrane bilayer interface. Between these helices, and TMH-6 and TMH-12 are two ECDs, much larger than in human type-V ABC transporters. As in Pdr5, the *N*-terminal and *C*-terminal halves of these two ECDs were stabilized by one and two disulfide bonds, respectively.

**Figure 2.**
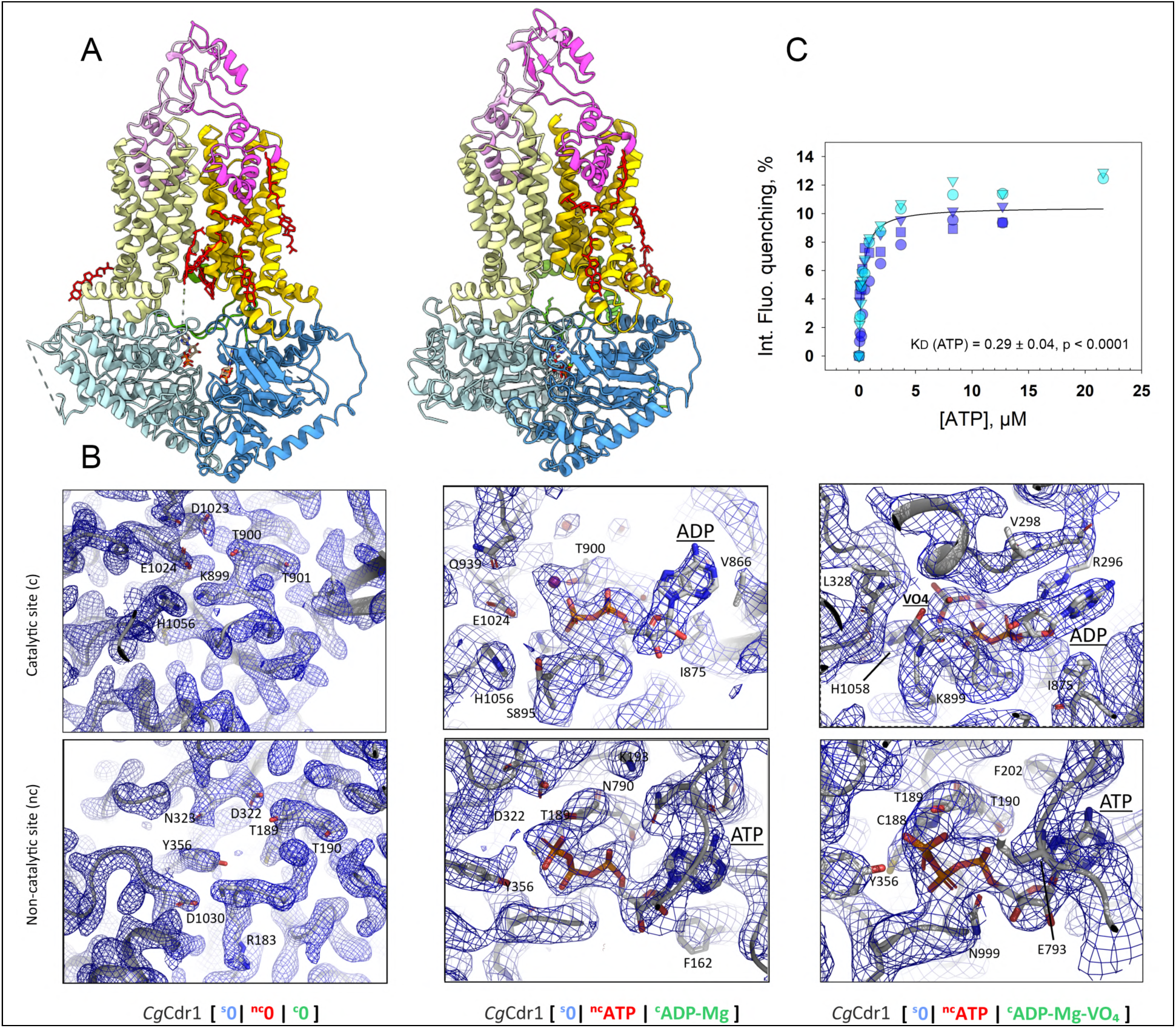
Models, cryo-EM map details, and ATP binding of *Cg*Cdr1. **A.** *Left:* Cartoon model of *Cg*Cdr1 [^s^0|^nc^ATP|^c^ADP-Mg] with nucleotides and tightly bound ergosterol molecules shown as sticks. *Right*: Cartoon model of *Cg*Cdr1 [^s^0|^nc^ATP|^c^ADP-Vi-Mg] with nucleotides and tightly bound ergosterol molecules shown as sticks. The domains are colored as in Figure 1D. **B.** Density maps of the nucleotide binding sites of *Cg*Cdr1 in different states, shown as mesh, at a contour level of 0.1 for [^s^0|^nc^0|^c^0], 0.3 for [^s^0|^nc^ATP|^c^ADP-Mg] and 0.52 for [^s^0|^nc^ATP|^c^ADP-Vi-Mg]. The nucleotides and the main nucleotide-interacting *Cg*Cdr1 residues are shown as sticks. **C.** Binding of ATP to *Cg*Cdr1 probed by intrinsic fluorescence quenching performed in two independent experiments of technical triplicates.

### ^nc^ATP and ^c^ADP bound to the ^nc^NBS and ^c^NBS of *Cg*Cdr1

The two maps that were obtained for the open and the closed conformations of *Cg*Cdr1 from the same purified sample incubated with ATP-Mg^2+^ and vanadate (see Supplementary Figure 6 for further details) both had an ATP molecule without magnesium bound to the ^nc^NBS (Figure 2B middle and right panels), confirming the previous observations in Pdr5 (Harris *et al*., 2021). The persistence of an ATP molecule bound to the ^nc^NBS in both the open and the closed conformations of *Cg*Cdr1 prompted us to evaluate its affinity with an intrinsic fluorescence quenching assay. Indeed, the data yielded saturation curves that fit well with a single ATP binding site (without Mg^2+^), with a *K*D of 0.3 µM (Figure 2C). The apo state structure of *Cg*Cdr1 (Figure 2B left panel) lacked electron densities for nucleotides bound to NBS. Inversely, the ^c^NBS of the open (Figure 2B middle panels) and the closed (Figure 2B right panels) conformations of *Cg*Cdr1 were bound to either an ADP-Mg^2+^ or an ADP-Vi-Mg^2+^ complex. These findings suggest that the release of vanadate had already occurred in a subset of the purified *Cg*Cdr1 particles while vanadate had locked the other particle population in a post-hydrolytic state resembling the [^nc^ATP|^c^ADP-Pi-Mg] state before the release of the γ-phosphate and the return of the transporter to its inward-facing conformation.

The two NBSs displayed markedly different ATP affinities, reflecting their asymmetric nature. ATP bound the ^nc^NBS with very high affinity (*K*D of 0.3 µM), substantially stronger than its affinity previously determined for the ^c^NBS of its *Ca*Cdr1 ortholog (≈150 µM) (Pata *et al*., 2023). Thus, at millimolar ATP concentrations typically found inside living yeast cells (Takaine *et al*, 2019), ATP is expected to remain predominantly bound to the ^nc^NBS of *Cg*Cdr1, supporting a role for ATP binding at this site as a structural and functional cofactor throughout the transport cycle of highly asymmetric fungal PDR transporters.

### Cryo-EM dynamics of ATP-driven motion

#### Capturing the motion of ATP-driven opening of the transporter

The ability to capture two closely related yet distinct conformational states of *Cg*Cdr1 from a single sample incubated with nucleotides and vanadate on a single grid, as illustrated in Figures 1–2, provided a unique opportunity to leverage the full potential of inter-particle variability for a more detailed analysis of the cryo-EM samples (Punjani & Fleet, 2021). Cryo-EM data processing, including 2D and 3D classifications, identified three distinct classes of particles. The two most abundant classes, which comprised 50% and 32% of the particles, corresponded to the closed [^s^0|^nc^ATP|^c^ADP-Vi-Mg] and to the open [^s^0|^nc^ATP|^c^ADP-Mg] conformations, respectively. The third class of particles represented an intermediate conformation that could not be resolved at high resolution (Supplementary Figure 6C).

We aligned the particles of all three classes with the most abundant one and performed a variability analysis starting from the closed [^s^0|^nc^ATP|^c^ADP-Vi-Mg] structure. The latent coordinate analysis revealed a non-Gaussian particle distribution (Supplementary Figure 7A), indicative of the presence of sub-states (Supplementary Figure 7B). Crucially, the continuum between these sub-states suggested that the selected particles should enable the capture of transitional conformations. Variability analysis was conducted using 3DVA in CryoSPARC (Punjani & Fleet, 2021), which generated twenty density maps ordered along a continuous and sequential conformational pathway. This 3DVA was performed for both primary components C0 and C1, the 2 main components representing the largest movements of *Cg*Cdr1 that could be deciphered from the dataset. The resulting density maps were further refined into 3D models using the previously developed phenix.varref tool (Afonine *et al*, 2023) (Supplementary Figure 7C, Supplementary Table 2) that allowed us to calculate the RMSD for each Cα atom of each frame relative to frame 1 (Supplementary data 1 and 2 for C0 and C1, respectively). Although the ordering of conformations derived from 3DVA does not represent any true kinetic information, the continuity and constrained nature of the observed transitions define a mechanically coherent pathway linking ATP hydrolysis to domain rearrangements.

#### Global and local 1D displacement analyses

The resulting 3D models (representing 20 different poses of *Cg*Cdr1 after ATP-hydrolysis) for component 0 are displayed superposed in Figure 3A and the animated version of this time-resolved movement is shown in Supplementary movie 1. We found component C0 captured large-scale conformational transitions, involving all domains of *Cg*Cdr1.

**Figure 3.**
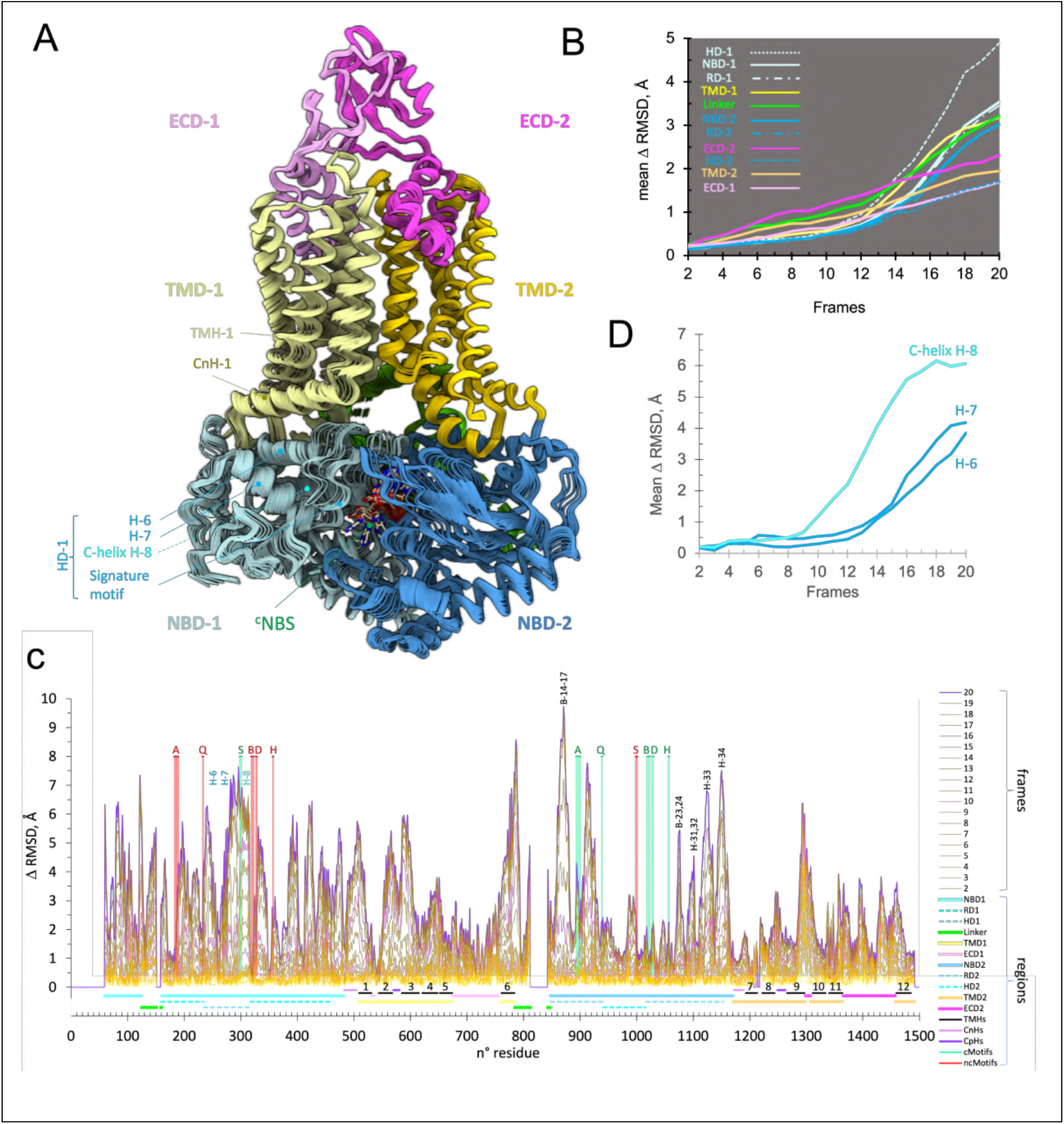
Conformational variability analysis of component C0 of the transition from the closed [^s^0|^nc^ATP|^c^ADP-Vi-Mg] to the open [^s^0|^nc^ATP|^c^ADP-Mg] conformation of *Cg*Cdr1. **A.** Superposition of the 3D models of the 20 frames obtained by 3DVA. The image shows the projected movements viewed from the catalytic side of the transporter. **B.** Plots of the mean of the Cα RMSD of frames 2 to 20 *vs* frame 1 for each of the indicated regions. Regions are colored as in panel A. the ‘helical’ and ‘RecA’ sub-domains, HD and RD, of each NBD are also included. **C.** Plots of Cα RMSD per residue for frames 2 to 20 relative to frame 1. The traces from 2 to 19 are colored with a gradient from yellow to brown; traces 10, 15, and 20 are colored with a gradient from pink to purple. The position of the different domains is indicated below the graph and the position of the various catalytic and non-catalytic motifs, colored in green and red, respectively, are displayed directly on the graph. A, B: Walker A, B, S: ABC signature motif, Q, D, and H: Q, D, and H loops (see detailed locations in Supplementary Figure 1C). **D.** Plots of the Cα RMSDs of frames 2 to 20 *vs* frame 1 of all H-6 (V243-K254), H-7 (R265-T279) and H-8 (G300-I313) residues.

At the domain level (Figure 3B), analysis of the RMSD variation revealed two groups of domain rearrangements based on their amplitude of motion: one with large displacements of 3-5 Å, including the NBDs, the linker, and TMD-1, and another with smaller displacements of up to 2 Å, comprising TMD-2 and the two ECDs. For the NBDs, consisting of the RecA-like (RD) and the helical (HD) subdomains (*49-51*) (see detailed locations in Supplementary Figure 1C), the analysis revealed a particularly large displacement (∼5 Å) of HD-1, a critical region for ATP-binding and hydrolysis at the ^c^NBS.

At the primary sequence level (Figure 3C), analysis of the RMSD variation showed that the conserved NBD motif residues—green for catalytic, red for non-catalytic—remained largely in low-mobility regions, consistent with their nucleotide-binding role. However, the ABC signature motif of NBD-1 and the Walker A motif of NBD-2, part of the ^c^NBS, displayed significantly larger displacements of 4 to 6.5 Å, highlighting their critically important contribution to ATP hydrolysis. As indicated in Figure 3C, several NBD regions were particularly important for the conformational changes of *Cg*Cdr1 triggered by ATP-hydrolysis. These were most notably the C-helix (H-8) that is connected to the C-loop of the ABC signature motif in HD-1 and several regions in RD-2, notably B-14 to B-17 and B-23-B-24 β-sheets, and H-31 to H-34 α-helices.

A closer inspection of the 3 α-helices H-6, H-7, and H-8 of HD-1 showed that they experienced distinct mobilities and timing, despite belonging to the same helical subdomain (Figure 3D). H-6 and H-7 exhibited similar mobilities (Figure 3D) with an up to 4 Å displacement of individual residues, displacements that were much more pronounced for residues closer to the ^c^NBS (Supplementary Figure 8A). However, H-8 experienced the largest displacement of 6 Å (Figure 3D) but, unlike H-6 and H-7 residues that experienced quite distinct mobilities across the entire helix, all H-8 residues retracted from the ^c^NBS to the same extent (Supplementary Figure 8A). Notably, the movement of H-8 started as early as in frame 8, preceding that of H-6 and H-7, which only started to move after frame 12. This sequence indicates that the retraction of H-8 from the catalytic center represents the earliest structurally detectable conformational change of *Cg*Cdr1, preceding the motion of other helical elements within the same subdomain and subsequent rearrangements of the transporter.

The relative displacements calculated for individual TMH residues confirm the generally higher mobility of TMD-1 helices than those of TMD-2 (Supplementary Figure 8, panels B *vs*. C). TMH-1, -2, -3, and -6 (panel B) undergo, in addition, an asymmetric deformation with larger displacements observed for their inner leaflet segments than their outer leaflet segments. The displacement of these TMD-1 helices was tightly coordinated with the H-6 and H-7 motions starting at frame 12, highlighting the retraction of H-8 as the earliest motion triggered by ATP-hydrolysis at the ^c^NBS.

The same analysis of C1 component (Supplementary Figure 9, Supplementary movie 2) showed lower, yet still significant, mobility of the protein, capping the mobility at a maximum of 2 Å (panel B). However, structural models gathered for this component revealed a particularly significant motion for TMH-2 and -5 with higher displacement values experienced by segments embedded in the inner leaflet of the bilayer that shift by up to 4–5 Å (Supplementary Figure 10).

#### Global and local 3D displacement analyses

Supplementary movie 1 reveals how the opening of *Cg*Cdr1 triggered by ATP hydrolysis is rooted along a vertical axis normal to the membrane plane passing through TMH-5 and the ATP molecule permanently bound to the ^nc^NBS right underneath. The movements of important regions in terms of angles and distances are detailed in Figure 4. They are anchored in TMD-2, which acts as a pivot, with an overall vibration of no more than 2.3 Å (panel A). The ^nc^NBS, positioned directly underneath TMD-2, also barely moves (Figure 4A, lower right panel). However, the catalytic Walker A and Walker B residues of NBD-2 that bind to the ADP-Vi-Mg^2+^ complex retract from NBD-1 by ∼4-5 Å (Figure 4A, lower right panel). The transition from the closed to the open conformations of NBD-1 caused a ∼10° rotation of the entire domain centered on the β-sheet B-8 residue T352 (Figure 4A, lower left panel). This rotation translated to TMD-1 with the same amplitude and centered at TMH-5 that is in contact with TMD-2 and doesn’t move much at all (Figure 4A, upper left panel). The rotation is visible for all TMHs and depicted for TMH-1 and TMH-3 (Figure 4A, upper left panel). TMH-2, however, experienced unique movements by partially unfolding the lower half of the helix (F558-I573), as described above (Figure 4A, upper left panel).

**Figure 4.**
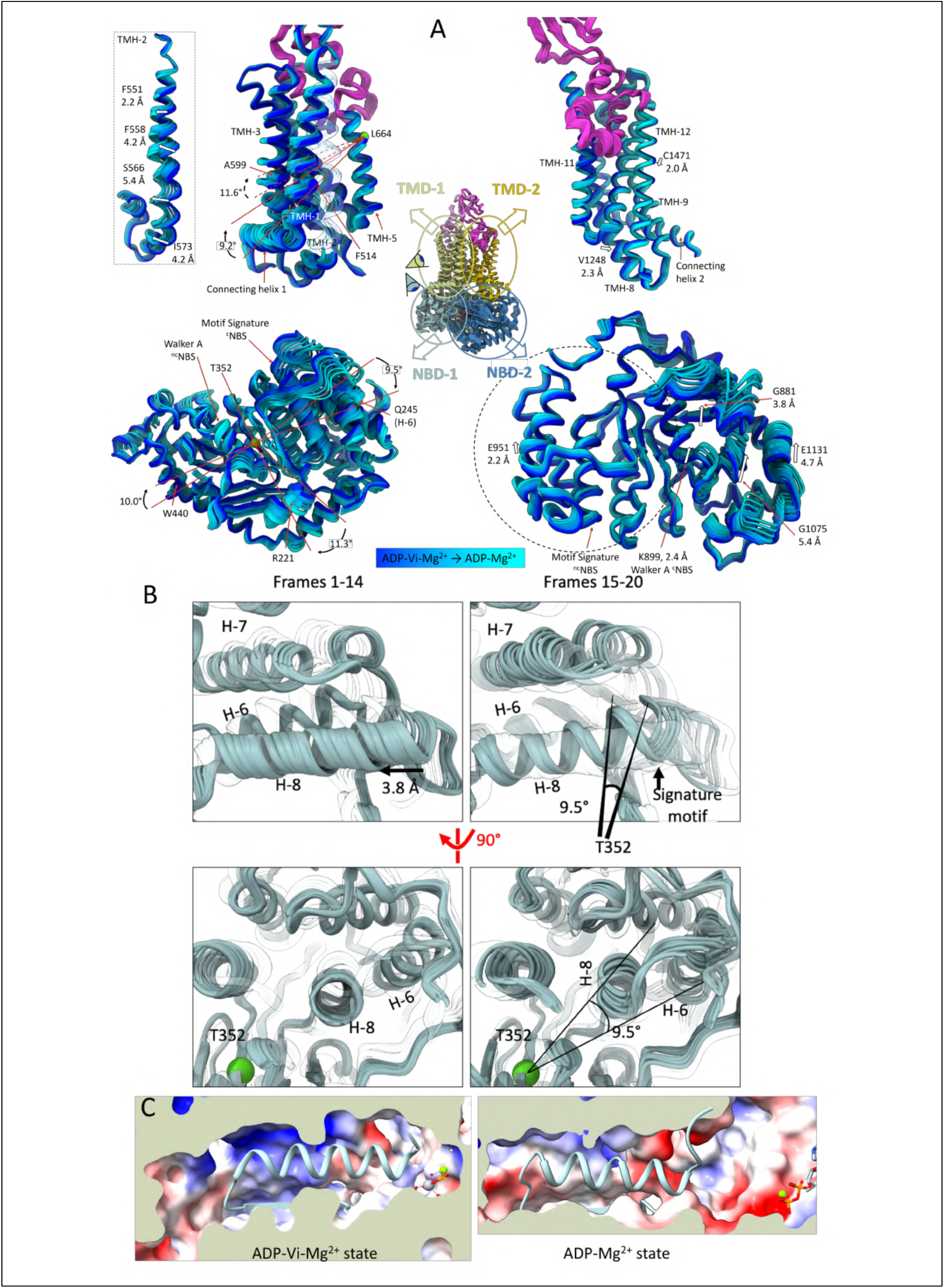
Three-dimensional transition from the closed [^s^0|^nc^ATP|^c^ADP-Vi-Mg] to the open [^s^0|^nc^ATP|^c^ADP-Mg] conformation of *Cg*Cdr1 triggered by a piston-like motion of the C-helix H-8. **A.** Projected 3D-motions of the NBD and TMD regions of component C0 particles of *Cg*Cdr1. The 20 different structures (i.e. frames) are shown as cartoons colored from blue to cyan depicting the dynamic transition from the closed to the open conformation of *Cg*Cdr1. TMH-2 is transparent in the TMD-1 image and presented separately right next to it, because its movement is different from the rest of TMD-1. Movements are listed for each domain with white arrows indicating directionality and amplitude. **B.** HD-1 subdomain and C-helix H-8 piston-like motions. Cartoon model of HD-1 colored in light green as NBD-1 in Figure 2. On the left side frames 1-14 are in solid color while the rest is transparent and on the right-side frames 1-14 are transparent and frames 15-20 are presented in solid color. The bottom panel shows the same representation turned through 90°. Helices are labeled on the image for clarity. Frames 1-14 show a piston-like motion of the C-helix in the plane of the helix, while frames 15-20 show a further displacement of the C-helix out of the plane that is a rotation of 9.5° relative to the fixed T352 position. **C.** Slice view of the C-helix shown as a cartoon with the electrostatic surface potential shown for the surrounding region. In the closed ADP-Vi-Mg^2+^ state, the *N*-terminus of the C-helix coordinates the γ-phosphate leaving a cavity at its *C*-terminus. In the open ADP-Mg^2+^ state, the C-helix has retracted into this cavity, filling up the void.

#### The C-helix H-8 triggers large conformational changes of CgCdr1 following ATP hydrolysis

A key feature highlighted by the variability analysis described above is the dramatic and rigid retraction of the C-helix H-8 from the ^c^NBS. In the [^s^0|^nc^ATP|^c^ADP-Vi-Mg] state, the *N*-terminal region of H-8 established H-bonds between the vanadate and residues G300 and G301 that are part of the conserved VSGG ABC signature motif. ATP-hydrolysis at the ^c^NBS appears to trigger a rigid 3.8-Å retraction of the H-8 helix from the vanadate towards the back of the domain in a piston-like motion (Figure 4B, Frames 1-14). Throughout this movement, there was minimal movement observed anywhere else in the protein. This initial movement was followed by a second type of movement shown in Figure 4B (frames 15-20) whereby the C-helix experienced a 9.5° rotation centered at T352 that matched the ∼10° rotation of the rest of the NBD-1 and parts of TMD-1, as mentioned above. Helix H-8 thus undergoes two distinct motions while moving away from the ADP-Vi-Mg^2+^ complex in a translation intimately linked to ATP hydrolysis. This piston-like movement is enabled by a cavity at the *C*-terminus of the helix (Figure 4C left panel) that is only present in the closed conformation but not in the inward-facing open conformation of *Cg*Cdr1 (Figure 4C left *vs* right panels). This initial piston-like motion precedes the domain-scale rearrangements following ATP hydrolysis.

#### Local constrictions of the TMDs squeeze the DBS

Contrary to the global movements revealed by the 3DVA of C0 component (Figure 4A left, Supplementary movie 1), the 3DVA of C1 component revealed a constriction-like movement of certain TMHs of TMD-1 towards the center of the DBS (Supplementary Figure 11, Supplementary movie 2), while *Cg*Cdr1 remained in the closed [^s^0|^nc^ATP|^c^ADP-Vi-Mg] vanadate-bound state during all stages of this constriction. But even in this conformation, no exit pathway was visible.

3DVA revealed that TMH-3, -4 and -6 appeared to collapse towards TMH-2 and -5 while the coupling helix CpH-1 between TMH-2 and -3 moved upwards and stretched between residues F565 and A570, transmitting a spring-like force to the rest of TMH-2, which compressed and moved TMH-2 upwards. A similar upward motion was observed for TMH-5. In TMD-2, the central part of TMH-8 (F1230-S1246) collapsed into the DBS, with TMH-9 and -10 moving as a rigid body and pivoting towards the center of the TMDs (Supplementary Figure 11). While the absence of the substrate and no visible exit pathway does not allow us to unambiguously attribute this movement to a certain stage of the transport cycle, we speculate that this ‘squeeze and push’ movement may also play a crucial role in substrate translocation, forcing the substrate through a hydrophobic exit valve (Alhumaidi *et al*, 2022) above the DBS and pushing the substrate through the ECD into the cell exterior.

### Ergosterol molecules bind to distinct *Cg*Cdr1/membrane interface sites

A common feature across these reconstructions was the presence of multiple densities into which ergosterol molecules fit well (colored in red in Figure 1C). We also quantified by MS/MS 11 ergosterol molecules and the equivalent of one phosphatidylethanolamine (PE)-type and one phosphatidylcholine (PC)-type lipid per *Cg*Cdr1 molecule in the purified sample (Supplementary Figure 12). Densities of the latter lipids could not be unambiguously assigned to the density maps, but those of ten ergosterol molecules could be resolved and were labeled Erg-1 to Erg-10 (Supplementary Figure 13A).

The distribution of the ergosterol molecules was asymmetric, with Erg-1 to Erg-6 binding to the catalytic face of the protein (Figure 5, Supplementary Figure 13B). Erg-1 bound to a cavity capped by hydrophobic residues F1370, W1371, F1373, M1374, and V1377 of helix H-38 in ECD-2 and was stabilized laterally by Y1325 and L1329 of TMH-10, M1350 and L1353 of TMH-11, and Y1472 and F1475 of TMH-12 (Supplementary Figure 13AB). Erg-2 bound near the cytoplasmic side of TMD-2, alongside TMH-10. Erg-3 was located within a groove between TMH-9, -10 and -12, in the center of the membrane bilayer. Erg-4 and Erg-5 were also positioned near the center of the membrane bilayer, between TMH-1 and TMH-11, facing the entrance to the DBS (Supplementary Figure 13B).

**Figure 5.**
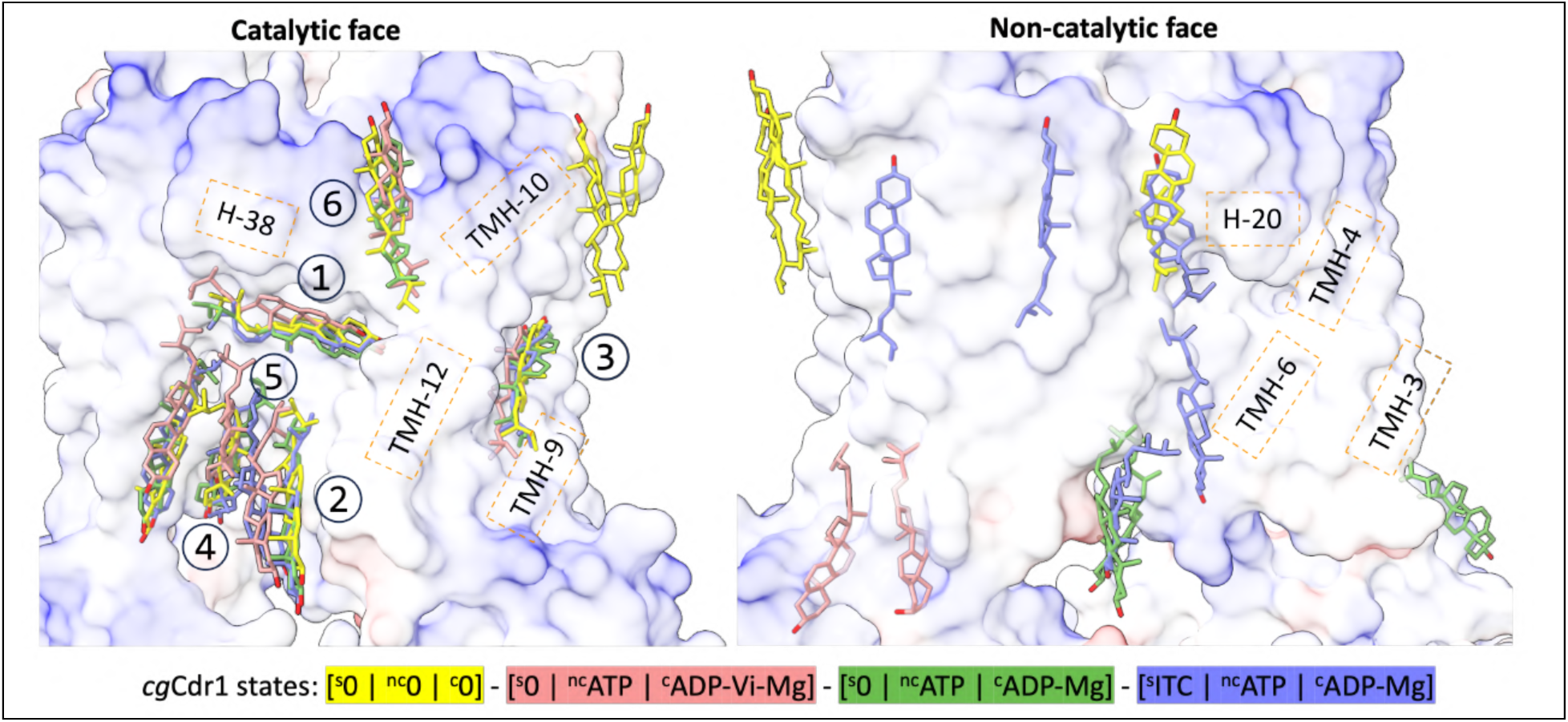
Location of endogenous ergosterol molecules bound to the TMD of *Cg*Cdr1. The TMD regions of the four states of *Cg*Cdr1 are superposed, with views of the catalytic (left) and non-catalytic (right) sides of the TMD. Ergosterol molecules are numbered and color-coded according to the state of *Cg*Cdr1 indicated at the bottom, with the apo state surface displayed as a reference.

It is noteworthy that *Ca*Cdr1 has been demonstrated to transport steroid hormones (Baghel *et al*, 2017). Several of the residues mentioned above were previously identified as key residues involved in steroid hormone transport by *Ca*Cdr1, specifically interacting with β-estradiol and corticosterone (Baghel *et al*., 2017; Krishnamurthy *et al*, 1998) (Supplementary Figure 13C). For instance, the Y1328A mutation in *Ca*Cdr1 (Y1325 in *Cg*Cdr1) altered β-estradiol transport, while the M1332A (L1329 in *Cg*Cdr1) and M1356A (L1353 in *Cg*Cdr1) mutations reduced corticosterone transport, consistent with our previous findings that the majority of residues critical for steroid hormone transport were located in TMH-10 and -11 (Baghel *et al*., 2017). As with Erg-1, several other ergosterol-interacting sites have been implicated in β-estradiol and/or corticosterone transport. Notably, L663 in *Ca*Cdr1 which corresponds to the Erg-10 interacting residue L662 in *Cg*Cdr1, is essential for the transport of both β-estradiol and corticosterone (Baghel *et al*., 2017).

The mobility of these ergosterol molecules was further evaluated by molecular dynamics simulations of *Cg*Cdr1 placed into a lipid bilayer. This analysis revealed reduced mobility of the ergosterol molecules (Supplementary Figure 13D) suggesting an important role of these molecules for the structural integrity of the pump. The reason for the asymmetric distribution of ergosterol molecules is unclear and warrants further investigation. One plausible hypothesis is that this distribution reinforces the rigidity of TMD-2, creating a structural anchor for TMD-1. An alternative hypothesis is that *Cg*Cdr1 facilitates the translocation of ergosterol between the inner leaflet (Erg-2) and the outer leaflet (Erg-6) through an intermediate location (Erg-1) reminiscent of the “credit card swipe mechanism” described for the transport of phosphatidylcholine-type lipids by human ABCB4 (Prescher *et al*, 2021).

### Itraconazole folds into the drug-binding site and stabilizes the inward-facing conformation

The affinity of the purified protein for itraconazole was estimated at a *K*_D_ of 0.4 µM (Figure 6A). As in the C20K membrane environment (Supplementary Figure 2C), itraconazole induced a limited yet significant reduction of *Cg*Cdr1 ATPase activity with a half maximal effect at a concentration of 3.5 µM but still plateauing at 75% beyond 10 µM (Figure 6B). This limited effect may reflect a slower translocation of itraconazole through the DBS due to its rather large substrate size, as also previously reported (Ernst *et al*, 2008; Golin *et al*, 2003). As expected from its binding mode, only one itraconazole molecule fit into the DBS, occupying a space from near the inner leaflet of the membrane bilayer to the upper half of the TMHs that constitute the DBS (Figure 6C-D).

**Figure 6.**
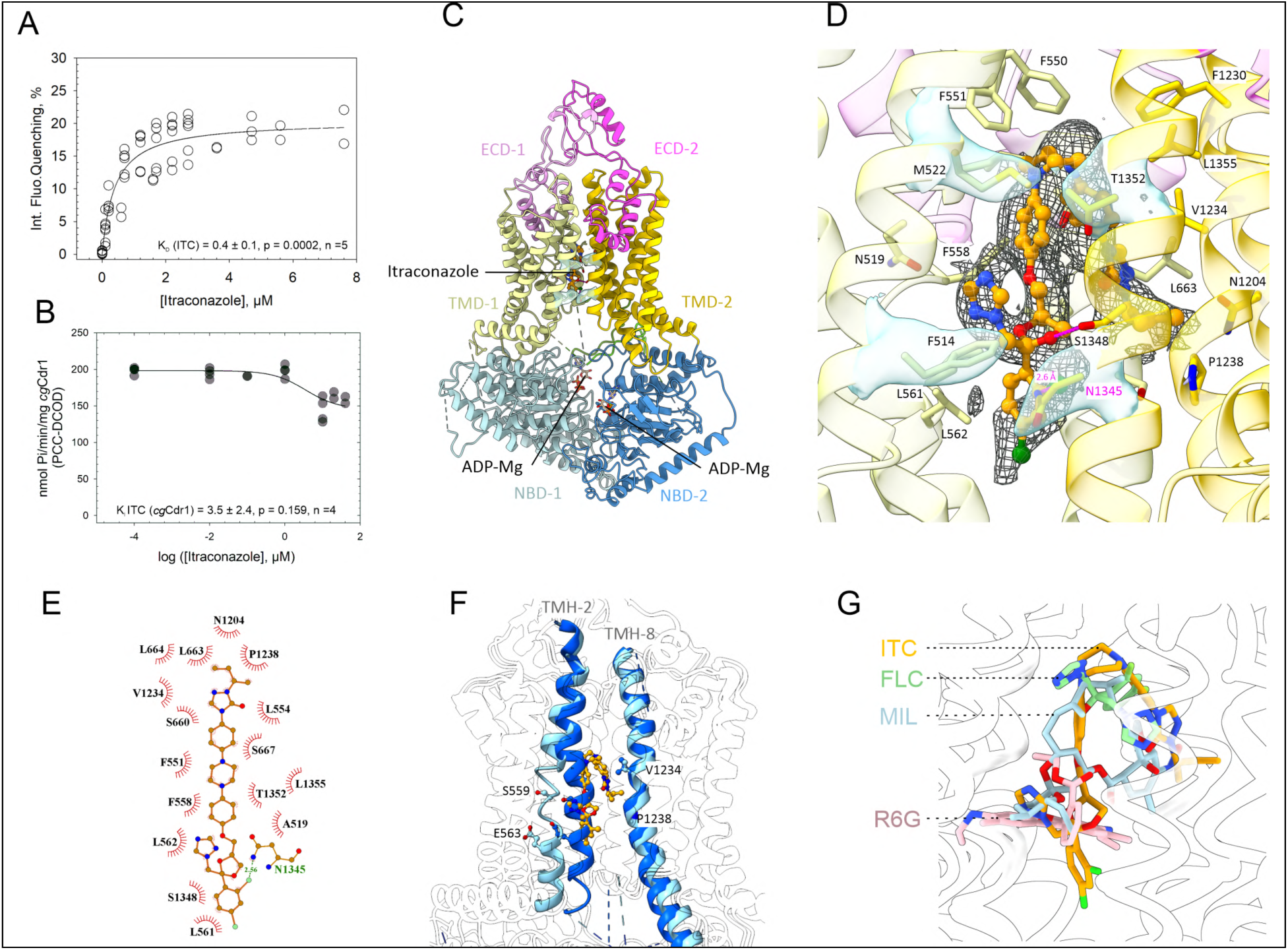
Binding of itraconazole to the DBS. **A.** Itraconazole binding to *Cg*Cdr1 measured by intrinsic fluorescence quenching**. B.** ATPase activity of purified *Cg*Cdr1 in the presence of increasing concentrations of itraconazole. **C.** Structure of *Cg*Cdr1 [^s^ITC|^nc^ATP|^c^ADP-Mg], domains colored as in Figure 1B, and itraconazole and nucleotides shown as sticks. The densities of F514, M522, N1345, and T1352 delineating the entry path are transparent blue. **D.** Enlarged itraconazole binding region. Itraconazole is shown as sticks and balls colored by atom type, with carbon in orange. The density map is displayed as a mesh at a contour level of 0.0225. Potential H-bonds with the indicated distances between itraconazole and residues of the DBS are depicted in magenta dashed lines. Residues establishing Van der Waals contacts are labeled and shown as sticks. The densities of F514, M522, N1345, and T1352 delineating the entry path are shown in transparent blue. **E.** 2D plot of the interactions established by itraconazole and the residues of the DBS. Itraconazole is drawn in orange, with its nitrogen atoms in blue, oxygen in red, fluorine in green. The labelled residues bordered with a red semi-circle establish Van der Waals interactions, while N1345 establishes an H-bond, as indicated in green. Plot generated by Ligplot. **F.** Local deformations of TMH-2 and -8 in the [^s^0|^nc^ATP|^c^ADP-Vi-Mg] and the [^s^ITC|^nc^ATP|^c^ADP-Mg] states are colored in dark and light blue, respectively. Boundary residues of the deformations are shown as sticks: S559 and E563 for TMH-2 and V1234 and P1238 for TMH-8. Itraconazole is shown in orange sticks as a reference. **G.** Superposition of *Cg*Cdr1 [^s^ITC|^nc^ATP|^c^ADP-Mg], with ITC in orange, *Ca*Cdr1 [^s^Fluconazole] (green, RMSD across 1297 pairs of Cα = 1.8 Å), *Ca*Cdr1 [^s^milbemycin oxime] (light blue, RMSD across 1,295 pairs of Cα = 1.9 Å) and *Sc*Pdr5 [^s^Rhodamine 6G] (pink, RMSD across 1,314 pairs of Cα = 1.3 Å). The proteins are shown as transparent white cartoons.

Binding of itraconazole to the DBS of *Cg*Cdr1 in the presence of ATP-Mg^2+^ froze the transporter in the inward-facing conformation after ATP-hydrolysis and the release of Pi (Figures 1C, 6C and Supplementary Figure 14). Further refinements identified a density in the crevice between the TMDs into which itraconazole could be fitted (Figure 6D). Two analyses were performed with the two separate ITC datasets, leading us to model ITC as the molecule binding to this DBS, even though the electron densities for itraconazole were of limited quality. The presence or absence of vanadate during the pre-incubation period of *Cg*Cdr1 with ATP-Mg^2+^ and itraconazole led to the same nucleotide loading state, with densities for ADP-Mg^2+^ in the ^c^NBS and ATP in the ^nc^NBS clearly visible (Supplementary Figure 14B).

The itraconazole structures in the absence or presence of vanadate were almost identical to the [^s^0|^nc^ATP|^c^ADP-Mg] state with an overall RMSD of 0.5 Å, indicating that itraconazole binding does not induce major conformational changes.

Itraconazole adopted an n-shaped conformation with the triazolone ring positioned near the bottom of the DBS. The bulkier dioxolane, triazole, and dichlorophenyl moieties were arranged in a triskelion-like fashion, oriented towards the entry pathway. Binding was facilitated by residues of TMH-1, -2, -5 -8 and -11. The chlorine atom of the dichlorophenyl group formed a halogen bond with the nitrogen atom of N1345. Residue S1348 was close enough to the oxygen atom of the dioxolane to establish an H-bond that may be critical for drug binding. This agrees with previous mutation studies of the equivalent residues S1360 in Pdr5 and T1351 in *Ca*Cdr1 that affected drug binding and efflux pump inhibition by FK506 (Egner *et al*, 2000; Egner *et al*, 1998; Kueppers *et al*, 2013; Nim *et al*, 2016).

Itraconazole primarily interacted with the protein through hydrophobic and Van der Waals interactions (Figure 6E) as reflected by the DBS’s composition of mostly aliphatic and aromatic residues: F551, L554, L562, I564, and F565 of TMH-2; L663 and M656 of TMH-5; V1234 and P1238 of TMH-8; and L1355 of TMH-11. The itraconazole binding pocket also featured hydrophilic regions at its periphery. Compared to the ligand-free inward-facing structures, there was no significant rearrangement of the side chains of these residues. Residues F550 and F551 of TMH-2 and F1230 of TMH-8 function as a hydrophobic lid, preventing itraconazole from advancing further toward the ECD by positioning themselves directly above the piperazine moiety. On the opposite side, the entry pathway was delineated by the lower regions of TMH-1 and -11, facilitating substrate entry from the membrane. However, in all inward-facing structures, this pathway was obstructed by F514 and N1345 at the bottom, as well as M522 and T1352 in the upper part of the entry route, with their Cryo-EM densities clearly defined (Figure 6D). Perhaps these residues function as a drug entry filter. This hypothesis was strengthened by the spatial vicinity (one helical turn upwards of F514) of a glycine residue, G518. The equivalent residue in *Ca*Cdr1 (G521) was previously identified critical for substrate gating, most likely because it provides the required conformational flexibility to the entry gate (Niimi *et al*, 2022).

During model building, there were noticeable local deformations of TMH-2 and to a lesser extent TMH-8, compared to the [^s^0|^nc^ATP|^c^ADP-Vi-Mg] state. Figure 6F shows the unwinding of two helical turns of TMH-2 between residues 559-565 due to a loss of the backbone H-bond between S559 and L562. This unwinding created space for itraconazole binding and conferred different flexibilities to the two halves of the helix: its lower half was positioned parallel to itraconazole, while the upper part leaned over the substrate, closing the exit pathway in this inward-facing conformation. TMH-8 also showed local deformations of residues V1234 to P1238, which generated additional space for itraconazole to bind (Figure 6F). TMH-2 and -8 thus appear to be the key structural elements for accommodating substrates within the DBS. This adaptability of the DBS is also complemented by the ability of ITC to fold over itself, thus optimizing its spatial occupancy within the DBS.

Densities for ITC were not as clear as nearby amino-acid residues indicating either flexibility of the ligand in the binding-pocket, several orientations of the ligand in the DBS or low occupancy (Supplementary Figure 15). This phenomenon was also observed in *Ca*Cdr1 and *Sc*Pdr5 maps. We think that these multidrug transporters, which have to accommodate multiple chemically unrelated ligands, possess a rather loose binding pocket, with the substrates residing there instead of being tightly bound, which may impede a well-defined density in cryoEM maps. Importantly, the location of itraconazole overlaps quite well with the positions of rhodamine 6G in Pdr5 (Harris *et al*., 2021) and with fluconazole and milbemycin oxime in *Ca*Cdr1 (Peng *et al*., 2024) (Figure 6G).

## Discussion - Model for the transport cycle of *Cg*Cdr1

The present *Cg*Cdr1 structures and the 3DVAs enabled us to visualize sequential steps of the large conformational changes experienced by the transporter during ATP-hydrolysis and the release of Pi. They allow us to propose a model for the transport cycle of *Cg*Cdr1, as depicted in Figure 7. In the absence of ATP-Mg at the ^c^NBS, the protein adopts an inward-facing, open, conformation represented by the [^s^0|^nc^0|^c^0] and the [^s^0|^nc^ATP|^c^ADP-Mg] structures. At this stage of the transport cycle the DBS is open towards the cytoplasm and accessible to substrates. This state was adopted by the [^s^ITC|^nc^ATP|^c^ADP-Mg] structure, which suggests that i) substrate-binding does not induce any major conformational changes of *Cg*Cdr1 and ii) substrates simply reside in, rather than tightly bind to, the DBS, which may partially explain the polyspecificity of the protein. Binding of ATP at the ^c^NBS switches the transporter into the closed conformation, with the TMHs collapsing into the DBS and forcing substrate transport into the extracellular space by a ‘squeeze and push’ movement, similar to the movements depicted for the component C1 particles of the 3DVA analysis of the [^s^0|^nc^ATP|^c^ADP-Mg] bound state. ATP hydrolysis at the ^c^NBS is tightly associated with the reopening of the transporter toward the inward-facing conformation. This dynamic was captured for the first time by carefully analyzing the structural continuum of the process. The Component 0 of the 3DVA describes this movement in detail, identifying the ABC signature motif and its downstream C-helix (H-8) of NBD-1 as triggers for opening the transporter. The present data also revealed that ATP remains bound to the ^nc^NBS throughout the entire transport cycle, with the site locked in place by the linker domain, acting as a pivot for domain rotation. This is further supported by its high affinity (*K_D_* ≈ 0.3 µM) for ATP, showing that the two NBSs have distinct roles for the substrate transport across cell membranes.

**Figure 7.**
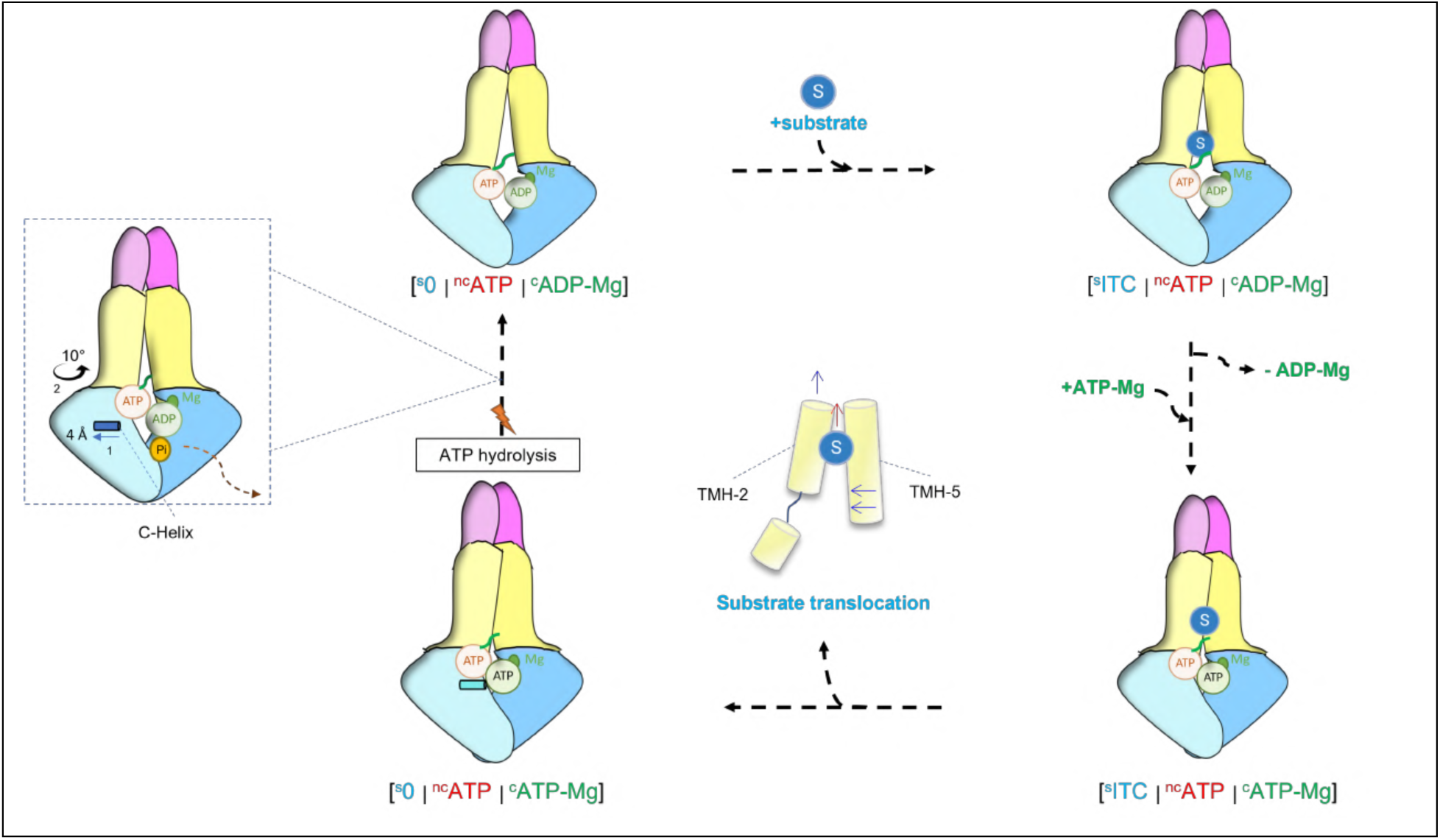
Proposed transport model for *Cg*Cdr1. NBD-1 and -2 are depicted in light blue and blue, TMD-1 and -2 in pale yellow and yellow, ECD-1 and -2 in rose and purple, respectively, and the linker domain is in green. The substrate translocation step possibly involves a ‘squeeze and push’ movement of the centrally located TMH-2 and -5 that is similar to their movement described for component C1 particles of the [^s^0|^nc^ATP|^c^ADP-Vi-Mg] dataset. ATP hydrolysis triggers the backward movement of the C-helix that dissociates the ^c^NBS.

## Material & Methods

### Materials

Bacto-yeast extract and bacto-peptone were purchased from Difco Laboratories, Detroit, MI, USA.

Luria Bertani (LB) and Yeast-Peptone-Dextrose (YPD) media were purchased from Carl Roth Gmbh & Co. Kg (Karlsruhe, Germany). Trans-PCC-α-M (Hovers *et al*., 2011) (PCC) was purchased from Glycon Biochemicals, Luckenwalde, Germany. Dicarboxylate oside detergent 9b (Nguyen *et al*, 2018) (DCOD 9b) was a gift from CALIXAR, Lyon, France. EDTA-free antiprotease mix was purchased from Roche SAS, Boulogne-Billancourt, France. HiFliQ Nickel-NTA columns were purchased from Generon, Slough, UK. Superdex 200 column was purchased from CYTIVA Europe Gmbh, Velizy-Villacoublay, France. Amicon®ultra-centrifugal filters were purchased from Merck KGaA, Darmstadt, Germany. Oligomycin, FK506 and itraconazole were purchased from Sigma, L’lsle-d’Abeau Chesnes, France, and prepared as stock solutions in 100% DMSO.

### Strains

Fluconazole-sensitive *Candida glabrata* DSY562 and fluconazole-resistant *Candida glabrata* DSY565 (Ferrari *et al*., 2009) were kindly provided by Pr. Dominique Sanglard (Lausanne, Switzerland).

The yeast strain AD/CgCdr1-His_6_ used in this study for producing *Cg*Cdr1 was derived from the *Saccharomyces cerevisiæ* AD1-8u^-^ strain in which genes coding for the ABC exporters Yor1, Snq2, Pdr5, Pdr10, Pdr11, Ycf1, Pdr15, and the transcription factor Pdr3 were deleted, rendering the strain hypersensitive to drugs (Decottignies *et al*, 1998). *Cg*C*DR1* encodes the protein sequence referenced as Q6FK23 in the Uniprot protein database, from the *C. glabrata* laboratory strain CBS 138, and was cloned with a C-terminal hexa-histidine tag integrated at the *PDR5* locus of AD1-8u^-^ to form AD/CgCdr1-His_6_ (Lamping *et al*., 2007). AD1-8u^-^ and AD/*Cg*Cdr1-His_6_ were a gift from Richard Cannon and Erwin Lamping.

### Methods

#### Drug susceptibility assays

The clinical strains *C. glabrata* DSY562 and DSY565 were revived on solid YPD medium for 48 h and then grown in liquid YPD at 30 °C overnight for three consecutive days before being used for drug susceptibility assays. On the final day, the culture was diluted in the morning to reach an OD_600-nm_ of 0.5 by the evening. The culture was diluted 1/1000 and 198 µL aliquots were distributed in duplicate into wells of a 96-well microtiter plate, and 2 µL of increasing concentrations of antifungals dissolved in DMSO were added to each well. The microtiter plate was incubated at 30 °C while shaking at 200 rpm overnight. The assay was stopped when the OD_600-nm_ of the wells without antifungal reached 0.9 to 1.1 OD_600-nm_ values after correction for the background absorbance of the medium without yeast, and OD_600-nm_ values analyzed using SigmaPlot v12.5 with the equation f = min + (max-min)/(1+10^(logEC_50_-x)).

#### Cdr1 expression and purification

Cultures, membrane preparation and Cdr1 purification were performed as previously described (Pata *et al*., 2023). Briefly, Cdr1 extraction from a membrane fraction pelleted at 20,000 x g (C20K) was performed by mixing a the membrane fraction suspended in Suspension Buffer (SB: 50 mM Tris-acetate pH 7.5, 1 mM EDTA, 20% glycerol, 1 mM PMSF, EDTA-free antiprotease) at a protein concentration of 5 g/L with 10 g/L PCC and 1 g/L DCOD 9b for 1 h at 4 °C with gentle agitation. The insoluble material was removed by ultracentrifugation at 100,000 x g for 1 h at 4 °C and the detergent-extracted supernatant was loaded onto a 5-mL HiFliQ Nickel-NTA column equilibrated with SB. The resin was washed with 50 mM HEPES-HCl, pH 7.5, 150 mM NaCl, 70 µM (2x CMC) PCC ± 4 µM DCOD, 20 mM imidazole and *Cg*Cdr1 was eluted by increasing the imidazole concentration to 150 mM. The pool of about 10 mL was 20x-concentrated using Amicon®ultra-centrifugal filters with a cutoff of 100 kDa at 1,500 x g to limit detergent over-accumulation (Gobet *et al*, 2023) and then loaded on a Superose 200 10/300 column with a running buffer containing 50 mM HEPES-HCl, pH 7.5, 100 mM NaCl, 70 µM (2x CMC) PCC, 4 µM DCOD. The peak eluting at 15.5 mL corresponding to *Cg*Cdr1 was collected, pooled and either stored at 4° C or frozen in liquid nitrogen and stored at -80 °C.

#### Ligand binding assays

Binding of itraconazole and ATP was probed by intrinsic fluorescence quenching on a SAFAS Xenius spectrophotofluorimeter (SAFAS, Monaco). Tryptophan residues and N-acetyl tryptophan amide (NATA, used as negative control) were excited at 290 nm with a slot of 5 nm and their fluorescence emission spectra were recorded between 310 and 380 nm with a slot of 5 nm for ATP or between 310 and 340 nm with a slot of 3 nm for itraconazole. Experiments were performed in a final volume of 200 µL in a quartz cuvette, to which increasing amounts of compounds were added, at a controlled temperature of 25°C and incubation times of 5 min. The raw quenching data were subtracted from the quenching induced by the DMSO in which itraconazole was dissolved. Note that the total amount of added DMSO was always maintained below 0.7%. NATA, suspended in the same buffer as *Cg*Cdr1, was used to correct nonspecific effects. Fluorescence emission curves were integrated between 310 and 380 nm, using the equation f = 100*ABS((F/F0)_Cdr1_-(F/F0)_NATA_), where F is the fluorescence emitted by *Cg*Cdr1 or NATA at a given ligand concentration and F0 being the fluorescence emitted before ligand addition. Ligand-binding models were applied to the datasets using Graphpad Prism 9, choosing the model displaying the lowest Akaike index criterion (AIC). Final plots were generated with SigmaPlot V12.5, using the equation for ligand binding of one site saturation, f = *Bmax*\*abs(x)/(*K*D + abs(x)).

#### ATPase activity

ATP hydrolysis was measured by quantifying the released inorganic phosphate (Pi)(Drueckes *et al*, 1995). *Cg*Cdr1-associated ATPase activity was assessed by suspending 10 µg/10 µL of the C20K membrane fraction or 4 µg *Cg*Cdr1/10 µL in a final volume of 100 µL of buffer containing (final concentrations) 8 mM MgCl₂, 5 mM ATP, and 60 mM Tris-HCl (pH 7.5), along with 1 mM NaN₃ and 5 mM KNO₃ as F₁F₀-ATPase and pyrophosphatase inhibitors (Centeno *et al*, 1994). Oligomycin (Decottignies *et al*., 1994), FK506 (Kralli & Yamamoto, 1996; Maesaki *et al*, 1998) or itraconazole were added to the reaction mixture at concentrations ranging from nM to µM, as indicated. After 30 min incubation at 30 °C, the reaction was stopped by adding 100 µL of a freshly prepared cold solution containing (final concentrations) 10 g/L SDS, 10 g/L ammonium molybdate, 2 g/L ascorbic acid, and 40 g/L H₂SO₄. The amount of released Pi was quantified by measuring the absorbance at 880 nm using a SAFAS Xenius plate reader. A standard curve was generated using known amounts of KH₂PO₄ (10–100 nmol) prepared under the same conditions in the absence of C20K or *Cg*Cdr1. Data were analyzed using SigmaPlot v12.5 with the equation f = min + (max-min)/(1+10^(logEC_50_-x)).

#### Lipid quantification

Internal standards (^13^C cholesterol, phosphatidylcholine (PC 14:1/17:0), and phosphatidylethanolamine (PE 14:1/17:0)) were added to the sample, and lipids were extracted twice using a chloroform/ethanol (2:1, v/v) mixture. The samples were centrifuged at 2,000 rpm for 5 min, and the organic phase collected and evaporated to dryness under a flow of nitrogen gas. For ergosterol analysis, the samples were derivatized with N,O-bis(trimethylsilyl)trifluoroacetamide (BSTFA) and analyzed by gas chromatography (GC) (HP 7890B, Agilent) coupled with triple quadrupole mass spectrometry (GC-MS/MS) using electron impact ionization (EI) mode (7000 C, Agilent). The GC-MS/MS system was equipped with a SolGel-1ms fused silica capillary column (Trajan, SGE; 60 m × 0.25 mm). Helium was used as the carrier gas at a flow rate of 1.2 mL/min. The split/splitless injector temperature was set to 280 °C. The oven temperature program was as follows: the initial temperature was set at 55 °C for 4 min, followed by an increase of 40 °C/min up to 250 °C, then 20 °C/min up to 310 °C, where it was held for 25 min. Samples were injected in splitless mode. The transfer line temperature of the mass spectrometer was set to 250 °C, and the ion source temperature was 200 °C. Nitrogen was used as the collision gas at a flow rate of 1.5 mL/min. The ionization energy was set to 70 eV. Ergosterol was detected using the multiple reaction monitoring (MRM) mode. Peak detection, integration, and quantification were performed using MassHunter software.

Phospholipid analyses were performed using an ultra-high-performance liquid chromatography tandem mass spectrometry (UHPLC-MS/MS) system consisting of a Shimadzu Nexera LC-30 instrument (Noisiel, Marne-la-Vallée, France) equipped with a Waters ACQUITY UPLC BEH Amide column (2.1 × 100 mm, 1.7 µm). Separation was achieved under gradient elution conditions: mobile phase A consisted of acetonitrile/water containing 10 mM ammonium formate (95:5, v/v), while mobile phase B consisted of acetonitrile/water containing 10 mM ammonium formate (50:50, v/v). Gradient elution started at 1 % B with a flow rate of 0.6 mL/min, increased to 20% B at 2 min, then to 80 % B at 5 min, before returning to the initial conditions. The column oven temperature was set to 45 °C, and 5 µL of the sample was injected. The UHPLC system was coupled to a linear ion trap quadrupole mass spectrometer (Qtrap 4500, Sciex, Les Ulis, France) equipped with an electrospray ionization (ESI) source. The instrument was operated with a spray voltage of -4,500 V, a source temperature of 500 °C, a curtain gas setting of 40, and optimized collision nitrogen gas. Negative ionization mode using scheduled multiple-reaction monitoring (MRM) was applied for phospholipid identification and quantification. Peak detection, integration, and quantification were performed using Analyst and Sciex OS software (Sciex).

#### Sample preparation for cryo-EM

Purified *Cg*Cdr1 was thawed and concentrated to 6-7 g/L using Amicon®ultra-centrifugal filters with a cutoff of 100 kDa at 1,500 x g. The sample (3 µL) was either applied onto a grid (apo state) or pre-incubated with 5 mM ATP-Mg^2+^ and with or without 30 µM itraconazole for 30 min at 4 °C followed by addition of 1 mM orthovanadate for 30 min prior to applying it to the grid.

#### Grid preparation and data acquisition

Apo and itraconazole-ATPMg-vanadate samples (Stockholm): Motion correction was performed with PatchMotion, and contrast transfer function (CTF) parameters were estimated from averaged movies using PatchCTF in cryoSPARC v4. To remove any form of bias, automatic particle selection was performed using a blob picker without the use of templates. The number of particle images was reduced by 2D and 3D classifications (Supplementary Figure 3, Supplementary Figure 4). Initial maps were calculated ab initio and the final maps were refined using cryoSPARC’s non-uniform refinement feature without any applied symmetry. Detailed image processing parameters can be found in the Supplementary Table 1. 3D variable analysis was performed using particles from consensus refinements, using a filter resolution of 4.5 Å and solving for 2 components.

#### ATPMg-vanadate samples (Grenoble)

UltraAufoil Au/Au 1.2/1.3 grids (Quantifoil) were glow discharged on air for 50 s at 30 mA (Emitech Glow Discharge). A volume of 3.5 µL of the mix was applied on freshly glow discharged grids at 20 °C and 100 % humidity using a Vitrobot Mark IV (Thermo Fisher Scientific). Excess liquid was blotted for 4 s at blot force 0 and 0.5 s drain time before vitrification in liquid ethane. Grids were verified with small data collection on a Thermo Scientific Glacios Cryo-TEM at 200 kV equipped with a Gatan K2 summit detector (IBS, Grenoble) prior to high-end data acquisition on a Thermo Scientific Titan Krios at 300 kV equipped with a Gatan K3 direct electron detector (ESRF CM01). Image acquisition parameters and analysis statistics are summarized in Supplementary Table 1.

#### Cryo-EM image processing

Apo and itraconazole-ATPMg-vanadate samples (Stockholm): Motion correction was performed with PatchMotion, and contrast transfer function (CTF) parameters were estimated from averaged movies using PatchCTF in cryoSPARC v4 (Structura Biotechnology Inc, Toronto). To remove any form of bias, automatic particle selection was performed using a blob picker without the use of templates. The number of particle images was reduced by 2D and 3D classifications (Supplementary Figure 3, Supplementary Figure 4). Initial maps were calculated *ab initio* and the final maps were refined using cryoSPARC’s non-uniform refinement feature without any applied symmetry. Detailed image processing parameters can be found in Supplementary Table 1. 3D variable analysis was performed using particles from consensus refinements, using a filter resolution of 4.5 Å and solving for 2 components.

*ATPMg-vanadate samples (Grenoble):* Data were processed using cryoSPARC v4. Movies were submitted to patch motion correction and patch CTF estimation jobs. Particle picking was performed externally using CrYOLO (Sphire) with the general model trained for low pass filtered images, and the resultant locations were imported to cryoSPARC. Particles were extracted at 384 pixels, Fourier-cropped at 192 pixels box size and curated with 2 rounds of mild 2D classification. All class averages displaying a single well centered particle with clear features of a micelle-embedded ABC transporter were selected for the following steps. 3D initial models were created with a 3-class *ab initio* reference-free reconstruction followed by heterogeneous refinement. All particles were refined against the best-looking volume with a homogeneous non-uniform refinement. The resultant particles were then re-extracted re-centered at full box size (384 pixels) and subjected to final round of 2D classification, excluding only featureless class averages. The selected particles were used either in a consensus homogeneous non-uniform refinement job or to generate 3D initial models in a 3-class *ab initio* reference-free reconstruction job followed by a heterogeneous refinement against the 3 resulted initial models. Two of those classes generated high-resolution reconstructions displaying two distinct conformations of ADP-Mg- and ADP-Mg-vanadate bound to the ^c^NBS of *Cg*Cdr1. The particles from each of these two classes were refined separately in homogeneous and non-uniform refinement jobs to generate conformation 1 and conformation 2, respectively (see Supplementary Figure 6). Particles from each of these two refinements, as well as, from the previous consensus refinement job, were subjected to 3DVA after filtering at 4-Å resolution and applying a soft mask generated by the refinement mask dilated by 20 Å.

#### 3D-model building and refinement

The apo-structure of Pdr5 (PDB 7P03) was docked into a 2.7 Å cryo-EM density map and improved with iterative rounds of manual building in Coot and Isolde followed by real_space_refine in Phenix. The final model was thus built using both sharpened and unsharpened maps. The final model was validated with MolProbity and EMringer and deposited in the Protein Data Bank under the accession code 6R81 and in the electron microscopy database EMDB-4749. Water molecules were added using the Douse_Phenix and their presence, geometry and distances were confirmed by manual inspection followed by relaxation in Isolde. The figures were constructed using Pymol 2.8 (Schroedinger, Inc. NY, USA) and UCSF ChimeraX (University of California, San Francisco) (Meng *et al*, 2023). PDB and EMDB codes and model statistics are provided in Supplementary Table 1.

#### Molecular dynamics simulations

Molecular dynamics simulations of *Cg*Cdr1 parameters were derived from the Generalized Amber Force Field version 2 (GAFF2), using Antechamber software. Partial atomic charges for ADP, ATP and Pi were computed using AM1-BCC semi-empirical quantum mechanical calculations. The [^s^0|^nc^ATP|^c^ADP-Pi-Mg] state was built with ADP, Pi, and Mg^2+^ being placed into the nucleotide binding domain (NBD) of *Cg*Cdr1 at their binding site by superposing the outward-facing conformation of *Cg*Cdr1. Following protein-lipid system building, charge neutrality was ensured by randomly removing the corresponding number of counterions using tleap software. The lipid bilayer contained 20 % ergosterol (CID 444679), 20 % DOPG (CID 11846228) and 60 % DOPE (CID 11597900) to better mimic biological membrane conditions. CHARMM-GUI membrane builder outputs were generated in PDB format and imported into MOE. Amber FF14SB was used to model protein residues, Lipid17 (or Lipid25) for lipids including ergosterol, and GAFF2 for ADP and Pi molecules. Experimentally resolved ergosterol positions were kept in the system, inserted in the membrane. Water molecules, Mg^2+^ ions, and counterions were modeled using the TIP3P water model along with monovalent and divalent ion parameters. The system was simulated with periodic boundary conditions. A 10 Å cutoff was applied for nonbonded interactions, including Coulomb and van der Waals potentials. Long-range electrostatics were treated using the particle mesh Ewald method.R.

Minimization and thermalization of the systems and MD simulations were carried out with Amber22 using CPU and GPU PMEMD versions. Minimization was carried out in three steps by sequentially minimizing: (i) water O-atoms (2,000 steps); (ii) all bonds involving H-atoms (5,000 steps); (iii) whole system (5,000 steps). Each system was then thermalized in two steps: (i) water molecules were thermalized to 100 K during 50 ps under (N,P,T) ensemble conditions using a 2 fs time integration in semi-isotropic conditions using Berendsen barostat; (ii) the whole system was then thermalized from 100 K to 300 K during 100 ps under (N,P,T) ensemble conditions with same condition for barostat. A pre-production step was introduced with semi-isotropic scaling barostat at 300 K using the production parameters for 100 ns. During this step we ensured that periodic boundary conditions parameters reached equilibrium. During this phase, the membrane was kept in contact with the protein and with the limit of the periodic boundary conditions. This pre-production phase was not included in the analyses.

Production runs were then carried out for 700 ns in triplicate starting from the same final equilibrated pre-production state. The production run with 2 fs integration timestep under (N,P,T) ensemble conditions with semiisotropic scaling. Temperature was maintained using the Andersen-like thermostat. Constant pressure, set at 1 bar was maintained with semi-isotropic pressure scaling using a Berendsen barostat. Only production trajectories were analyzed and presented. Visualizations of coordinate projections were generated with VMD software and tcl scripts.

## Acknowledgments

We thank Pr. Dominique Sanglard for the gift of clinical isolates of *Candida glabrata* DSY562 and DSY565. We also gratefully acknowledge support from the CNRS/IN2P3 Computing Center (Lyon - France) for providing computing and data-processing resources needed for this work.” A special thanks to Alexis Michon and his team for management of the IBCP computing facilities.

This work used the platforms of the Grenoble Instruct-ERIC center (ISBG; UAR 3518 CNRS-CEA-UGA-EMBL) within the Grenoble Partnership for Structural Biology (PSB), supported by FRISBI and GRAL, financed within the University Grenoble Alpes graduate school (Ecoles Universitaires de Recherche). Cryo-EM sample screening, optimization, and data collections for the apo and ITC-bound protein were performed at the Cryo-EM Swedish National Facility in Stockholm, Sweden funded by the Knut and Alice Wallenberg, Family Erling Persson and Kempe Foundations, SciLifeLab, Stockholm University, and Umeå University.

## Funding

J.P.’s Ph. D was supported by the doctoral school EDISS 205 and R&D BOOSTER – IMABGEN grants, Agence Nationale de la Recherche (Française), ((French) National Research Agency): ANR-24-CE44-3527 RESTOR to PFA and ABO as coordinators and VCH, RTE, EBE and SAG as partners, Auvergne-Rhone-Alpes Region innovation program: R&D booster to PFA, VCH, MDU and AMO,

INSTRUCT-ERIC grant number 24618 to VCH, PFA, LZE and GSH,

FRISBI (ANR-10-INBS-0005-02) and GRAL, financed within the University Grenoble Alpes graduate school (Ecoles Universitaires de Recherche) CBH-EUR-GS (ANR-17-EURE-0003) to GSH and LZE.

The IBS-ISBG EM facility is supported by the Auvergne-Rhône-Alpes Region, the Fondation Recherche Medicale (FRM), the fonds FEDER and the GIS-Infrastructures en Biologie Santé et Agronomie (IBISA),

The Knut and Alice Wallenberg foundation (2023.0201) and the Swedish research council (2021-03992) to BWI and MHO.

RCA and ELA acknowledge Marsden Fund grant MFP-UOO2210 awarded by the Royal Society of New Zealand.

## Author contributions

Conceptualization: PFA, JPA, VCH

Methodology: PFA, JPA, VCH, BWI, EZA, RBA, NSA, CDE, AMO, ABA, SAG, EBE, MDU, RTE, ELA,

Investigation: PFA, JPA, VCH, BWI, EZA

Funding acquisition: PFA

Project administration: PFA

Supervision: PFA

Writing – original draft: PFA, JPA, VCH

Writing – review & editing: PFA, JPA, VCH, BWI, EZA, RBA, NSA, CDE, AMO, ABA, SAG, EBE, MDU, RTE, GSH, MHO, ABO, ELA, RCA, RPR

## Data and materials availability

3D Data from variability refinements have been deposited in the public database Zenodo (see Supplementary Table 2). Other data are available in the main text or the supplementary materials. Material and correspondence should be addressed to Pierre Falson and Vincent Chaptal.

## Supplementary Information

Supplementary Figures 1 to 15

Supplementary Tables 1 and 2

Supplementary Movies 1 and 2

Supplementary Data 1 and 2

