## Supplementary Information for "Cryo-EM of ATP-driven dynamics and itraconazole binding of the fungal drug efflux ABC pump *Candida glabrata* Cdr1"

Pata *et al.*,

**This PDF file includes:**

Supplementary Figures 1 to 15

Supplementary Tables 1 and 2

Supplementary Movies 1 to 3

Supplementary Data 1 and 2

### Supplementary Figure 1

### A

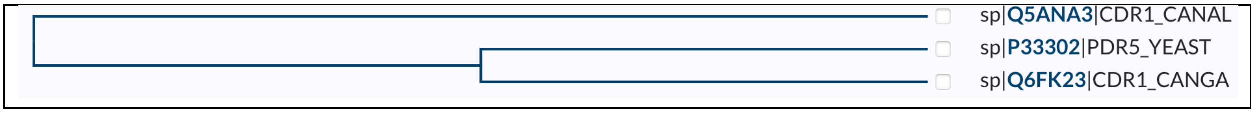

### B

| %Id | *ca*Cdr1 | *cg*Cdr1 | Pdr5 |
|---|---|---|---|
| *Ca*Cdr1 | 100 | 54 | 56 |
| *Cg*Cdr1 | | 100 | 74 |
| Pdr5 | | | 100 |

### C

```
                                              NBD-1->
              i         i         i         i         i         i         i
CgCdr1        ---MSLASDKKD-ADVASTTTTAQDDDNLSTYHGFDHHVQDQVRQLARTLTQQSSLHQKK 0056
CaCdr1        MSDSKMSSQDESKLEKAISQDSSSENHSINEYHGFDAHTSENIQNLARTFTHDSFKDDSS 0060
Pdr5          MPEAKLNNNVNDVTSYSSASSSTENAADLHNYNGFDEHTEARIQKLARTLTAQSMQNSTQ 0060
                  .: .: :.  . : :  ::.:  .:  *:*** *.. .:::****:* :*   ....
CgCdr1 2D                                         H-1
Apo           ---xxxxxxxxx-xxxxxxxxxxxxxxxxlllllllhhhhhhhhhhhhhhhhllllxxxx
ATPADPViMg    ---xxxxxxxxx-xxxxxxxxxxxxxxxxxxxxxxxxxxxxxxxxxxxxxxxxxxxxxx

              i         i         i         i         i         i         i
CgCdr1        ----------EHTLPEEGINPIFTNTEADDYNPRLDPTSDEFSSAEWVQNMSNISNSDPD 0106
CaCdr1        AGLLK---YLTHMSEVPGVNPYEHE---EINNDQLNPDSENFNAKFWVKNLRKLFESDPE 0114
Pdr5          SAPNKSDAQSIFSSGVEGVNPIFSDPEAPGYDPKLDPNSENFSSAAWVKNMAHLSAADPD 0120
                          .     *:**    :      : :*:* *::*.:  **:*: ::  :**:
CgCdr1 2D                         B-1                 B-2 H-2
Apo          -----------xxxxxxxxxxxxxxxxxxxxxxxxxlllllllhhhhhhhhhhhhhhhll
ATPADPViMg   -----------xxlllllllleeeelllllllllllllllleeehhhhhhhhhhhhhhhll

                                                               RD-1N->
              i         i         i         i         i         i         i
CgCdr1        YYKPYSLGCYWKDLVATGESADIEYQANFLNGPYKGLKTVYNTVVPSTASSKDKNFKILK 0166
CaCdr1        YYKPSKLGIGYRNLRAYGVANDSDYQPTVTNALWKLATEGFRHF---QKDDDSRYFDILK 0171
Pdr5          FYKPYSLGCAWKNLSASGASADVAYQSTVVNIPYKILKSGLRKF---QRSKETNTFQILK 0177
              :***  .**   :::* * * : *   **  .. *   :*   .   . .     ... . *.***
CgCdr1 2D       B-3    B-4    B-5          H-3
Apo           lllllllleeeelleeeeeexxxxxxxxxxxxxxxxxxxxxxxxxxxxxxxxxxxxlllll
ATPADPViMg    lleeeelleeelleeeellllllllllllhhhhhhhhhhhhhhhlxxxxxxxllllllll

                           ncWalker-A
              i         i         i         i         i         i         i
CgCdr1        SMEGAVNPGELLVVLGRPGSGCTTLLKSISSNTHGFNIAKESTISYSGMTPNDIRKHFRG 0226
CaCdr1        SMDAIMRPGELTVVLGRPGAGCSTLLKTIAVNTYGFHIGKESQITYDGLSPHDIERHYRG 0231
Pdr5          PMDGCLNPGELLVVLGRPGSGCTTLLKSISSNTHGFDLGADTKISYSGYSGDDIKKHFRG 0237
               *:.  :.**** *******:**:****:*: **:**.:.  :: *:*.* : .**.:*:**
CgCdr1 2D     B-6       B-7         H-4                   B-7       H-5
Apo           eeeeeellllleeeellllllhhhhhhhhhllllllllllleeeeellllhhhhhhhlll
ATPADPViMg    lleeelllleeeeeellllllhhhhhhhhllllllllllllleeeelllllhhhhhhhhl
```

```
                <-RD-1N
            HD-1->
             ncQ-loop
             i         i         i         i         i         i         i
CgCdr1        EVVYNAEADIHLPHLTVYQTLLTVARLKTPQNRLKGIDRETYARHLTEVAMATFGLSHTR 0286
CaCdr1        DVIYSAETDVHFPHLSVGDTLEFAARLRTPQNRGEGIDRETYAKHMASVYMATYGLSHTR 0291
Pdr5          EVVYNAEADVHLPHLTVFETLVTVARLKTPQNRIKGVDRESYANHLAEVAMATYGLSHTR 0297
              :*:*.**:*:*:*:***:*  :**   .***:*****  :*:***:**.*::.* ***:******
CgCdr1 2D      B-6   B-7      H-6                    H-7
Apo           leeeelleeeeellllhhhhhhhhhhhlllllllllllhhhhhhhhhhhhhhhlllllll
ATPADPViMg    leeeelleeeeellllhhhhhhhhhhhlllllllllllhhhhhhhhhhhhhhhlllllll

                                                   <- HD-1
                                              RD-1C->
                       cABC Signature               ncWalker-B ncD-loop
             i         i         i         i         i         i         i
CgCdr1        NTKVGNDLVRGVSGGERKRVSIAEVSICGSKFQCWDNATRGLDSATALEFIRALKVQASI 0346
CaCdr1        NTNVGNDFVRGVSGGERKRVSIAEASLSGANIQCWDNATRGLDSATALEFIRALKTSAVI 0351
Pdr5          NTKVGNDIVRGVSGGERKRVSIAEVSICGSKFQCWDNATRGLDSATALEFIRALKTQADI 0357
              **:****:****************.*:.*:::**********************..* *
CgCdr1 2D                    H-8              B-8         H-9
Apo           llllllllllllllhhhhhhhhhhhhhhllleeeeellllllhhhhhhhhhhhhhhhhh
ATPADPViMg    llllllllllllllhhhhhhhhhhhhhhllleeeeellllllhhhhhhhhhhhhhhhhh

                      ncH-loop
             i         i         i         i         i         i         i
CgCdr1        SNAAATVAIYQCSQDAYDLFDKVCVLYDGYQIYFGPAGKAKEYFQKMGYVSPERQTTADF 0406
caCdr1        LDTTPLIAIYQCSQDAYDLFDKVVVLYEGYQIFFGKATKAKEYFEKMGWKCPQRQTTADF 0411
Pdr5          SNTSATVAIYQCSQDAYDLFNKVCVLDDGYQIYYGPADKAKKYFEDMGYVCPSRQTTADF 0417
               :::  :*************:**  **  :****::*  *  ***:**:.**:  .*.*******
CgCdr1 2D       B-8         H-10     B-9      B-10       H-11      B-11     H-12
Apo           hlleeeeelllllhhhhhhlleeeeeelleeeeeeelllhhhhhhhhleeeellllhhhh
ATPADPViMg    hllleeeelllllhhhhhhlleeeeeellleeeeeelllhhhhhhhhleeeellllhhhh

                                                                <-RD-1C
             i         i         i         i         i         i         i
CgCdr1        LTAVTSPSERIINQDYINRGIFVPQTPKEMWEYWRASEDHADLIKEIDSKLSDNYD-ANL 0465
CaCdr1        LTSLTNPAEREPLPGYE---DKVPRTAQEFETYWKNSPEYAELTKEIDEYFVECERSNTR 0468
Pdr5          LTSVTSPSERTLNKDMLKKGIHIPQTPKEMNDYWVKSPNYKELMKEVDQRLLNDDE-ASR 0476
              **::*.*:**      .       :*:*  :*:   **   *  ::  :*  **:*.  :  :      .
CgCdr1 2D                  H-13         H-14       H-15                H-16
Apo           hhhllllllllllhhhhhllllllllhhhhhhhhhllhhhhhhhhhhhhhhhxxxx-xxx
ATPADPViMg    hhhhlllllllllhhhhhhlllllllhhhhhhhhhllhhhhhhhhhhhhhhhlllh-hhh

                    H-17: Connecting helix 1 (CnH-1)  TMD-1->
             i         i         i         i         i         i         i
CgCdr1        AEIKDAHVARQSKRARPSSPYTVSYGMQIKYLLIRNFWRIKQSSGVTLFMVIGNSSMAFI 0525
CaCdr1        ETYRESHVAKQSNNTRPASPYTVSFFMQVRYGVARNFLRMKGDPSIPIFSVFGQLVMGLI 0528
Pdr5          EAIKEAHIAKQSKRARPSSPYTVSYMMQVKYLLIRNMWRLRNNIGFTLFMILGNCSMALI 0536
                 :::*:*:*:**:.:**:******:  **::*  :  **:  *::  .  ..  :*  ::*:    *.:*
cgCdr1 2D                                H-17                  TMH-1
Apo           xxxxxxxxxxxxxxxxxlllllllhhhhhhhhhhhhhhhhhhlllhhhhhhhhhhhhhhh
```

```
ATPADPViMg      hhhhhhhhhhllllllllllllllhhhhhhhhhhhhhhhhhhlllhhhhhhhhhhhhhhh
                      ECD-1                         H-19: Coupling helix 1 (CpH-1)

               i         i         i         i         i         i         i
CgCdr1          LGSMFYKVMKHNTTSTFYFRGAAMFFAVLFNAFSSLLEIFSLFEARPITEKHRTYSLYHP 0585
CaCdr1          LSSVFYNLSQ--TTGSFYYRGAAMFFAVLFNAFSSLLEIMSLFEARPIVEKHKKYALYRP 0586
Pdr5            LGSMFFKIMKKGDTSTFYFRGSAMFFAILFNAFSSLLEIFSLYEARPITEKHRTYSLYHP 0596
                *.*:*::: :   *.:**:**:*****:***********:**:*****.***:.*:**:*
CgCdr1 2D             H-18     TMH-2                         H-19          TMH-3
Apo             hhhhhlhhhhlllllhhhhhhhhhhhhhhhllllllllhhhhhhlhhhhhhhhhlllllh
ATPADPViMg      hhhhhllllllllhhhhhhhhhhhhhhhhhhhhhhhhhhhhhhlllhhhhhhhhhlllllh

               i         i         i         i         i         i         i
CgCdr1          SADAFASILSEVPAKLITAVCFNIIYYFLVNFRRNGGVFFFYFLINIVAVFAMSHLFRCV 0645
CaCdr1          SADALASIISELPVKLAMSMSFNFVFYFMVNFRRNPGRFFFYWLMCIWCTFVMSHLFRSI 0646
Pdr5            SADAFASVLSEIPSKLIIAVCFNIIFYFLVDFRRNGGVFFFYLLINIVAVFSMSHLFRCV 0656
                ****:**::**:* **  ::.**:::**:*:**** * **** *: * ..* ******.:
CgCdr1 2D                                          TMH-4
Apo             hhhhhhhhhhhhlhhhhhhhhhhhhhhhhllllllhhhhhhhhhhhhhhhhhhhhhhhhh
ATPADPViMg      hhhhhhhhhhhhlhhhhhhhhhhhhhhhhllllllhhhhhhhhhhhhhhhhhhhhhhhhh

               i         i         i         i         i         i         i
CgCdr1          GSVSKTLSAAMVPASMLLLGLSMYSGFAIPRTKILGWSKWIWYINPLAYLFESLMINEFH 0705
CaCdr1          GAVSTSISGAMTPATVLLLAMVIYTGFVIPTPSMLGWSRWINYINPVGYVFESLMVNEFH 0706
Pdr5            GSLTKTLSEAMVPASMLLLALSMYTGFAIPKKKILRWSKWIWYINPLAYLFESLLINEFH 0716
                *:::.::* **.**::***.: :*:**.**  .:* **:** ****:.*:****::****
CgCdr1 2D             TMH-5                        H-20      H-21
Apo             hhhllllhhhhhhhhhhhhhhhhhhlllllllllllllhhhhhhllhhhhhhhhhhhhhhl
ATPADPViMg      hhhlllhhhhhhhhhhhhhhhhhhhllllllllllhhhhhhhhhlhhhhhhhhhhhhhhl

               i         i         i         i         i         i         i
CgCdr1          DRKFPCSQYIPSGSVYNNVPADSRICSSVGAIRGNDYVLGDDFLRESYSYLHKHKWRGFG 0765
CaCdr1          GREFQCAQYVPSGPGYENISRSNQVCTAVGSVPGNEMVSGTNYLAGAYQYYNSHKWRNLG 0766
Pdr5            GIKFPCAEYVPRGPAYANISSTESVCTVVGAVPGQDYVLGDDFIRGTYQYYHKDKWRGFG 0776
                . :* *::*:* *  * *:   . :*: **:: *:: * * :::  :*.* :..***.:*
CgCdr1 2D         B-12       H-22a  H-22b           B-13 H-23      H-24 TMH-6
Apo             lleeelllllllllllllllhhhllllllllllllleeehhhhhhhhllllhhhllhhhh
ATPADPViMg      lleeellllllllhhhllllhhhllllllllllllleeehhhhhhhhllllhhhhhhhhh

                                       Linker
               i         i         i         i         i         i         i
CgCdr1          IGLAYVIFFLVLYLILCEYNEGAKQKGEILVFPQNIVRRMKKERKLKNVSSDNDVEIGDV 0825
CaCdr1          ITIGFAVFFLAIYIALTEFNKGAMQKGEIVLFLKGSLKKHKRKTAASNKGDIEAGPVAGK 0826
Pdr5            IGMAYVVFFFFVYLFLCEYNEGAKQKGEILVFPRSIVKRMKKRGVLTEKNANDPENVGER 0836
                * :.:.:**: :*: * *:*:** *****::* :. ::: *:.   .: .  :   :.
CgCdr1 2D                                         H-24b
Apo             hhhhhhhhhhhhhhhhhlllxxxxxxxxxxxxxxxxxxxxxxxxxxxxxxxxxxxxxxxx
ATPADPViMg      hhhhhhhhhhhhhhhhhhllllllllllllllllhhhhhhhhlllllxxxxxxxxxxxxx

                                       NBD-2->
                                        RD-2N->
               i         i         i         i         i         i         i
CgCdr1          SDIS-DKKILADSSDESEESGANIGLSQSEAIFHWRNLCYDVQIKKETRRILNNVDGWVK 0884
CaCdr1          LDYQDEAEAVNNEKFTEKGSTGSVDFPENREIFFWRDLTYQVKIKKEDRVILDHVDGWVK 0886
Pdr5            SDLSSDRKMLQESSEEESDTYGEIGLSKSEAIFHWRNLCYEVQIKAETRRILNNVDGWVK 0896
                 * . : : : :..  .. : ..:.: :.. **.**:* *:*:** * * **::******
```

```
CgCdr1 2D                            B-14  B-15   B-15b B-16   B-16b B-17
Apo         xxxx-xxxxxxxxxxxxxxxxxxlllleeeleeeeeeeeeeeeelleeeeeelleeeeee
ATPADPViMg  xxxx-xxxxxxxxxxxxxlllllllleeeeeeleeeelleeellleeeeeeleeeeleeee

                                                                    <-RD-2N
                                                                HD-2->
                   ^cWalker-A                                  ^cQ-loop
i         i         i         i         i         i         i
CgCdr1      PGTLTALMGASGAGKTTLLDCLAERVTMGVIT-GEVSVDGKQRDDSFARSIGYCQQQDLH 0943
CaCdr1      PGQITALMGASGAGKTTLLNCLSERVTTGIITDGERLVNGHALDSSFQRSIGYVQQQDVH 0946
Pdr5        PGTLTALMGASGAGKTTLLDCLAERVTMGVIT-GDILVNGIPRDKSFPRSIGYCQQQDLH 0955
            ** :***************:**:**** *:** *:  *:*   *.** ***** ****:*
CgCdr1 2D      B-18       H-25           B-19 B-19b            B-20
Apo         llleeeeeelllllllhhhhhhhlllllleee-lllllllllllllllllleeellllll
ATPADPViMg  llleeeeeellllllhhhhhhhhllllllleee-leeeeeleeeellllllllllllleeellllll

                                                               ^ncABC signature
           i         i         i         i         i         i         i
CgCdr1      LKTSTVRESLRFSAYLRQPADVSIEEKNQYVEDVIKILEMEQYADAVVGVPGEGLNVEQR 1003
CaCdr1      LETTTVREALQFSAYLRQSNKISKKEKDDYVDYVIDLLEMTDYADALVGVAGEGLNVEQR 1006
Pdr5        LKTATVRESLRFSAYLRQPAEVSIEEKNRYVEEVIKILEMEKYADAVVGVAGEGLNVEQR 1015
            *:*:****:*:*******  .:* :**: **: **.:*** .****:*** *********
CgCdr1 2D        H-26              H-27                             H-28
Apo         lllllhhhhhhhhhhhhlllllllhhhhhhhhhhhhhllllllllllllllllllllhhhh
ATPADPViMg  lllllhhhhhhhhhhhhllllllhhhhhhhhhhhhhhllllllllllllllllllllhhhh

                        <-HD-2
                   RD-2C->
                        ^cWalker-B ^cD-loop                 ^cH-loop
           i         i         i         i         i         i         i
CgCdr1      KRLTIGVELAAKPKLLVFLDEPTSGLDSQTAWSICQLMKKLANHGQAILCTIHQPSAILM 1063
CaCdr1      KRLTIGVELVAKPKLLLFLDEPTSGLDSQTAWSICKLMRKLADHGQAILCTIHQPSALIM 1066
Pdr5        KRLTIGVELTAKPKLLVFLDEPTSGLDSQTAWSICQLMKKLANHGQAILCTIHQPSAILM 1075
            *********.******:******************:**:***:**************::*
CgCdr1 2D                  B-21        H-29               B-22      H-30
Apo         hhhhhhhhhllllllleeeelllllllhhhhhhhhhhhhhhhhhllleeeellllllhhhh
ATPADPViMg  hhhhhhhhhhllllleeeellllllllllhhhhhhhhhhhhhhhlleeeellllllhhhh

           i         i         i         i         i         i         i
CgCdr1      QEFDRLLFLQRGGKTVYFGDLGDGCKTMIDYFESHGSHKCPPDANPAEWMLEVVGAAPGS 1123
CaCdr1      AEFDRLLFLQKGGRTAYFGELGENCQTMINYFEKYGADPCPKEANPAEWMLQVVGAAPGS 1126
Pdr5        QEFDRLLFMQRGGKTVYFGDLGEGCKTMIDYFESHGAHKCPADANPAEWMLEVVGAAPGS 1135
             *******:*:**:*.***:**:.*:***:***.:*:. ** :********:********
CgCdr1 2D        B-23    B-24    H-31                    H-32
Apo         hlllleeeeellleeeeeellhhhhhhhhhhhhhllllllllllllhhhhhhhhlllllll
ATPADPViMg  hhllleeeeellleeeeeellllllhhhhhhhhhhllllllllllhhhhhhhhhllllll

                                                   <-RD-2C
                                             H-35: Connecting helix 2 (CnH-2)
           i         i         i         i         i         i         i
CgCdr1      HANQDYHEVWRNSDEYQKVQEELEWMSNELPKKNTNNSETVHKEFATGVLYQCKLVSLRL 1183
CaCdr1      HAKQDYFEVWRNSSEYQAVREEINRMEAELSKLPRDNDPEALLKYAAPLWKQYLLVSWRT 1186
Pdr5        HANQDYYEVWRNSEEYRAVQSELDWMERELPKKGSITAAEDKHEFSQSIIYQTKLVSIRL 1195
             **:***.******.**: *:.*:: *. ** *    .      :::  :  *  *** *
CgCdr1 2D        H-33    H-34                               H-35
```

```
Apo         llllllhhhhhllhhhhhhhhhhhhhhhhhhhllllllllllllllllhhhhhhhhhhhh
ATPADPViMg  lllllhhhhhhhllhhhhhhhhhhhhhhhllllllllllllllllllllhhhhhhhhhhh

                   TMD-2
           i         i         i         i         i         i         i
CgCdr1      FQQYWRSPDYLWSKFFLTIFNNIFIGFTFFKADRSLQGLQNQMLAVFMFTVIFNPLLQQY 1243
CaCdr1      IVQDWRSPGYIYSKIFLVVSAALFNGFSFFKAKNNMQGLQNQMFSVFMFFIPFNTLVQQM 1246
Pdr5        FQQYWRSPDYLWSKFILTIFNQLFIGFTFFKAGTSLQGLQNQMLAVFMFTVIFNPILQQY 1255
            : * ****.*::**::*.:   :* **:****  .:*******::**** : ** ::**
CgCdr1 2D          TMH-7                       TMH-8
Apo         hhhhhllhhhhhhhhhhhhhhhhhhhhhlllllllllllhhhhhhhhhhhllhhhhhhhh
ATPADPViMg  hhhhhllhhhhhhhhhhhhhhhhhhhhhllxxxxlhhhhhhhhhhhhhhhhhhhhhhhhh

         H-36: Coupling helix 2 (CpH-2)                           ECD-2
           i         i         i         i         i         i         i
CgCdr1      LPSFVQQRDLYEARERPSRTFSWKAFIVSQILVEIPWNILAGTVAFVIYYYAIGFYSNAS 1303
CaCdr1      LPYFVKQRDVYEVREAPSRTFSWFAFIAGQITSEIPYQVAVGTIAFFCWYYPLGLYNNAT 1306
Pdr5        LPSFVQQRDLYEARERPSRTFSWISFIFAQIFVEVPWNILAGTIAYFIYYYPIGFYSNAS 1315
            ** **:***:**.** ******* :** .**  *:*::: .**:*:. :** :*:*.**:
CgCdr1 2D    H-36                 TMH-9                           H-37
Apo         hhhhhhhhhhhhhllllllllllhhhhhhhhhhhhhhhhhhhhhhhhhhhhhhhlhhhhhh
ATPADPViMg  hhhhhhhhhhhhhllllllllllhhhhhhhhhhhhhhhhhhhhhhhhhhlllllllhhhhh

           i         i         i         i         i         i         i
CgCdr1      VAHQLHERGALFWLFSCAFYVVYIGSLALFCISFNQVAEAAANMASLMFTLSLSFCGVLVT 1363
CaCdr1      PTDSVNPRGVLMWMLVTAFYVYTATMGQLCMSFSELADNAANLATLLFTMCLNFCGVLAG 1366
Pdr5        AAGQLHERGALFWLFSCAFYVVYVGSMGLLVISFNQVAESAANLASLLFTMSLSFCGVMTT 1375
             : .:: **.*:*::  ***** .::. : :**.::*: ***:*:*:**:.*.****:.
CgCdr1 2D       TMH-10                          TMH-11
Apo         hhlllhhhhhhhhhhhhhhhhhhhhhhhhhhhhlllhhhhhhhhhhhhhhhhhhhlllll
ATPADPViMg  hlllhhhhhhhhhhhhhhhhhhhhhhhhhhhhllllhhhhhhhhhhhhhhhhhlllllll

           i         i         i         i         i         i         i
CgCdr1      PNGMPRFWIFMYRVSPLTYLIDGMLSTGVANVAIKCSNYELLRFSPAANLTCGEYLGPYL 1423
CaCdr1      PDVLPGFWIFMYRCNPFTYLVQAMLSTGLANTFVKCAEREYVSVKPPNGESCSTYLDPYI 1426
Pdr5        PSAMPRFWIFMYRVSPLTYFIQALLAVGVANVDVKCADYELLEFTPPSGMTCGQYMEPYL 1435
            *. :* ******* .*:**:::.:*:.*:**. :**:: * : ..*  . :*. *: **:
CgCdr1 2D         H-38     H-39                      B-25      H-40 H-41
Apo         llllllhhhhhhhhlhhhhhhhhhhhhhllllllllllllleeelllllllhhhhlhhhh
ATPADPViMg  llllllhhhhhhhhlhhhhhhhhhhhhhllllllllllllleeelllllllhhhhlhhhh

           i         i         i         i         i         i         i
CgCdr1      QTVKTGYIVDPSATDTCELCPYSHTNDFLSSVSSKYSRRWRNWGIFICYIAFNYIAGIFL 1483
CaCdr1      KFAGG-Y-FETRNDGSCAFCQMSSTNTFLKSVNSLYSERWRNFGIFIAFIAINIILTVIF 1484
Pdr5        QLAKTGYLTDENATDTCSFCQISTTNDYLANVNSFYSERWRNYGIFICYIAFNYIAGVFF 1495
            : .   *  :    .:* :*  * ** :* .*.* **.****:****.:**:* *  :::
CgCdr1 2D                   B-26    H-42       TMH-12
Apo         hhhllllllllllllleeelllllhhhhhhhlllllllhhhhhhhhhhhhhhhhhhhhhh
ATPADPViMg  hhhllllllllllllleeelllllhhhhhhhllllhhhhhhhhhhhhhhhhhhhhhhhhh

           i         i
CgCdr1      YWLARVPKKSGKLAKK- 1499
CaCdr1      YWLARVPKGNREKKNKK 1501
Pdr5        YWLARVPKKNGKLSKK- 1511
            ******** . :  :*
```

```
CgCdr1 2D
Apo          hhhhlllxxxxxxxxx-
ATPADPViMg   hhhhllllxxxxxxxx-
```

**D**

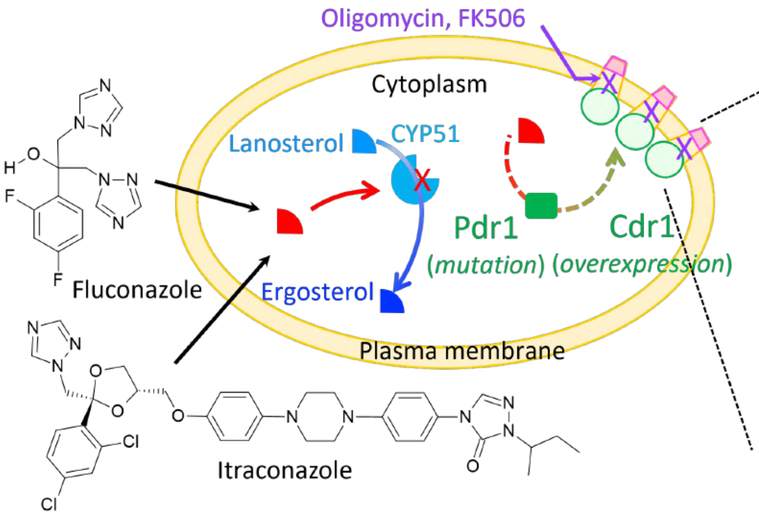

**E**

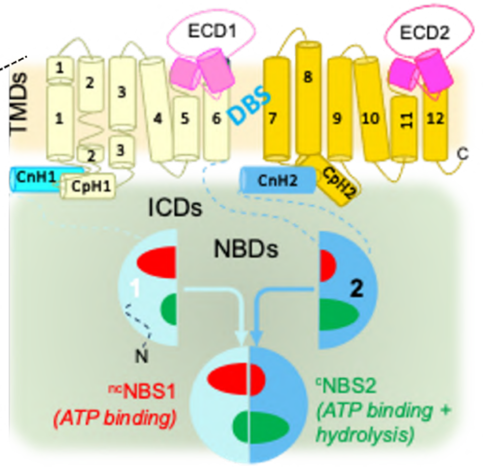

**Supplementary Figure 1. Phylogeny, similarity, primary sequence alignment of *Ca*Cdr1, *Cg*Cdr1 and Pdr5, 2D structures of molecular models resolved in the study and mechanism of azole antifungal resistance.** Sequences were taken from Uniprot, Q5ANA3 for *Ca*Cdr1, Q6FK23 for *Cg*Cdr1 and P33302 for Pdr5. **A**. Phylogeny. **B**. Identity matrix. **C**. Clustal alignment. Symbols "*", ":" and "." correspond to identical, functionally very close and close residues, respectively. Data were generated with the Align tool of UniProt. Main motifs are also indicated. The primary sequence of *Cg*Cdr1 is repeated for the apo and [$^{s}$0|$^{nc}$ATP|$^{c}$ADP-Vi-Mg] cryo-EM structures of this study, showing the different domains and secondary structures of residues involved in loops (l), helices (h) and sheets (e), colored as in Figure 2: pale blue and blue for NBD-1 and NBD-2, yellow and dark yellow for TMD-1 and TMD-2, pink and magenta for ECD-1 and ECD-2. HD and RD correspond to the Helical and RecA sub-domains of each NBD. **D.** Mechanism of Cdr1 overexpression induced by azole antifungal stress. In response to Cyp51 inhibition by azoles, yeast upregulate Cdr1 expression by approximately ten-fold through mutations of the zinc-cluster transcription factor, Pdr1, enhancing the efflux of azoles. This process can be blocked by inhibitors such as oligomycin or FK506. **E.** Cdr1 topology. Cdr1 consists of extracellular domains (ECDs), transmembrane domains (TMDs), intracellular domains (ICDs), and nucleotide-binding domains (NBDs). When assembled, NBD-1 and NBD-2 form two nucleotide-binding sites: a catalytic site (NBS-2, dark green) that hydrolyzes ATP and a non-catalytic site (NBS-1, red). Drugs bind to the drug-binding site (DBS) and translocate through the TMD to the cell exterior. Structural and functional coupling of the NBDs with the TMDs is mediated by connecting helices (CnH-1, CnH-2) and coupling helices (CpH-1, CpH-2).

**Supplementary Figure 2**

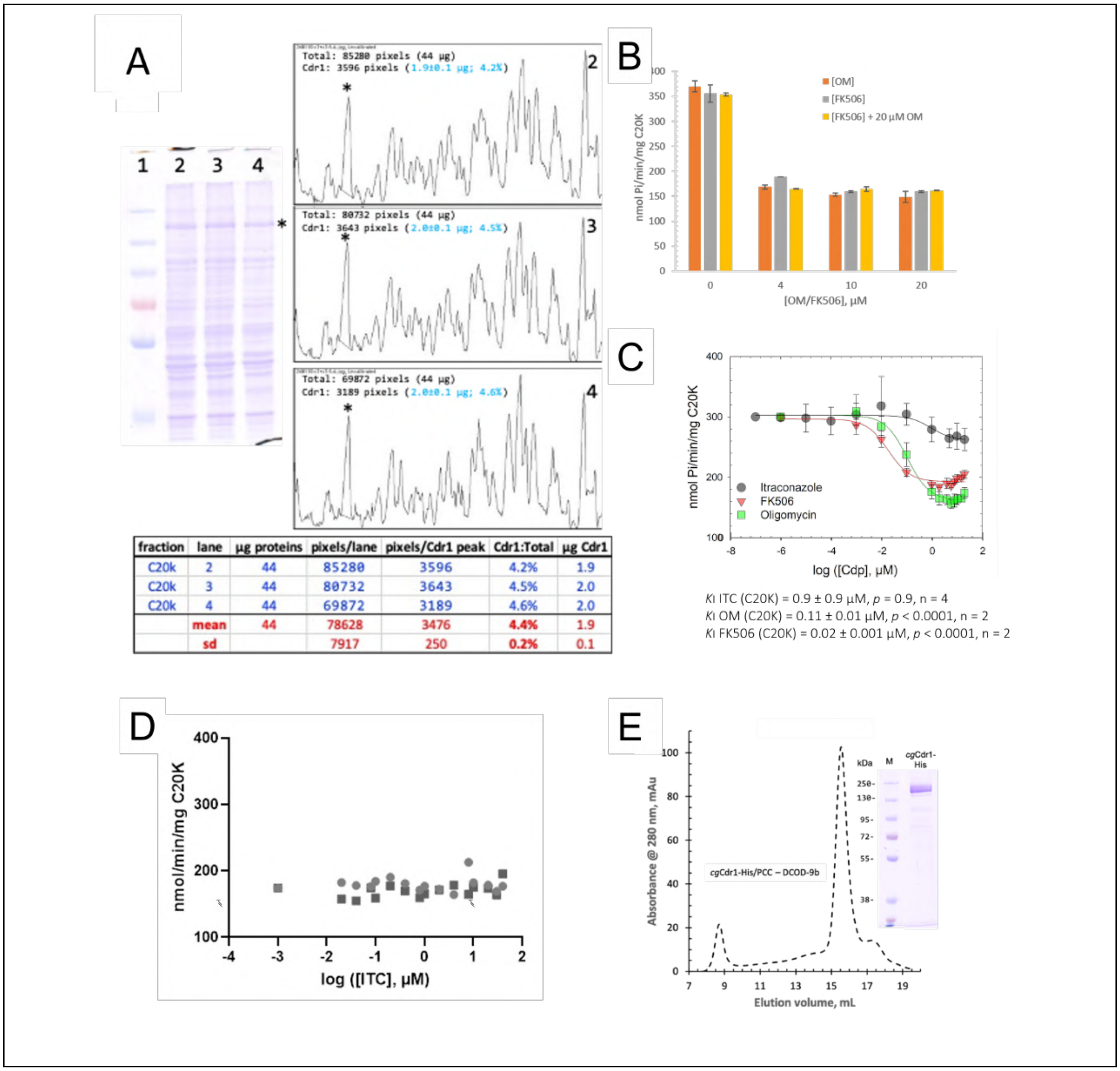

| fraction | lane | µg proteins | pixels/lane | pixels/Cdr1 peak | Cdr1:Total | µg Cdr1 |
|---|---|---|---|---|---|---|
| C20k | 2 | 44 | 85280 | 3596 | 4.2% | 1.9 |
| C20k | 3 | 44 | 80732 | 3643 | 4.5% | 2.0 |
| C20k | 4 | 44 | 69872 | 3189 | 4.6% | 2.0 |
| | mean | 44 | 78628 | 3476 | 4.4% | 1.9 |
| | sd | | 7917 | 250 | 0.2% | 0.1 |

Supplementary Figure 2. Production and purification of *Cg*Cdr1. A. Quantification of *Cg*Cdr1 in *S. cerevisiae* membrane fractions. Forty-four micrograms of membrane fraction enriched in *Cg*Cdr1 pelleted at 20,000 x g (see Methods) were loaded in triplicate and run on a 10% SDS PAGE, Coomassie-stained and then quantified with ImageJ. B. Oligomycin and FK506 inhibition of the ATPase activity of *Cg*Cdr1 in the C20K membrane fraction. ATPase activity assayed by Pi release as a function of oligomycin (OM) or FK506 concentration with or without 20 µM oligomycin. C. ATPase sensitivity to inhibitors and substrates of *Cg*Cdr1 in C20K membranes. D. Absence of additional inhibition of the ATPase activity of *Cg*Cdr1 in the C20K membrane fraction by itraconazole and oligomycin. Membranes were incubated as in panel B with 20 µM oligomycin and the indicated concentrations of itraconazole. E. Superose 200 10/300 chromatogram profile of *Cg*Cdr1 solubilized with 70 µM PCC and 4 µM DCOD-9b and purified as described in Methods.

SDS-PAGE of the main peak. Image taken with permission from Pata J, Moreno A, Wiseman B, Magnard S, Lehlali I, Dujardin M, Banerjee A, Högbom M, Boumendjel A, Chaptal V *et al* (2023) Purification and characterization of Cdr1, the drug-efflux pump conferring azole resistance in *Candida* species. *Biochimie* 220: 167–178.

.

**Supplementary Figure 3**

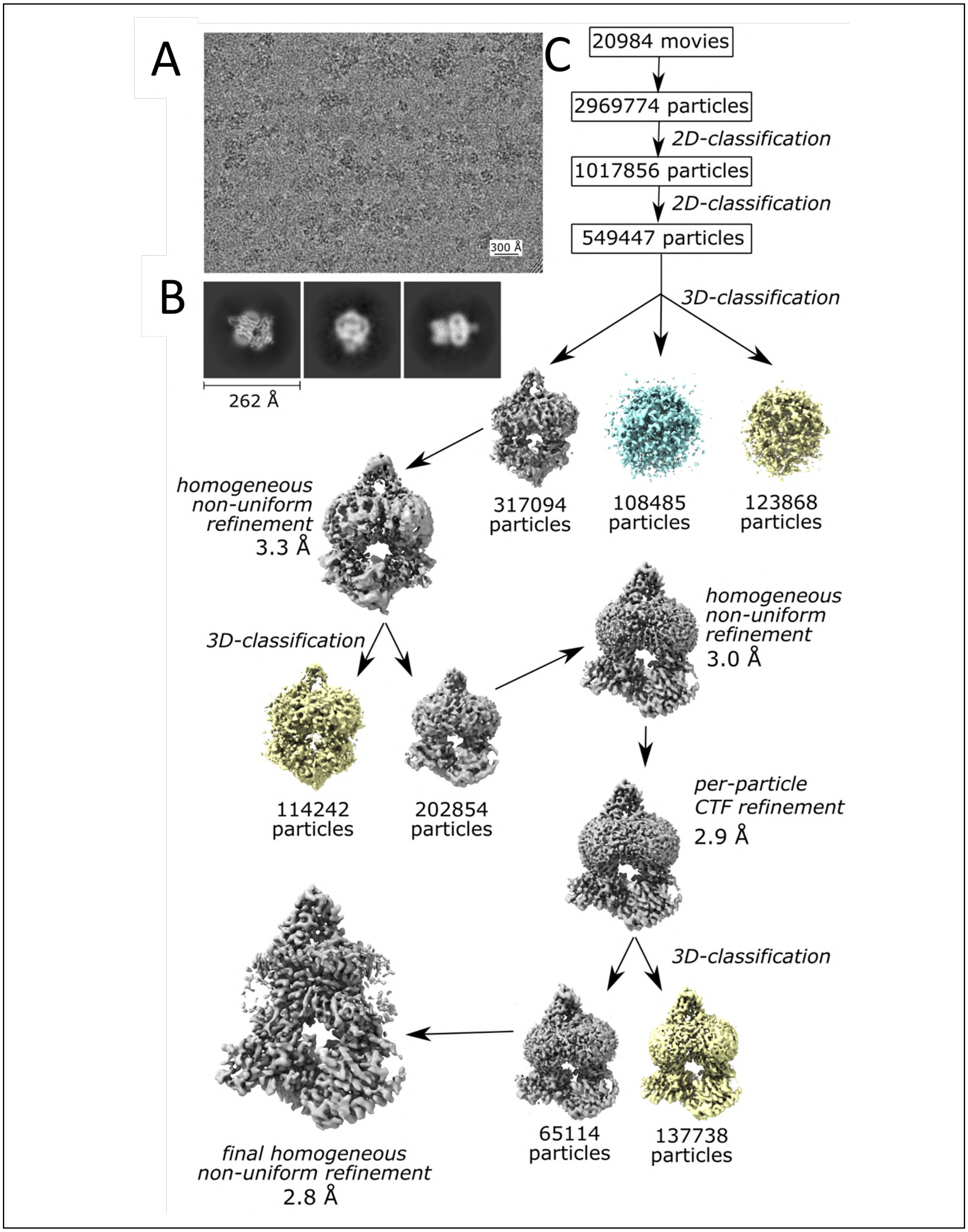

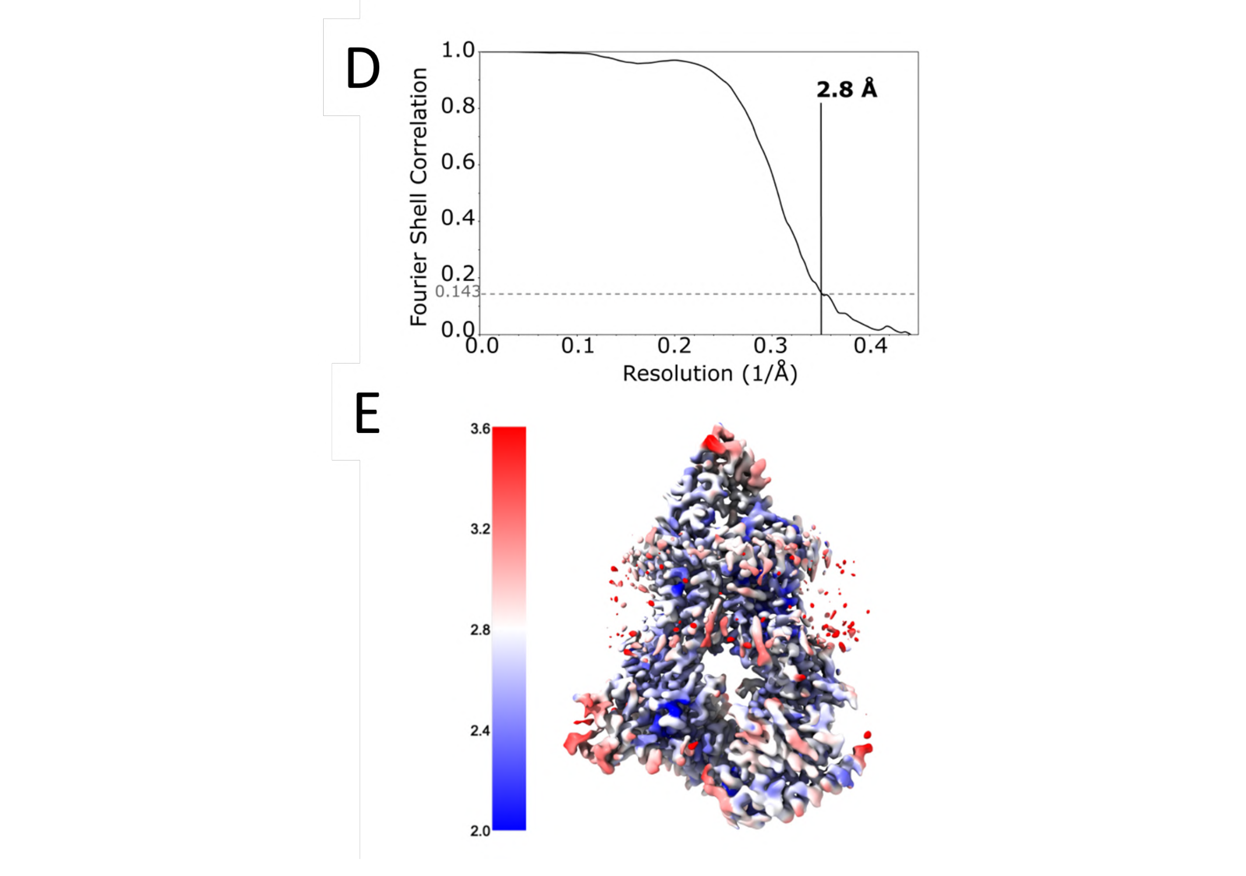

**Supplementary Figure 3. Cryo-EM particle processing workflow for *Cg*Cdr1 apo. A.** Typical micrograph from a single data collection of 20,984 micrographs used for automatic particle picking for the 2D classification. **B**. 2D class averages of *Cg*Cdr1. **C**. Particle processing. After two rounds of 2D-classification, particles were further classified by multiple rounds of 3D classification and homogeneous refinement. The final set of particles was homogeneously refined using the initial reference-free generated models to generate a 2.8 Å density map. The final map was further improved with per-particle CTF refinement, and cryoSPARC's non-uniform homogeneous refinement. **D**. Fourier Shell Correlation (FSC) of the final volume. **E**. Local resolution estimation of the final volume.

**Supplementary Figure 4**

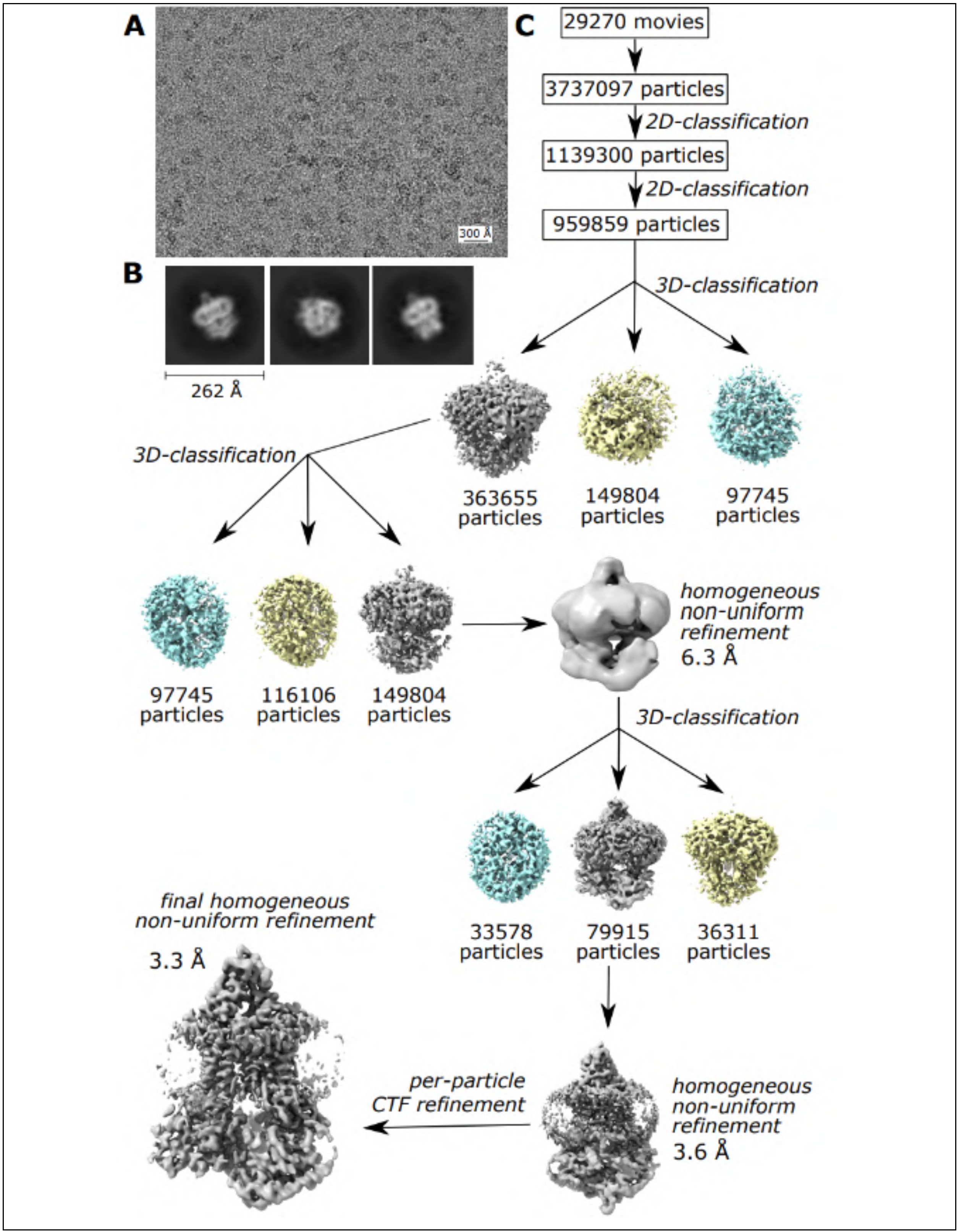

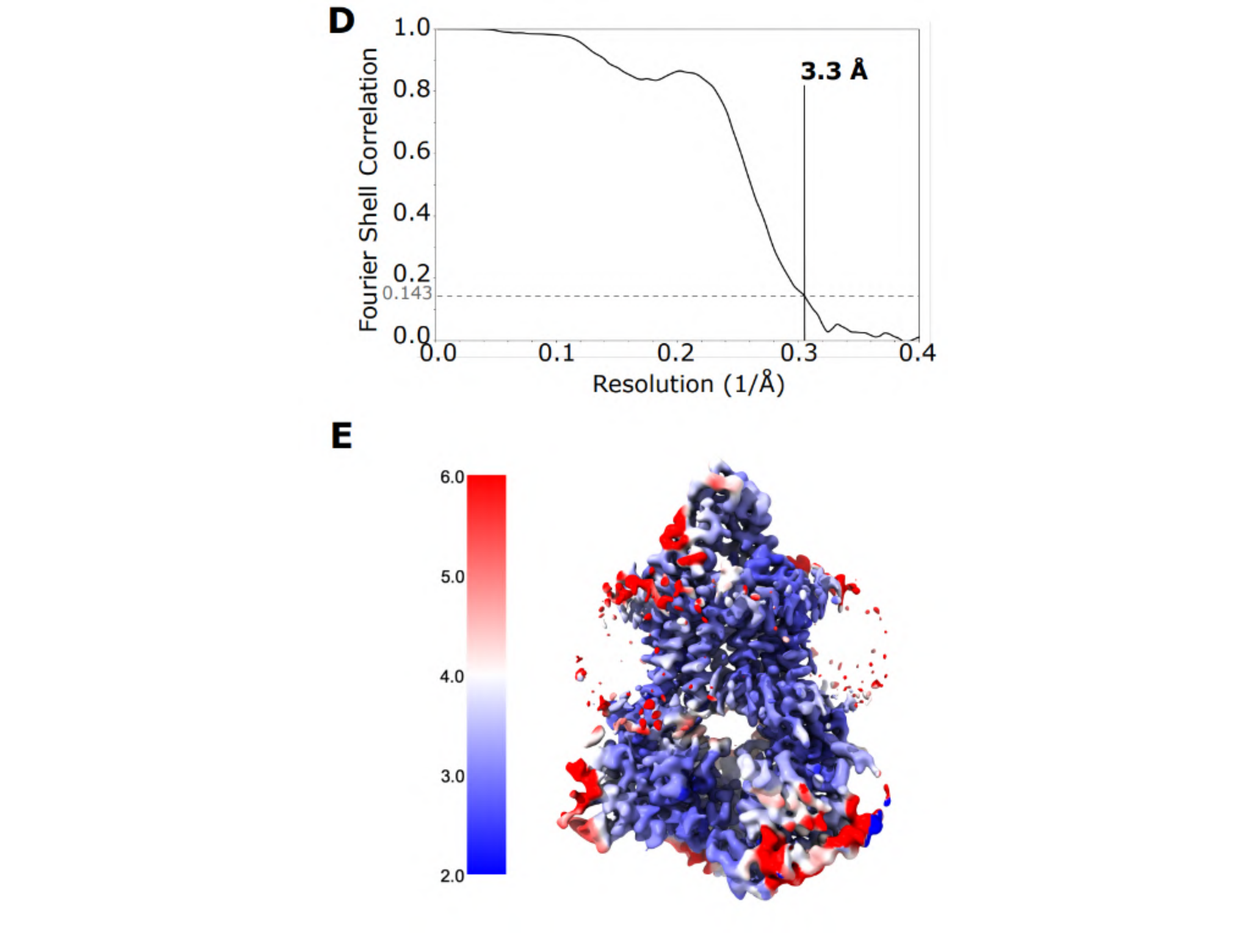

**Supplementary Figure 4. Cryo-EM particle processing workflow for *Cg*Cdr1 incubated with itraconazole, ATP-$Mg^{2+}$ and vanadate. A.** Typical micrograph from a single data collection of 29,270 micrographs used for automatic particle picking for the 2D classification. **B.** 2D class averages of *Cg*Cdr1. **C.** Particle processing. After two rounds of 2D-classification, particles were further classified by multiple rounds of 3D classification and homogeneous refinement. The final set of particles was homogeneously refined using the initial reference-free generated models to generate a 3.3 Å density map. The final map was further improved with per-particle CTF refinement, and cryoSPARC's non-uniform homogeneous refinement. **D**. Fourier Shell Correlation (FSC) of the final volume. **E.** Local resolution estimation of the final volume.

**Supplementary Figure 5**

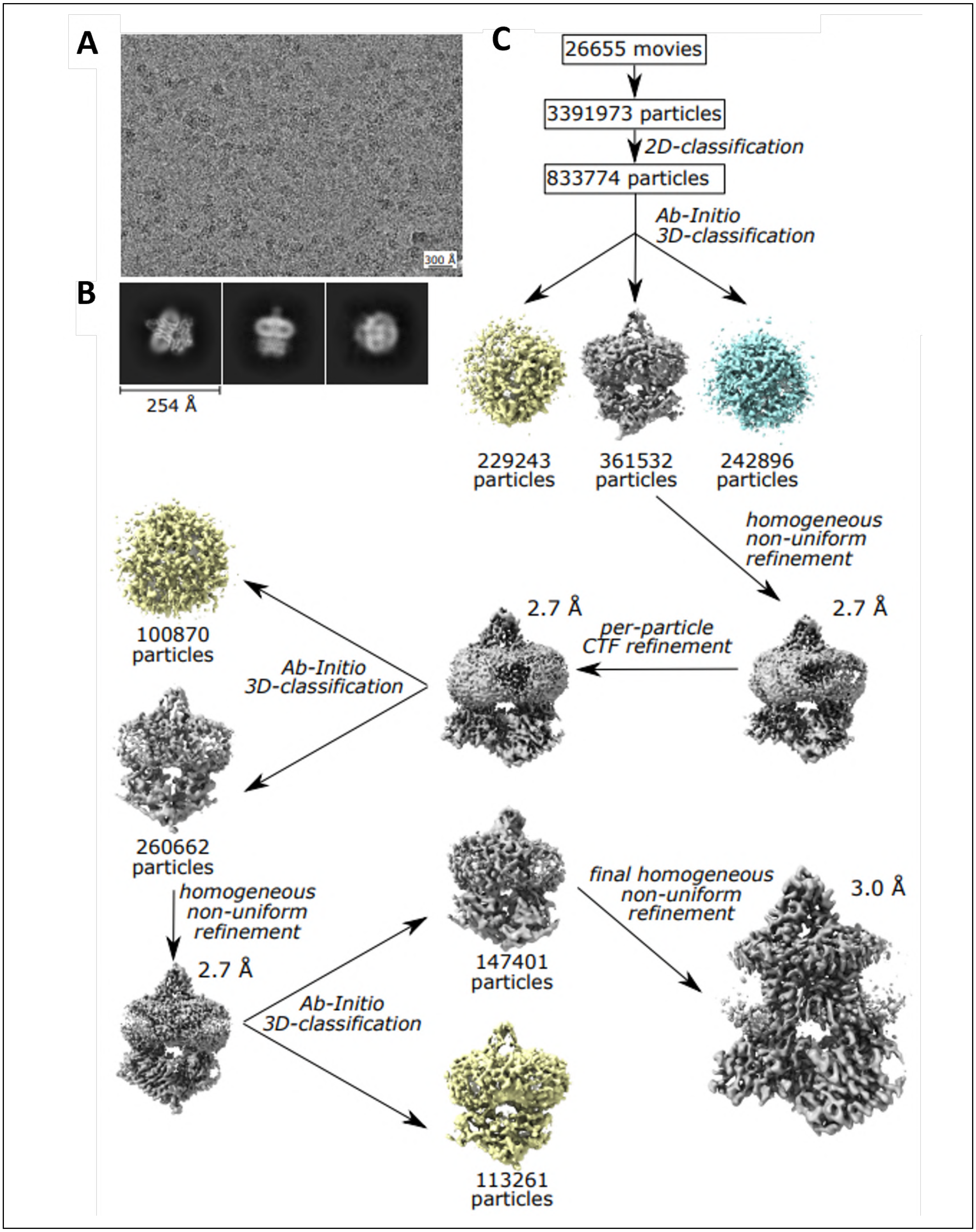

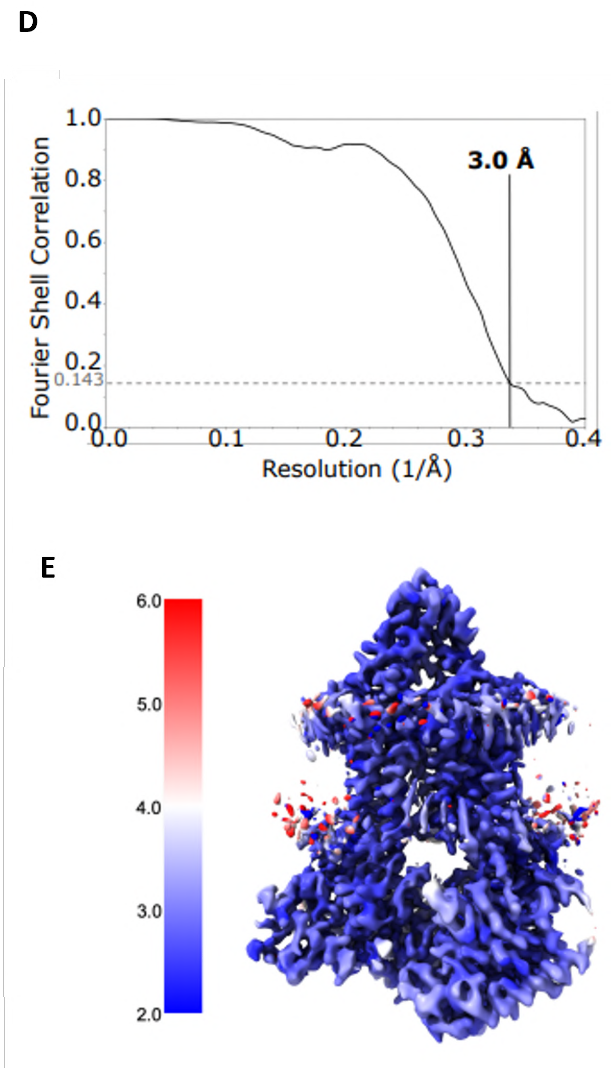

**Supplementary Figure 5. Cryo-EM particle processing workflow for *Cg*Cdr1 incubated with itraconazole and ATP-$Mg^{2+}$. A.** Typical micrograph from a single data collection of 29,270 micrographs used for automatic particle picking for the 2D classification. **B.** 2D class averages of *Cg*Cdr1. **C.** Particle processing. After two rounds of 2D-classification, particles were further classified by multiple rounds of 3D classification and homogeneous refinement. The final set of particles was homogeneously refined using the initial reference-free generated models to generate a 2.7 Å density map. The final map was further improved with per-particle CTF refinement, and cryoSPARC's non-uniform homogeneous refinement. **D**. Fourier Shell Correlation (FSC) of the final volume. **E.** Local resolution estimation of the final volume.

**Supplementary Figure 6**

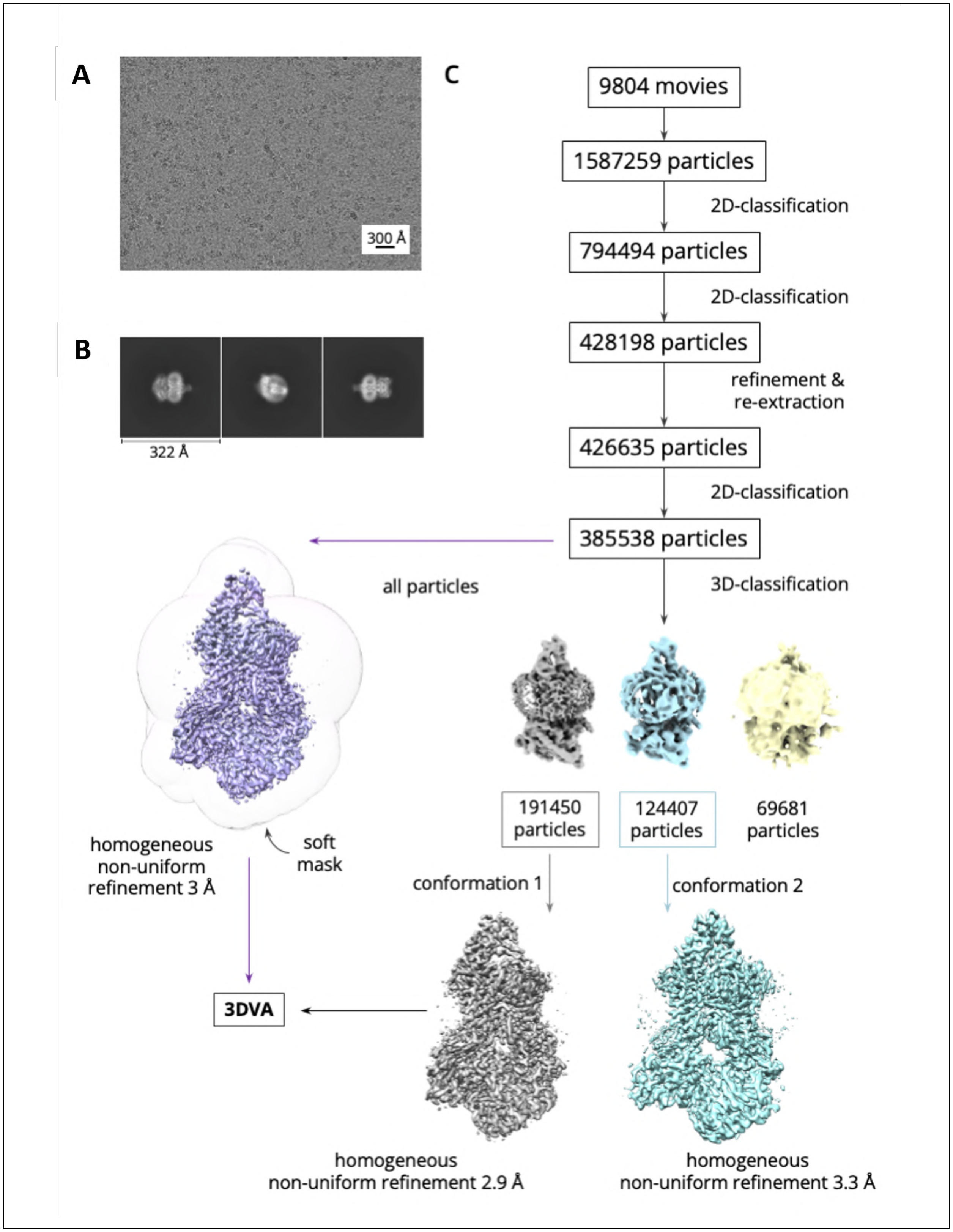

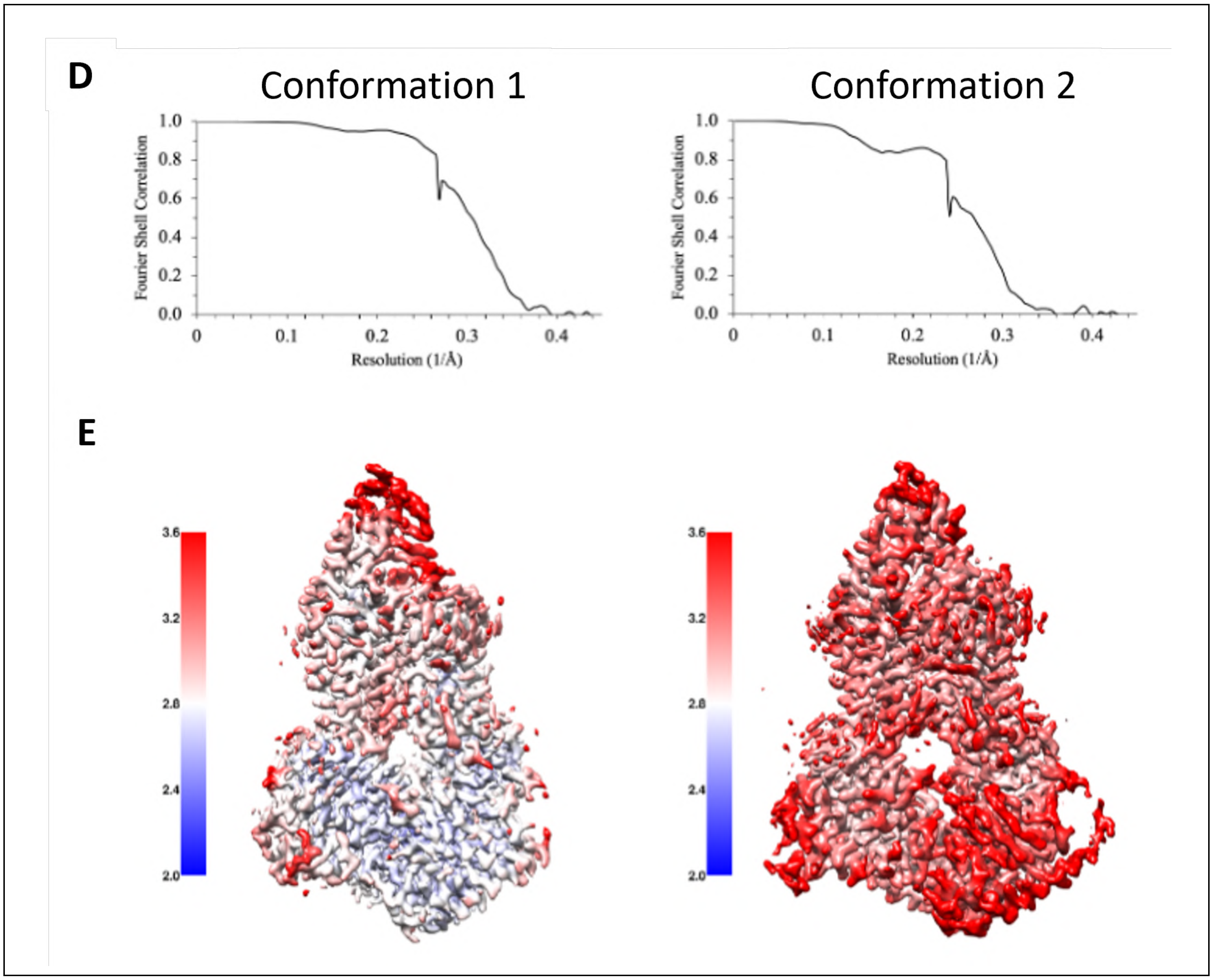

**Supplementary Figure 6. Cryo-EM particle processing workflow for *Cg*Cdr1 incubated with ATP-$Mg^{2+}$ and vanadate. A.** Typical micrograph from a dataset of 9,084 movies that shows *Cg*Cdr1 particles embedded in vitreous ice. crYOLO-picked particles were used for the 2D classifications. **B.** Selected 2D class averages of *Cg*Cdr1. **C.** Image processing. After two rounds of 2D-classification, particles were re-extracted, re-centered and submitted for a final round of 2D classification before being further classified in 3D by 3 classes of *ab initio* reference-free generated models and subsequent heterogeneous refinement with the 3 generated volumes. Two of the classes showed distinct conformations and their respective particles were refined separately by non-uniform homogeneous refinement. Alternatively, all particles of the final subset were used for a consensus non-uniform homogeneous refinement. **D.** Fourier Shell Correlation (FSC). **E.** Local-resolution estimation of the final volumes for both conformations.

### Supplementary Figure 7

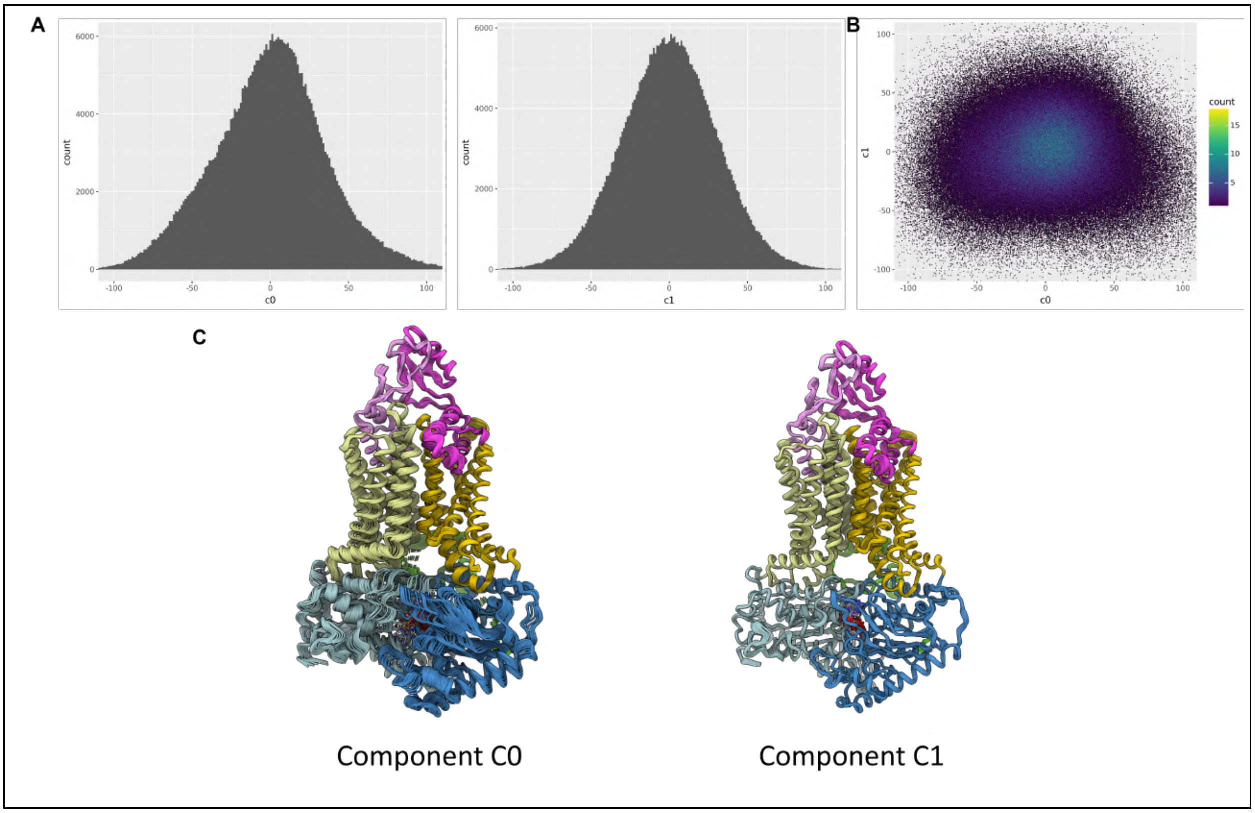

**Supplementary Figure 7. Particle distribution in latent space for the transition from the $[^{s}0|^{nc}ATP|^{c}ADP\text{-}Vi\text{-}Mg]$ state to the $[^{s}0|^{nc}ATP|^{c}ADP\text{-}Mg]$ state . A**. Number of particles as a function of latent coordinate, represented by principal and secondary variability components C0 and C1, respectively. **B**. Same as panel A but represented in 2 dimensions. Each dot represents a particle that has been plotted according to its latent coordinate in the 2 main components C0 and C1, as listed on the x- and y-axes of the graph. The color gradient represents the number of particles in each bin according to the scale on the right of the image. **C**. 3D models generated by 3DVA and VARREF analyses for each component.

**Supplementary Figure 8**

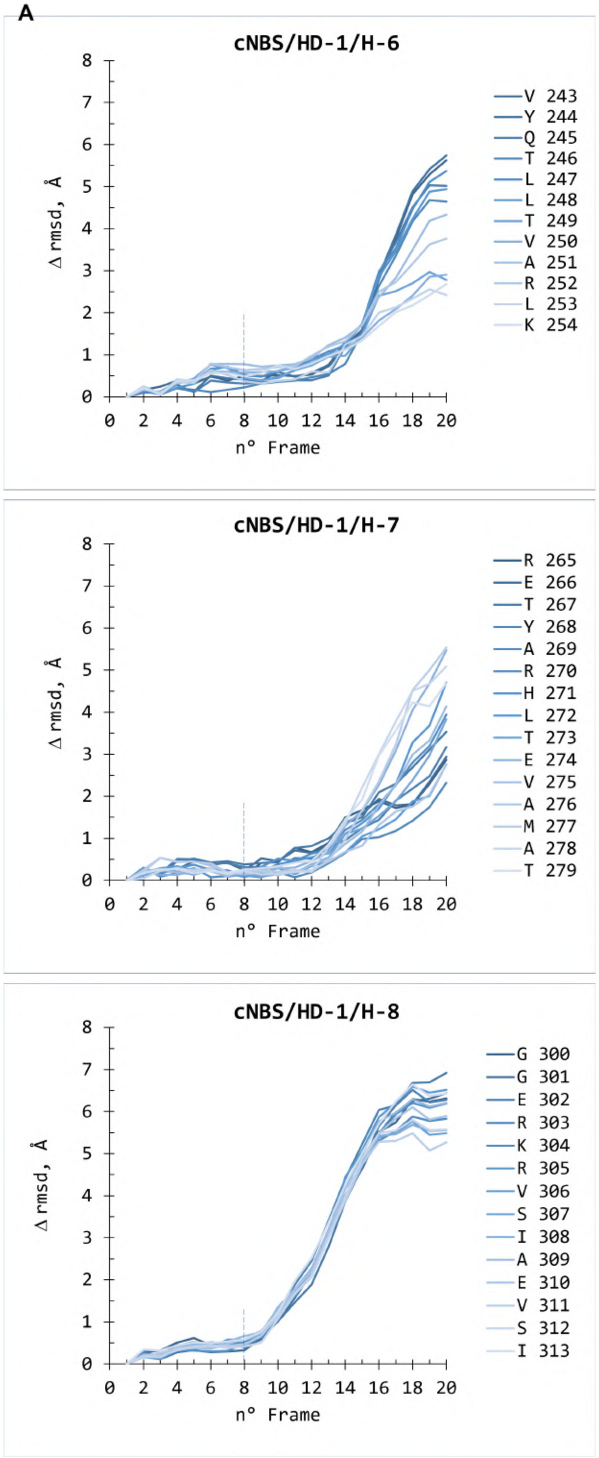

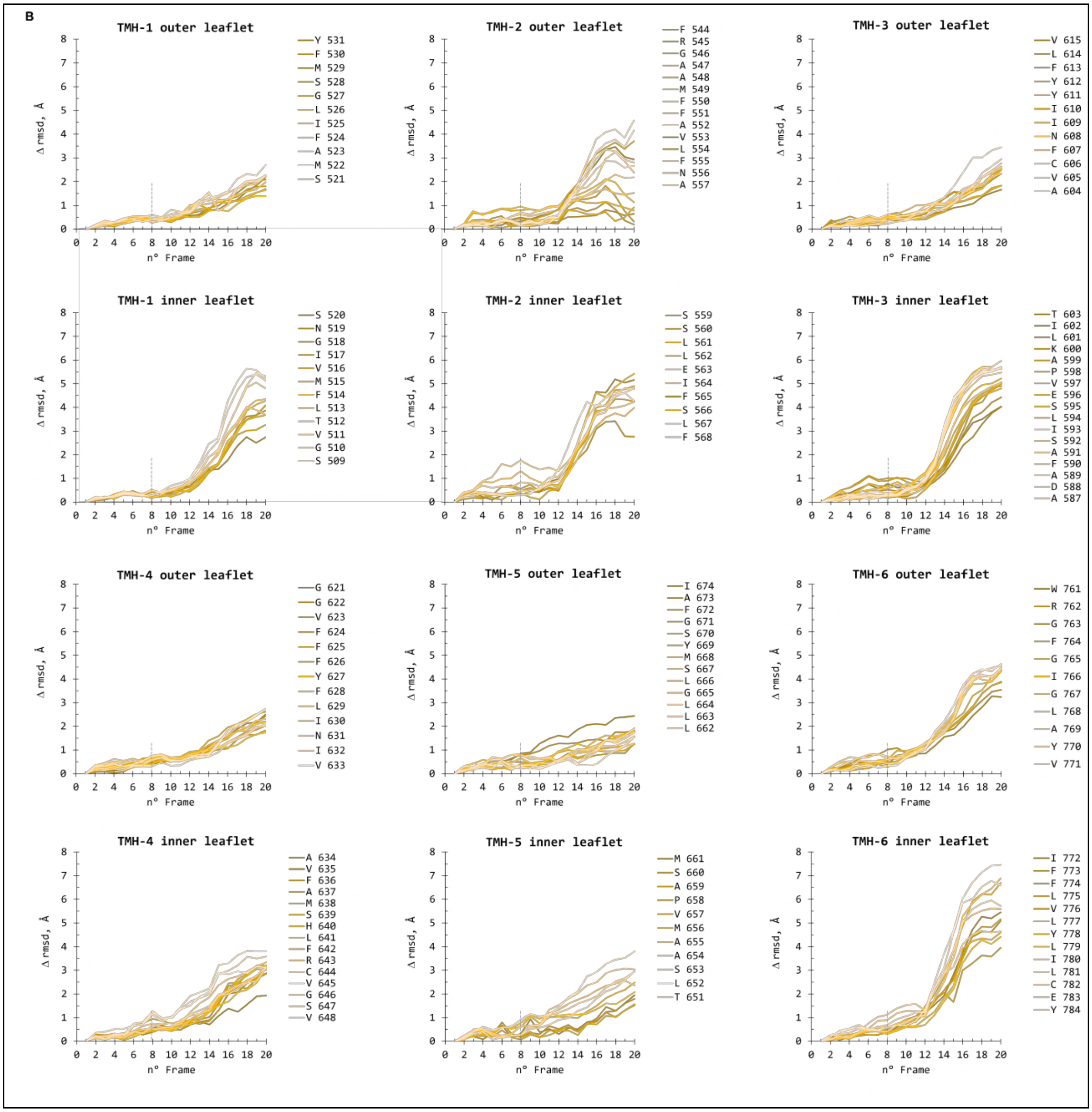
B
TMH-1 outer leaflet
TMH-2 outer leaflet
TMH-3 outer leaflet
TMH-1 inner leaflet
TMH-2 inner leaflet
TMH-3 inner leaflet
TMH-4 outer leaflet
TMH-5 outer leaflet
TMH-6 outer leaflet
TMH-4 inner leaflet
TMH-5 inner leaflet
TMH-6 inner leaflet
Δ rmsd, Å
n° Frame
Y 531
F 530
M 529
S 528
G 527
L 526
I 525
F 524
A 523
M 522
S 521
F 544
R 545
G 546
A 547
A 548
M 549
F 550
F 551
A 552
V 553
L 554
F 555
N 556
A 557
V 615
L 614
F 613
Y 612
Y 611
I 610
I 609
N 608
F 607
C 606
V 605
A 604
S 520
N 519
G 518
I 517
V 516
M 515
F 514
L 513
T 512
V 511
G 510
S 509
S 559
S 560
L 561
L 562
E 563
I 564
F 565
S 566
L 567
F 568
T 603
I 602
L 601
K 600
A 599
P 598
V 597
E 596
S 595
L 594
I 593
S 592
A 591
F 590
A 589
D 588
A 587
G 621
G 622
V 623
F 624
F 625
F 626
Y 627
F 628
L 629
I 630
N 631
I 632
V 633
I 674
A 673
F 672
G 671
S 670
Y 669
M 668
S 667
L 666
G 665
L 664
L 663
L 662
W 761
R 762
G 763
F 764
G 765
I 766
G 767
L 768
A 769
Y 770
V 771
A 634
V 635
F 636
A 637
M 638
S 639
H 640
L 641
F 642
R 643
C 644
V 645
G 646
S 647
V 648
M 661
S 660
A 659
P 658
V 657
M 656
A 655
A 654
S 653
L 652
T 651
I 772
F 773
F 774
L 775
V 776
L 777
Y 778
L 779
I 780
L 781
C 782
E 783
Y 784

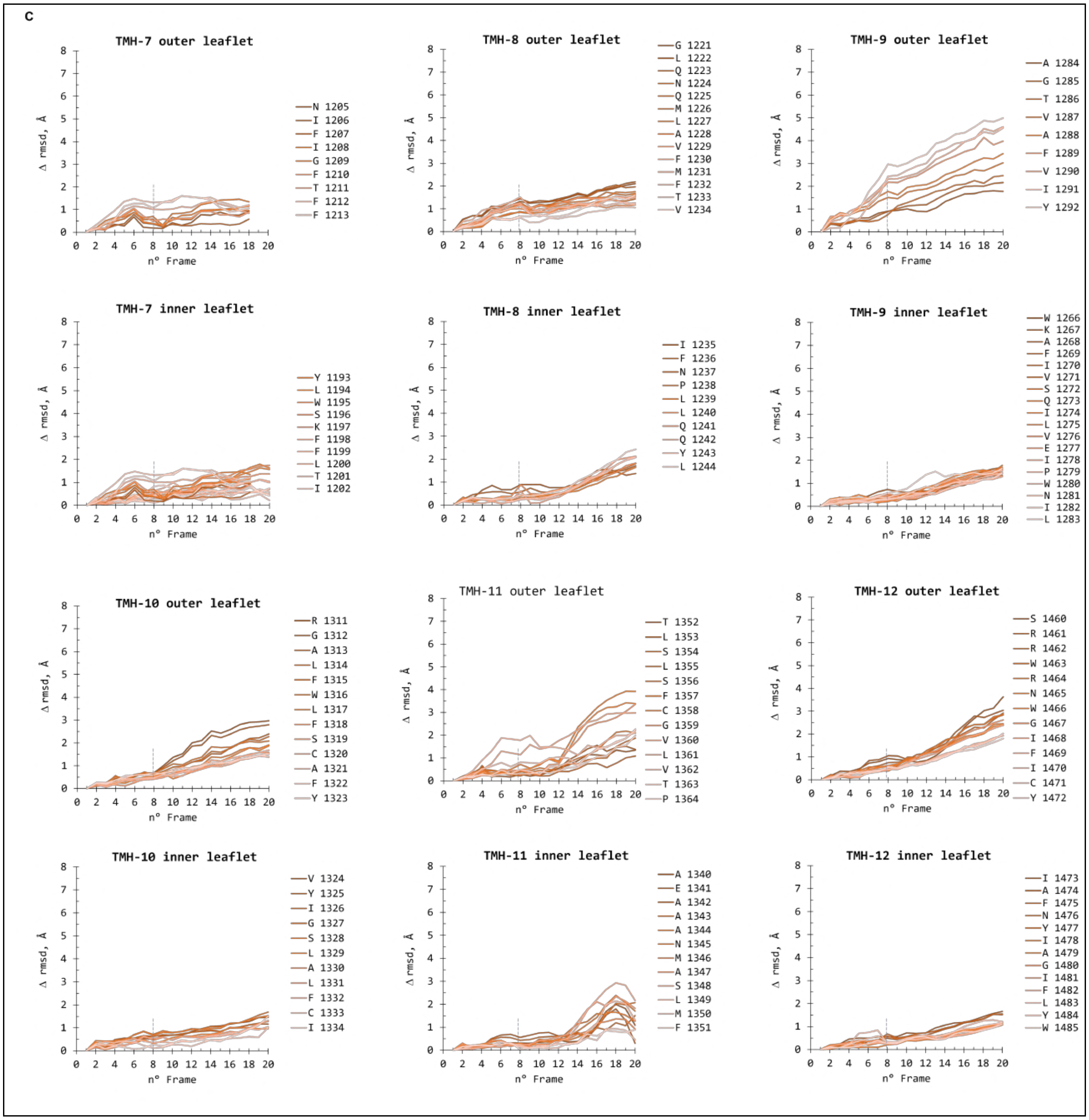

**Supplementary Figure 8. RMSD variation for each residue of the HD-1 and TMD helices along the 20 frames of component C0 particles representing the transition from the closed [s0|ncATP|cADP-Vi-Mg] to the open [s0|ncATP|cADP-Mg] states of *Cg*Cdr1. A.** HD-1 helices. **B.** TMD-1 helices. **C.** TMD-2 helices. The variation of RMSD is calculated for frames 2 to 20 in respect of frame 1 as detailed in Supplementary data 1.

### Supplementary Figure 9

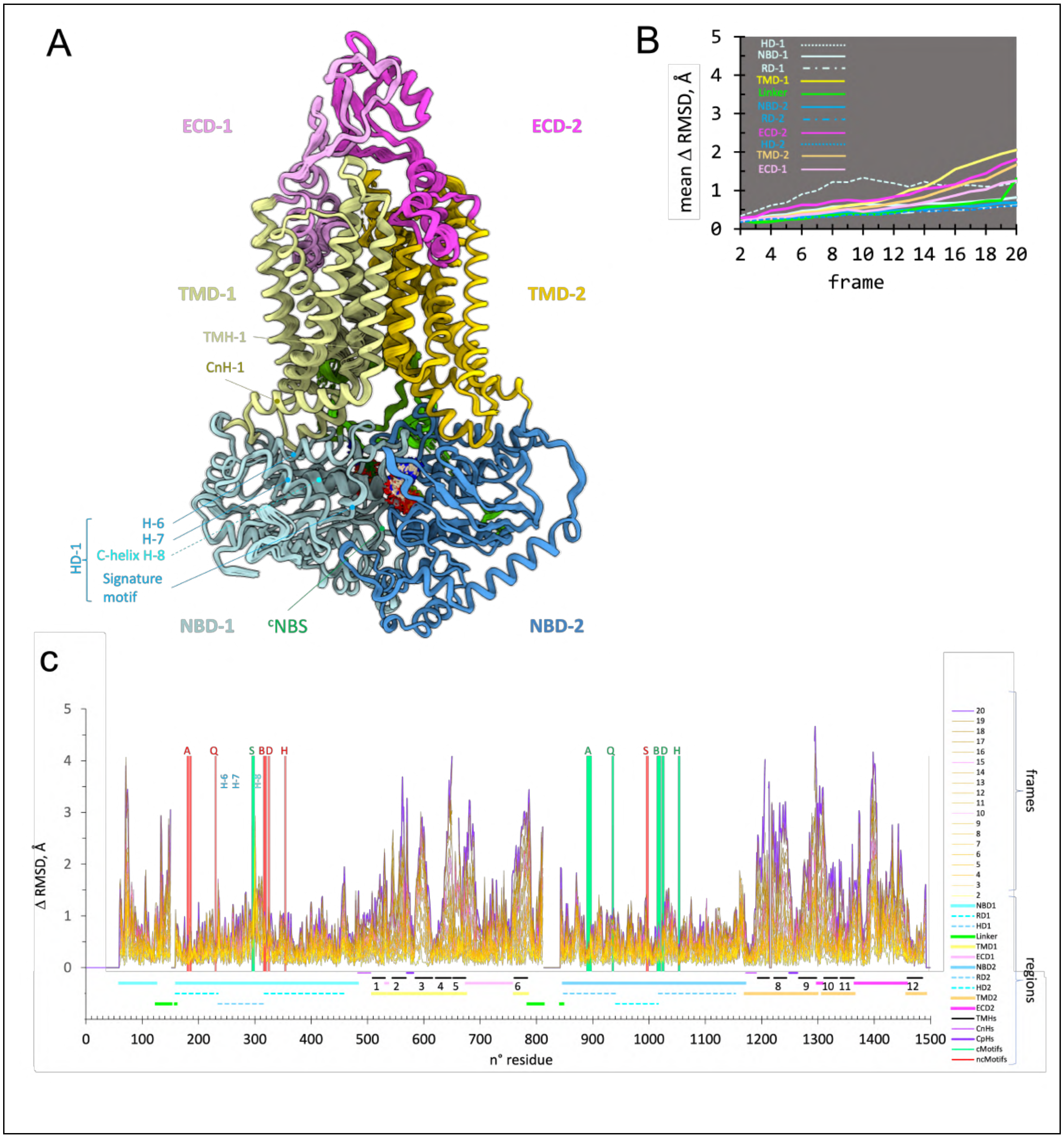

**Supplementary Figure 9. Conformational variability analysis of the transition from the closed [$^{s}$0|$^{nc}$ATP|$^{c}$ADP-Vi-Mg] to the open [$^{s}$0|$^{nc}$ATP|$^{c}$ADP-Mg] states of *Cg*Cdr1 of component C1 particles.** The legend of panels A-C is the same as that for Figure 3.

### Supplementary Figure 10

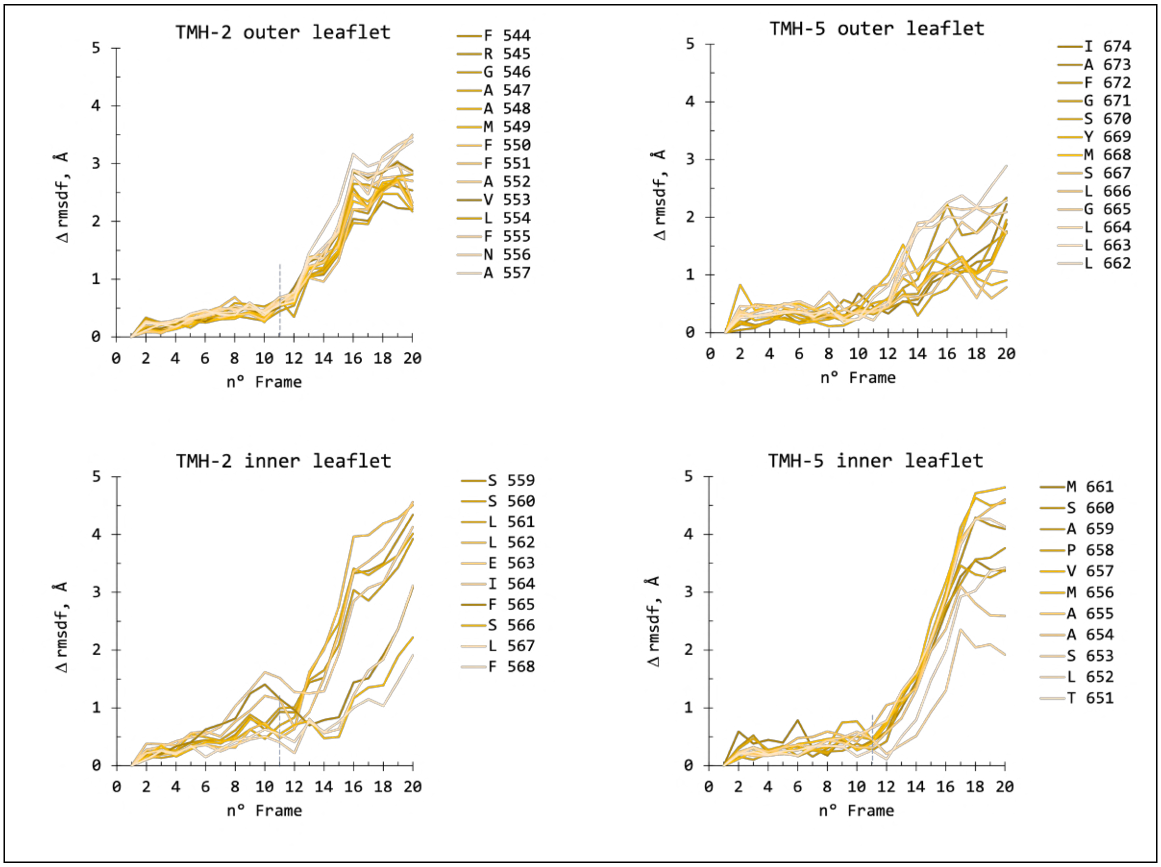

**Supplementary Figure 10. RMSD variation of each residue of TMH-2 and -5 along the 20 component C1 frames from the closed [$^{s}$0|$^{nc}$ATP|$^{c}$ADP-Vi-Mg] to the open [$^{s}$0|$^{nc}$ATP|$^{c}$ADP-Mg] states of *Cg*Cdr1.** Legend is as for Supplementary Figure 8. The variation of RMSD is calculated for frames 2 to 20 in respect of frame 1 as detailed in Supplementary data 2.

### Supplementary Figure 11

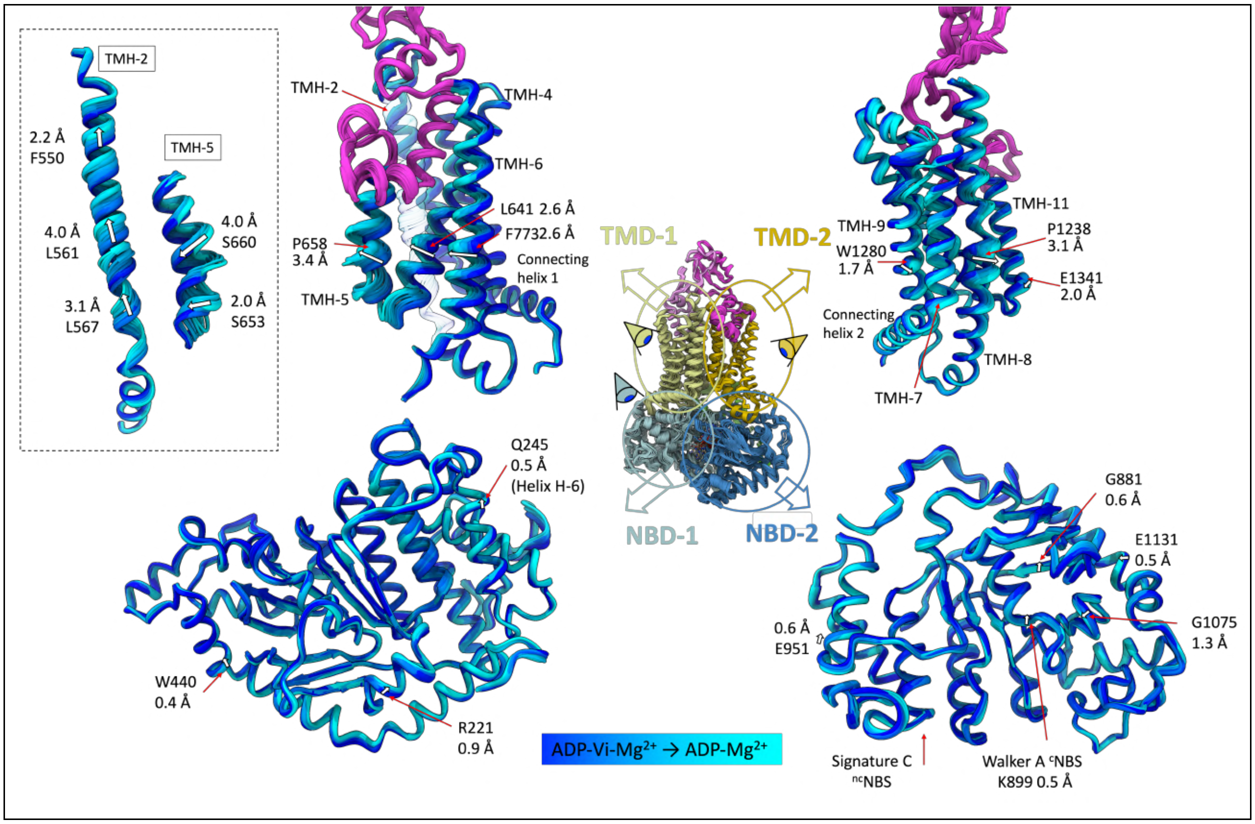

**Supplementary Figure 11. Conformational transition from the closed [$^{s}$0|$^{nc}$ATP|$^{c}$ADP-Vi-Mg] to the open [$^{s}$0|$^{nc}$ATP|$^{c}$ADP-Mg] states of component C1 particle structures highlighting the 4 major domains of *Cg*Cdr1.** The structure is shown as a cartoon colored from blue to cyan to depict the transition across the different models. TMH-2 and TMH-5 are transparent in the whole TMD-1 image and presented independently in the inset to the left, as their movement is different from that of the rest of TMD-1, showing a simultaneous constriction of TMH-5 and an upward movement of TMH-2. This movement closes the DBS cavity from the cytoplasmic side and would move the substrate upwards towards the extracellular space. Movements are listed for each domain with white arrows showing the directionality and the amplitude is given.

### Supplementary Figure 12

| | | | | | | | | | | | |
|---|---|---|---|---|---|---|---|---|---|---|---|
| | Ergosterol, PE and PC lipides quantitation by HPLC in Cdr1/PCC-DCOD9b sample | | | | | | | | | | |
| | **Ergosterol** | Protein | | | sample | Ergosterol | | | | | **erg/Cdr1** |
| | | g/L | mol/L | MW, g/mol | µL assay | µg Erg added | µg Erg quantified | g/L | MW, g/mol | mol/L | **mol/mol** |
| | Cdr1 | 0.9 | 5.3E-06 | 170000 | 450 | 0 | 10.8 | 2.4E-02 | 397 | 6.0E-05 | **1.1E+01** |
| | Buffer | | | | 1000 | 5 | 4.5 | 4.5E-03 | 397 | 1.1E-05 | |
| **Samples** | | | | | | | | | | | |
| | **PE** | 16:0 16:0 | 16:0 16:1 | 16:0 18:0 | 16:0 18:1 | 16:1 16:1 | 16:1 18:0 | **16:1 18:1** | 18:0 18:1 | 18:1 18:1 | **total** |
| | mol/L | 3.6E-08 | 1.40E-06 | 2.73E-08 | 9.49E-07 | 5.72E-07 | 1.28E-07 | **2.61E-06** | 4.84E-08 | 1.60E-07 | **mol/mol** |
| | mol/mol Cdr1 | 6.8E-03 | 2.6E-01 | 5.1E-03 | 1.8E-01 | 1.1E-01 | 2.4E-02 | **4.9E-01** | 9.1E-03 | 3.0E-02 | **1.1E+00** |
| | **PC** | 16:0 16:1 | 16:0 18:0 | 16:0 18:1 | **16:1 16:1** | 16:1 18:0 | **16:1 18:1** | 18:0 18:1 | 18:1 18:1 | | **total** |
| | mol/L | 5.79E-07 | 3.68E-07 | 3.13E-07 | **1.83E-06** | 2.25E-07 | **1.80E-06** | 2.01E-07 | 2.39E-07 | | **mol/mol** |
| | mol/mol Cdr1 | 1.1E-01 | 7.0E-02 | 5.9E-02 | **3.5E-01** | 4.3E-02 | **3.4E-01** | 3.8E-02 | 4.5E-02 | | **1.1E+00** |

**Supplementary Figure 12. HPLC quantification of ergosterol, PC and PE species bound to *Cg*Cdr1 in the PCC-DCOD9b complex.** Lipids were quantified as described in Supplementary Methods.

### Supplementary Figure 13

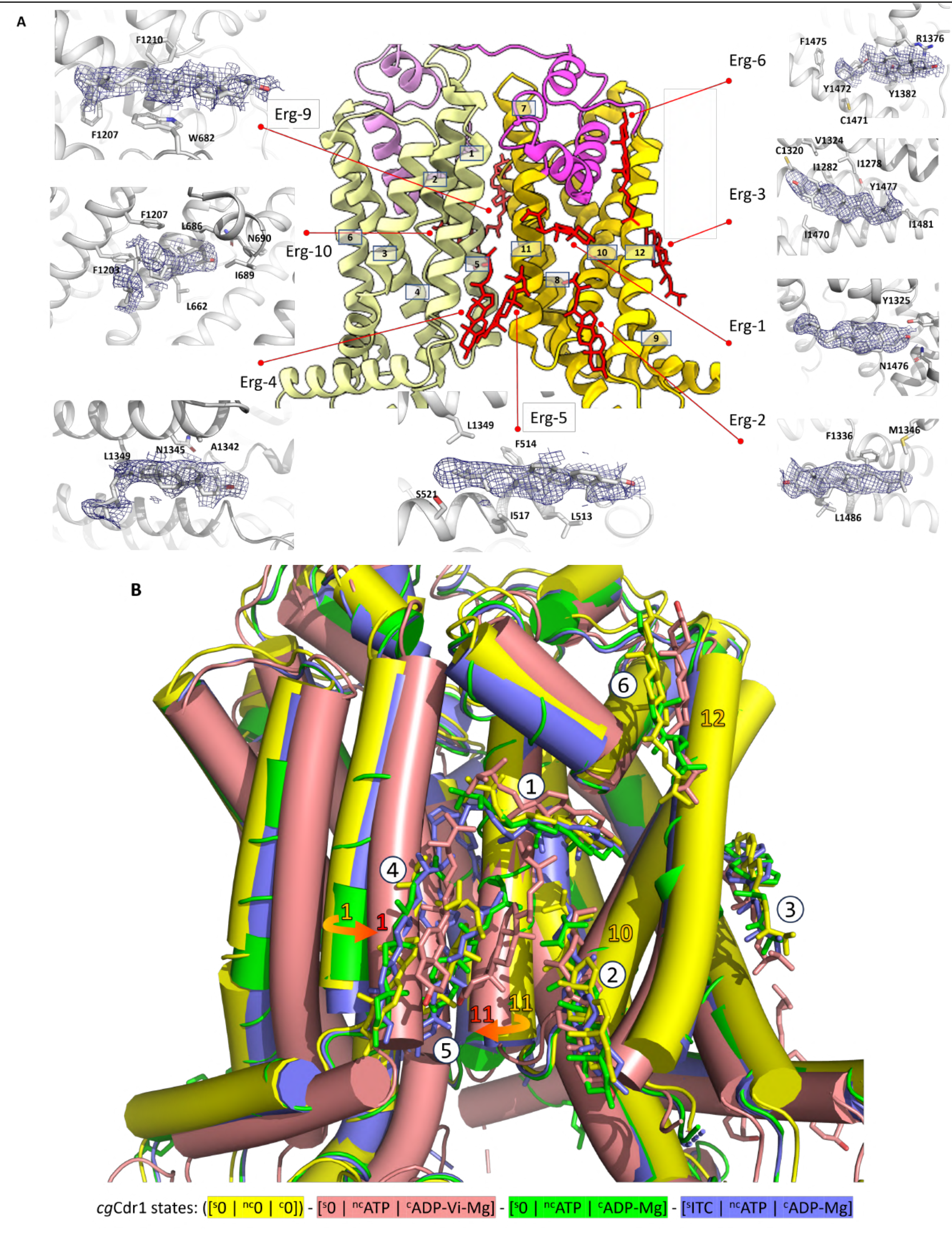

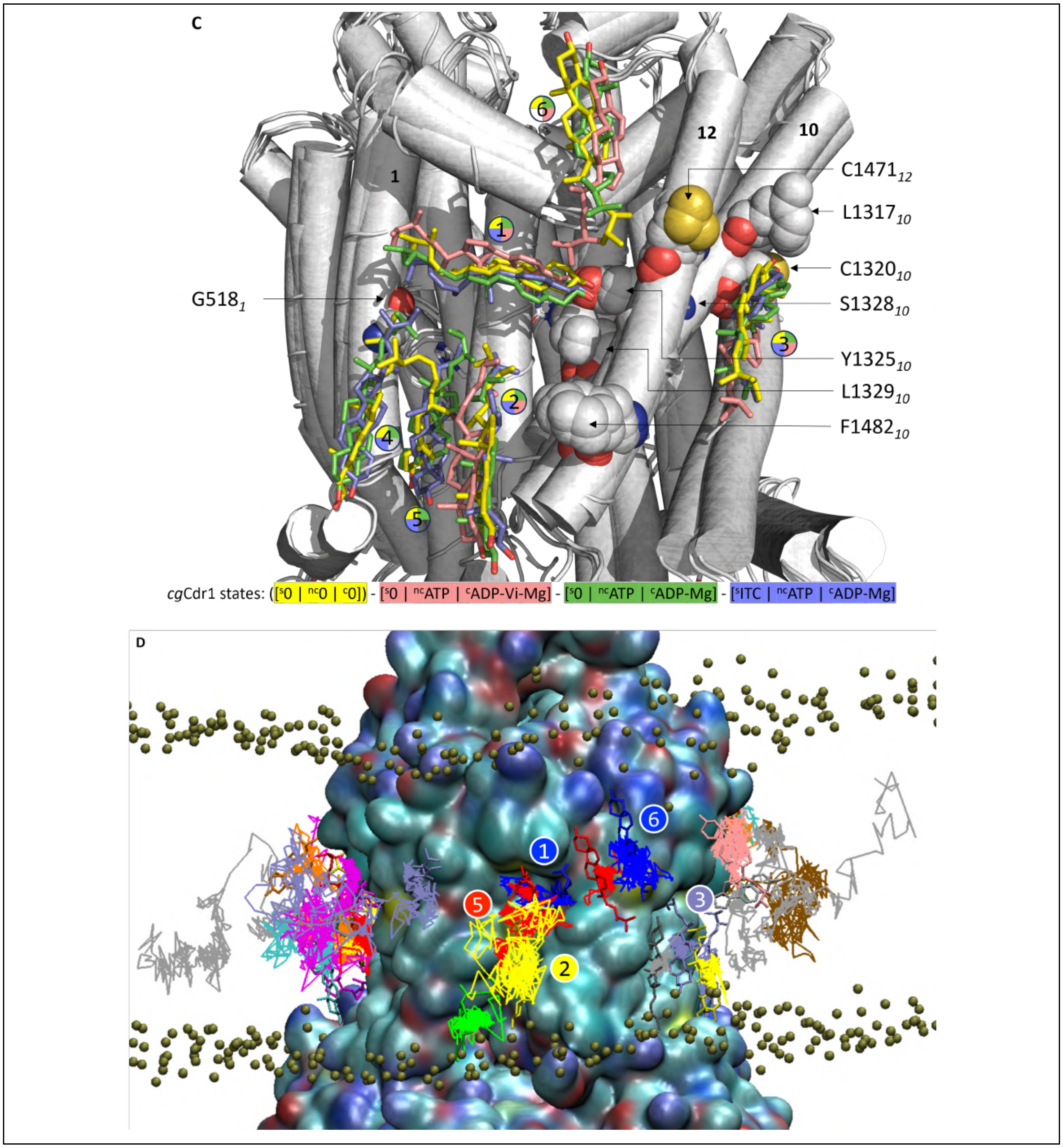

**Supplementary Figure 13. Distribution of ergosterol molecules bound to the TMDs of *Cg*Cdr1. A.** Positions and densities compatible with ergosterol molecules in the Apo state. **B** Superposition of the ergosterol molecules identified in the three different $^{c}$NBS states of CgCdr1 in the presence or absence of itraconazole in the DBS. **C.** Residues interacting with ergosterol molecules that have been shown to be critically important for steroid hormone transport in *Ca*Cdr1. **D.** *In silico* evaluation of the mobility of bound ergosterol molecules. Molecular dynamics simulation of the *Cg*Cdr1 [$^{nc}$ATP|$^{c}$ADP-Mg] state with bound ergosterol molecules embedded into a lipid membrane containing 20 % ergosterol, 20 %

DOPG and 60 % DOPE. Traces show the mobility of each ergosterol molecule in the simulation triplicate with the structure of their destination displayed in the same color.

### Supplementary Figure 14

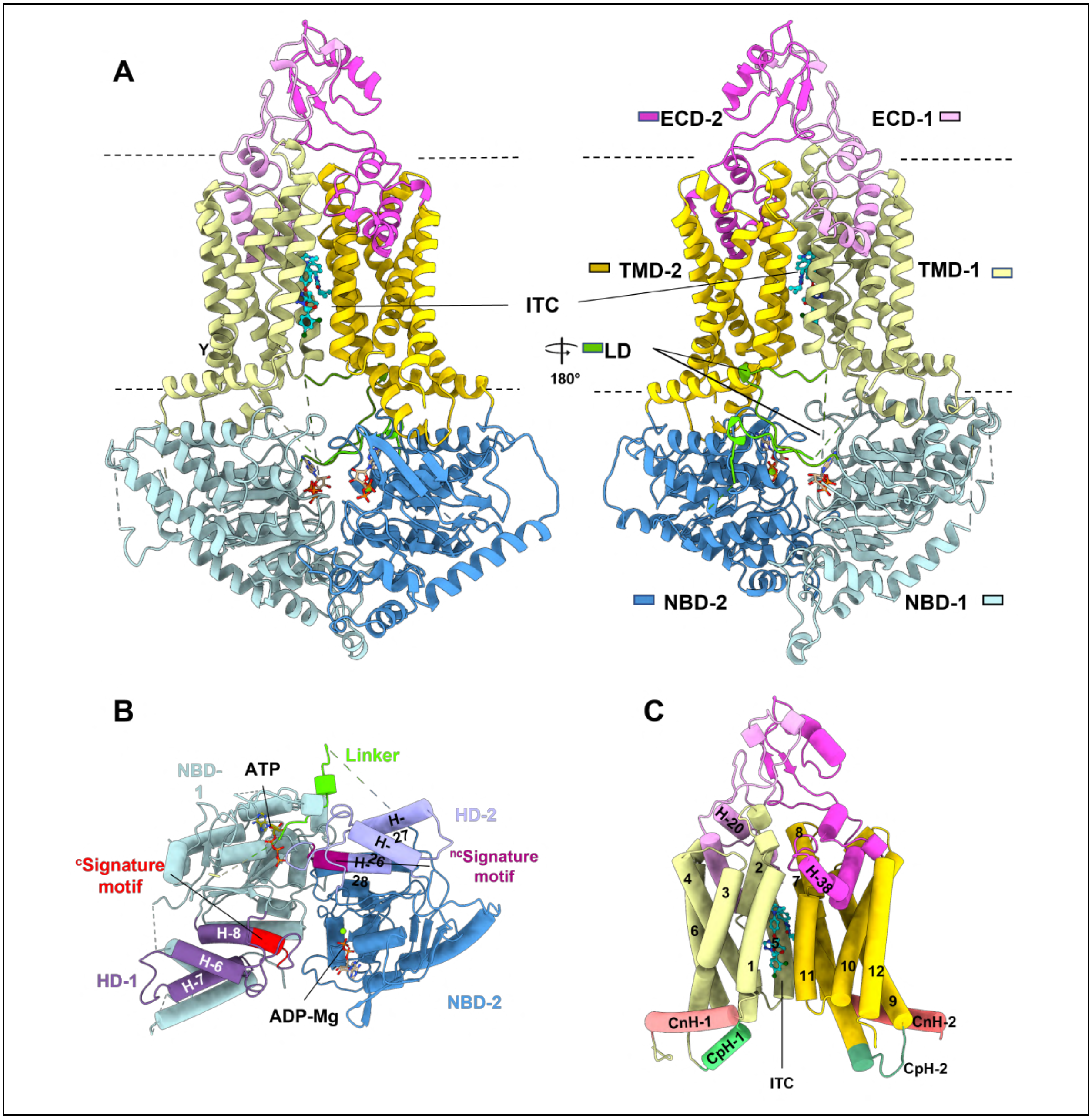

**Supplementary Figure 14. Overall and detailed views of the cryo-EM structure of *Cg*Cdr1 [$^{s}$ITC|$^{nc}$ATP|$^{c}$ADP-Mg. A:** Longitudinal views from the $^{c}$NBS (left) and $^{nc}$NBS (right). **B.** NBDs viewed from the cytoplasm. **C.** TMD viewed in the plane of the membrane. *Cg*Cdr1 is shown as a cartoon, and nucleotides and itraconazole are shown as sticks. Each domain is colored according to Figure 2 and secondary structures are numbered as in Supplementary Figure 1C.

**Supplementary Figure 15**

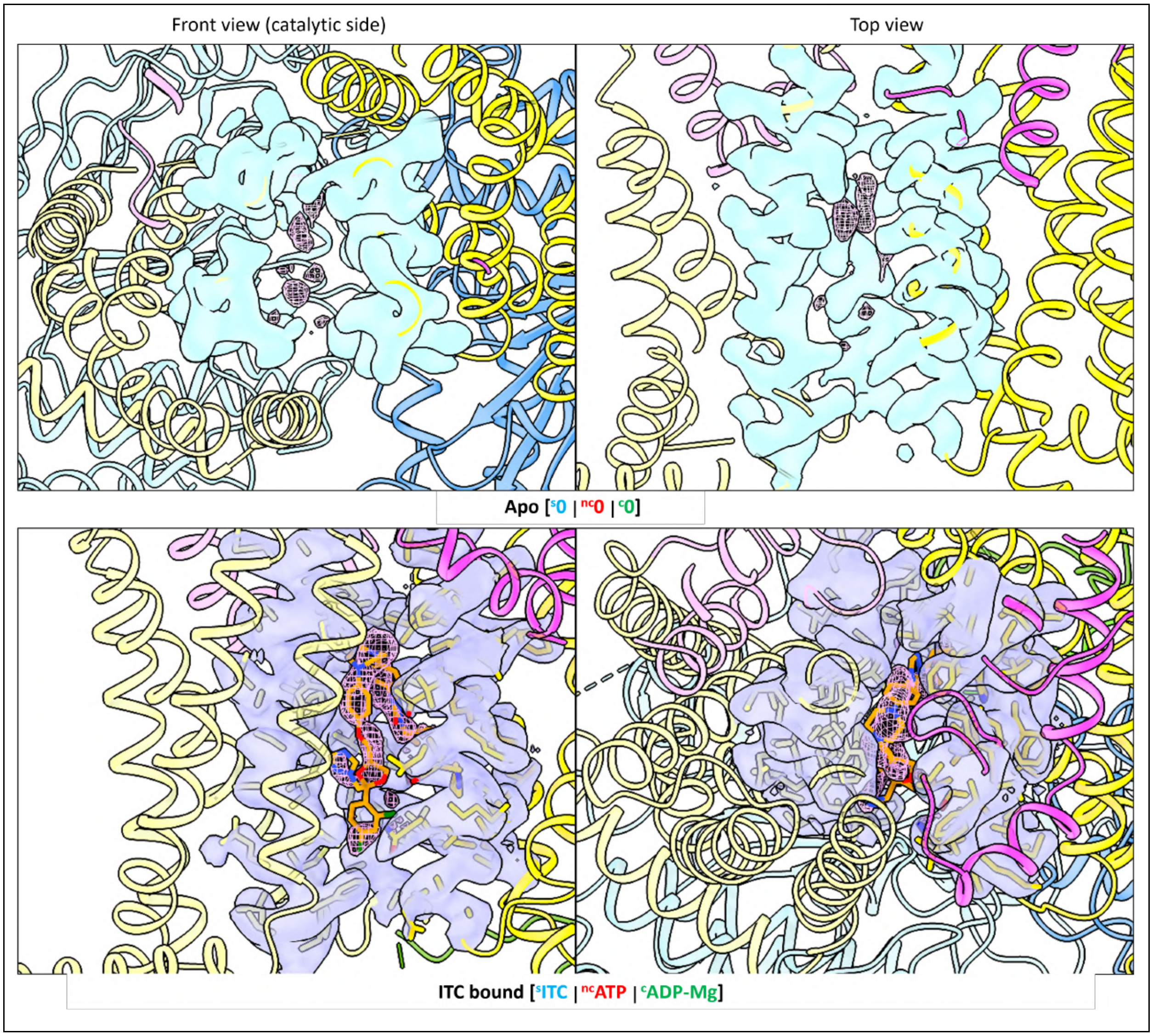

**Supplementary Figure 15. Front and top view of the DBS of the apo state and the ITC-bound state of *Cg*Cdr1**. Cartoon model of the apo and the ITC-bound state of *Cg*Cdr1, with the domains coloured as in figure 1D. Densities of residues of TMH2, 5, 8, 11 are shown in cyan (upper row, apo) and lilac (lower row, ITC-bound state). The contents of the DBS are shown at the same contour, in pink as a mesh. The ITC molecule is shown in orange as sticks. Contour $\sigma_{(apo)}$= 0.114, Contour $\sigma_{(ITC\text{-}bound)}$= 0.0499

### Supplementary Table 1. *Cg*Cdr1 cryo-EM data collection, refinement and validation statistics

| **State** | **Apo** | **[$^{s}$ITC$\vert^{nc}$ATP$\vert^{c}$ADP-Mg]** | **[$^{s}$ITC$\vert^{nc}$ATP$\vert^{c}$ADP-Mg-VO$_4$]** | **[$^{s}$0$\vert^{nc}$ATP$\vert^{c}$ADP-Mg-VO$_4$]** | **[$^{s}$0$\vert^{nc}$ATP$\vert^{c}$ADP-Mg]** |
|---|---|---|---|---|---|
| EMD code | 54582 | 54583 | 54581 | 54488 | 54500 |
| PDB code | 9S4T | 9S4U | 9S4S | 9S2C | 9S2H |
| Data collection and processing | | | | | |
| Magnification | 165000× | 165000× | 165000× | 105000× | |
| Voltage (kV) | 300 | 300 | 300 | 300 | |
| Electron exposure ($e^{-}$/Å$^2$) | 40 | 40 | 40 | 40.12 | |
| Defocus range (μm) | 0.6 to 1.8 | 0.6 to 1.8 | 0.6 to 1.8 | 0.5 to 2.5 | |
| Pixel size (Å) | 0.525 | 0.507 | 0.509 | 0.839 | |
| Extraction box size (pixels) | 500 | 500 | 500 | 384 | |
| Symmetry imposed | C1 | C1 | C1 | C1 | C1 |
| Initial particle images (no.) | 2969774 | 3391973 | 3737097 | 1587259 | |
| Final particle images (no.) | 65114 | 147401 | 79915 | 191450 | 124407 |
| Map resolution (Å) | 2.8 | 3.0 | 3.3 | 2.9 | 3.3 |
| FSC threshold | 0.143 | 0.143 | 0.143 | 0.143 | 0.143 |
| Map resolution range (Å) | 5.6 to 1.8 | 9.2 to 1.5 | 12 to 1.9 | 18 to 1.8 | 26 to 1.8 |
| Refinement | | | | | |
| Initial model used (PDB code) | [$^{s}$0$\vert^{nc}$ATP$\vert^{c}$ADP-Mg] | [$^{s}$0$\vert^{nc}$ATP$\vert^{c}$ADP-Mg] | [$^{s}$0$\vert^{nc}$ATP$\vert^{c}$ADP-Mg] | AlphaFold model | [$^{s}$0$\vert^{nc}$ATP$\vert^{c}$ADP-Mg-VO$_4$] |
| Model resolution (Å) | | | | | |
| FSC threshold | 0.5 | 0.5 | 0.5 | 0.5 | 0.5 |
| Resolution (Å) | 3.1 | 3.2 | 3.4 | 3.0 | 3.3 |
| Model composition | | | | | |
| Non-hydrogen atoms (no.) | | | | | |
| Protein residues (no.) | 10851 | 10909 | 11157 | 11410 | 11263 |
| Ligands (no.) | 1313 | 1320 | 1335 | 1388 | 1358 |
| | 11 | 12 | 16 | 12 | 14 |
| *B* factors (Å$^2$) | | | | | |
| Protein | 68.87 | 38.19 | 100.79 | 49.99 | 136.4 |
| Ligand | 92.27 | 56.94 | 119.06 | 72.01 | 152.56 |
| R.m.s. deviations | | | | | |
| Bond lengths (Å) | 0.002 | 0.003 | 0.002 | 0.005 | 0.003 |
| Bond angles (°) | 0.513 | 0.698 | 0.554 | 0.670 | 0.602 |
| Validation | | | | | |
| MolProbity score | 1.73 | 2.15 | 1.96 | 2.07 | 2.1 |
| Clashscore | 3.72 | 8.19 | 8.58 | 8.77 | 8.2 |
| Poor rotamers (%) | 2.73 | 2.9 | 1.56 | 2.34 | 2.05 |

| | | | | | |
|---|---|---|---|---|---|
| Ramachandran plot | | | | | |
| Favored (%) | 96.47 | 94.93 | 95.0 | 95.36 | 93.77 |
| Allowed (%) | 3.45 | 5.07 | 5.0 | 4.57 | 6.16 |
| Disallowed (%) | 0.08 | 0 | 0 | 0 | 0 |

**Supplementary Table 2. Availability of data from variability refinements**

| Dataset name | DOI |
| --- | --- |
| Cdr1 apo | 10.5281/zenodo.16533987 |
| Cdr1 complexed with ITC, ATP an ADP-Mg | 10.5281/zenodo.16534106 |
| Transition from ATP-ADP-$VO_4$ to ATP-ADP<br>component C0 | 10.5281/zenodo.16530760 |
| Transition from ATP-ADP-$VO_4$ to ATP-ADP<br>component C1 | 10.5281/zenodo.16533789 |

### Supplementary movies

#### Supplementary movie 1

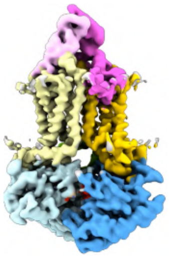

**Movie showing the conformation transition from frame 1 to frame 20 of *Cg*Cdr1 particle populations in component 0 from the closed [$^{s}$0 |$^{nc}$ATP|$^{c}$ADP-Vi-Mg] to the open [$^{s}$0 |$^{nc}$ATP|$^{c}$ADP-Mg] states.** The movie starts with the density maps and then the corresponding 3D-models, from the catalytic to the non-catalytic sides.

#### Supplementary movie 2

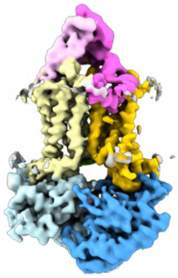

**Movie showing the conformation transition from frame 1 to frame 20 of *Cg*Cdr1 particle populations in component 1 from the closed [$^{s}$0 |$^{nc}$ATP|$^{c}$ADP-Vi-Mg] to the open [$^{s}$0 |$^{nc}$ATP|$^{c}$ADP-Mg] states.** The movie starts with the density maps and then the corresponding 3D-models, from the catalytic to the non-catalytic sides.

### Supplementary data

**Supplementary data 1:** Frames 2 to 20 *vs* frame 1 RMSD variations of the conformational variability analyses of *Cg*Cdr1 [$^{s}$0|$^{nc}$ATP|$^{c}$ADP-Vi-Mg$^{2+}$] to [$^{s}$0|$^{nc}$ATP|$^{c}$ADP-Mg$^{2+}$] states (Component C0)

**Supplementary data 2:** Frames 2 to 20 *vs* frame 1 RMSD variations of the conformational variability analyses of *Cg*Cdr1 [$^{s}$0|$^{nc}$ATP|$^{c}$ADP-Vi-Mg$^{2+}$] to [$^{s}$0|$^{nc}$ATP|$^{c}$ADP-Mg$^{2+}$] states (Component C1)
